# Topology, Energy Landscapes and Folding Kinetics of RNA Hairpins

**DOI:** 10.64898/2026.09.21.753325

**Authors:** Anjali Verma, Pratyush Tiwary, Yunrui Qiu

**Affiliations:** Institute for Physical Science and Technology, University of Maryland, College Park, MD 20742, USA; Biophysics Program, University of Maryland, College Park, MD 20742, USA; Institute for Health Computing, University of Maryland, Bethesda, MD 20852, USA; Department of Chemistry and Biochemistry, University of Maryland, College Park, MD 20742, USA; Department of Chemistry and Biochemistry, University of Notre Dame, Notre Dame, IN, USA

## Abstract

Many RNAs occupy multiple inter-converting structures in order to perform key biological functions. Characterizing RNA conformational ensembles is therefore critical to illuminating the mechanisms by which RNAs fold, unfold, and undergo precise structural rearrangements in response to cellular signals. However, resolving the equilibrium populations and folding kinetics of RNA ensembles is nontrivial due to the intrinsic ruggedness of RNA conformational landscapes and wide range of timescales spanned by RNA structural dynamics. To address these challenges, we present a *de novo* hierarchical multiscale computational framework that integrates diverse secondary and tertiary structure generation, adaptive unbiased molecular dynamics, a generative AI model termed latent thermodynamic flows (LaTF), and Markov state modeling to efficiently map global RNA folding landscapes and associated dynamics. Applied to the GCAA tetraloop, HIV-TAR stem-loop, and microROSE RNA thermometer, this framework yields *∼*1.3 ms of cumulative sampling and resolves temperature-dependent conformational landscapes comprising numerous native-like, misfolded, partially folded, and unfolded metastable states. Across all three RNA hairpins, temperature substantially reshapes landscape ruggedness, metastable state populations, and folding mechanisms. Our predicted ensembles show overall agreement with key static and dynamic biophysical observables, including Nuclear Magnetic Resonance (NMR)-resolved structures, Nuclear Overhauser Effect (NOE)-derived distance restraints, and residual dipolar couplings (RDCs). Together, these results establish a general physics–informed AI approach for accurately modeling RNA conformational heterogeneity, equilibrium thermodynamics, and long-timescale kinetics.

## I. INTRODUCTION

Ribonucleic acid (RNA) plays a central role in cellular biology, serving not only as an intermediary in genetic information transfer, but also as a versatile regulator of biological function. While known for its participation in many canonical cellular processes such as transcription and translation[1], enzymatic catalysis[2], and protein synthesis[3], RNA has also been implicated in multiple human diseases, including cancer, neurological disorders, and viral infections[4, 5]. To carry out diverse biological functions, RNAs form highly structured three-dimensional shapes designed to facilitate specific interactions with a wide range of biomolecules. Among the structural motifs adopted by RNA, hairpins, or stem-loops, are particularly widespread[6, 7]. These motifs typically comprise a Watson-Crick base-paired stem capped by a loop containing unpaired or noncanonically interacting nucleotides. By folding a single RNA strand back on itself, hairpins create compact local architectures that promote global RNA folding[8, 9], modulate RNA stability and degradation[10], and mediate RNA-protein recognition[11].

A wealth of computational and experimental evidence underscores the necessity of conceptualizing RNA as a dynamic ensemble rather than a single static structure[12–21]. Transitioning towards an ensemble-level description, i.e., characterizing RNA topological heterogeneity, energy landscapes, and associated thermodynamic and kinetic properties, is essential for elucidating functional mechanisms and enabling the rational design of RNA-targeted therapeutics[18–20, 22–24]. While recent advances in structural biology have facilitated the experimental and computational determination of static RNA structures for native states[25, 26], comprehensive mapping of dynamic RNA ensembles, including alternative metastable states and transient intermediates, remains in its early stages[15–17]. Moreover, the sensitivity of RNA dynamics to environmental factors such as temperature, pH, and ion concentrations makes the accurate construction of environmentally aware conformational ensembles particularly challenging[15, 17].

A diverse array of experimental methodologies, ranging from nuclear magnetic resonance (NMR) spectroscopy and cryo-electron microscopy (cryo-EM) to chemical probing, have been employed to study RNA structural dynamics. NMR is capable of probing dynamics across a broad temporal spectrum spanning picosecond-scale molecular tumbling, millisecond-scale base flipping, and minute-scale tertiary folding[27–32]. However, NMR is often constrained by limitations in spatial resolution[33] and cannot be used to study large RNA systems. Similarly, while cryo-EM can capture diverse static RNA structures of long-lived states, its application is typically hindered by coarse temporal (e.g., second-scale) and spatial (e.g., *∼* 4 − 7Å) resolution[34–37]. Biochemical approaches, such as chemical probing and proximity ligation, offer the advantage of high-throughput mapping of RNA base-pairing patterns in both *in vivo* and *in vitro* contexts, although these techniques remain limited to nucleotide-level spatial resolution and second-level temporal resolution[38–40].

In contrast to experimental techniques, computational approaches, particularly all-atom molecular dynamics (MD) simulations, have the potential to circumvent the aforementioned limitations. In principle, MD offers a high-fidelity description of structural dynamics for any RNA and can report on conformational changes with atomistic spatial detail and femtosecond temporal resolution. However, the modeling of RNA conformational ensembles via MD simulations has progressed only modestly over the past decade, constrained by challenges in force field (ff) accuracy, sampling efficiency, and data analysis. Although iterative developments in the AMBER and CHARMM families have improved classical RNA ffs, they have yet to reach the benchmark of reliability established for protein ffs[41–49]. This gap arises from multiple sources including: (i) the intrinsic complexity of RNA interactions, which are difficult to describe simultaneously with high fidelity[41–44]; (ii) the relative scarcity of experimental data, such as NMR observables or folding free energies, which limits the scope of ff calibration[46–49]; and (iii) the rugged nature of RNA energy landscapes, which hinders sampling convergence and complicates ff benchmarking[41, 44, 45]. As such, while current ffs can be used to qualitatively map RNA energy landscapes and identify metastable states, quantitative reports of thermodynamic and kinetic properties should be approached with caution.

Even with accurate ffs, RNA simulations are hindered by rugged folding landscapes and heterogeneous dynamics occurring across multiple orders of magnitude[50, 51]. Full exploration of RNA conformational landscapes by all-atom, unbiased MD is computationally prohibitive as it requires sampling transitions between different base-pairing geometries (*∼ µs*), ion-binding patterns (*∼ ms*), and distinct backbone folds (*∼ s*)[41, 52]. While enhanced sampling methods, such as metadynamics and generalized ensemble simulations effectively accelerate inter-basin transitions, they distort transition pathways and estimates of kinetic quantities[53–57]. Many of these techniques also rely on prior knowledge of optimal collective variables (CVs), which are more difficult to identify for flexible RNA systems than for proteins[41, 52]. Even if a well-sampled conformational ensemble is obtained, the structural heterogeneity exhibited by many RNAs still complicates downstream data analysis and mechanistic interpretation. For example, detailed structural descriptors are necessary to characterize RNA’s diverse interaction patterns[58, 59], and principled dimensionality reduction approaches are similarly required to resolve the non-degenerate manifolds separating biologically relevant metastable states[41, 60–64].

Beyond MD simulations, alternative ‘top-down’ computational frameworks provide a complementary paradigm for *in silico* RNA structure modeling by directly decoding secondary or tertiary structures from sequence. Classical secondary structure predictors typically employ dynamic programming combined with experimentally parameterized free-energy models (e.g., ‘Turner rules’) to identify the most stable and suboptimal conformations[65–72]. Tertiary structure predictors are mostly based on fragment assembly, in which local structural motifs from template libraries are assembled into full-length models, followed by a relaxation process which enables moderate sampling of structural heterogeneity[73–76].

Recently, advances in artificial intelligence (AI) have reshaped the field of structural biology, enabling *de novo* prediction of protein and RNA structures at an unprecedented scale. For RNA, machine learning (ML) scoring functions and neural network-based predictors have become increasingly popular for secondary structure prediction[77–79]. At the tertiary structure level, however, ML-based approaches must operate in a data-scarce regime due to the limited number of solved RNA structures. Current AI-based methods for 3D RNA structure prediction include covariation-based methods (e.g., AlphaFold3[26] and Boltz2[80]) or self-supervised large language models trained on massive sequence datasets to enhance transferability[81–85]. Nevertheless, when compared with AI-based protein structure prediction models, the corresponding RNA models still show limited predictive accuracy and are incapable of capturing RNA conformational diversity. Furthermore, the extraction of kinetic information from these top-down approaches remains limited, if not infeasible[86].

In this work, we address the challenges of computationally mapping RNA folding landscapes by building a hierarchical multiscale framework for *de novo* modeling of RNA equilibrium structural ensembles and their underlying kinetics. We adopt a recently developed classical RNA ff and focus on RNA hairpin systems for which this ff has been systematically tested[44, 87, 88]. Our framework integrates three complementary components: (i) secondary and tertiary structure prediction methods to efficiently assemble diverse hairpin topologies across distinct free-energy basins; (ii) generative-AI-accelerated adaptive all-atom MD simulations to enhance sampling of conformational transitions; and (iii) Markov state models (MSMs)[89–92] to construct a state-based representation of the RNA folding landscape and associated kinetics.

Using this workflow, we perform hundreds of microseconds of all-atom, explicit-solvent simulations to systematically characterize three hairpins at two different temperatures: the GCAA tetraloop (10 nt; cumulative simulation time, *∼*470 *µ*s), the HIV-TAR stem-loop (29 nt; *∼*480 *µ*s), and the MicroROSE RNA thermometer (29 nt; ∼385 *µ*s). The resulting ensembles and kinetic models show quantitative and qualitative agreement with multiple experimental observables, contextualizing previous results within a full conformational ensemble and providing rich atomistic detail. This comprehensive sampling approach enables a bottom-up, microscopic analysis of RNA hairpin thermodynamics and kinetics, revealing highly polymorphic conformational landscapes, temperature-modulated ensembles, and kinetic heterogeneity that approaches glassy non-ergodicity, which we term ‘glacial’. The resolved folding mechanisms, pathways, and intermediate or ‘excited’ states provide a mechanistic foundation for understanding RNA hairpin functions and biomolecular interactions, while advancing rational, structure-based RNA-targeted drug discovery. Finally, while ff optimization is beyond the scope of the current pipeline, we believe this work also opens a pathway for interested RNA ff researchers to test and improve their models.

## II. RESULTS

### A. A Hierarchical Multiscale Framework for *De Novo* Characterization of RNA Structural Dynamics

RNA energy landscapes are frustrated by many competing interactions, including but not limited to base pairing, base stacking, and solvent- or ion-mediated interactions [93]. This frustration, combined with the local stability of RNA secondary structures, gives rise to many diverse, long-lived, non-native conformational states. Consequently, RNA folding often proceeds through ‘kinetic partitioning’, in which distinct molecular subpopulations follow different pathways over widely separated timescales[94, 95]. These heterogeneous dynamical processes frequently exceed the reach of conventional unbiased MD simulations, and as such, RNA conformational landscapes often appear non-ergodic when viewed at typical simulation timescales. Markov state modeling, which discretizes conformational space into longlived metastable states and models transitions between them as Markovian jumps, provides a possible strategy to overcome this limitation: rather than relying on prohibitively long simulations, MSMs can reconstruct long-timescale dynamics from the local transitions sampled by many short simulations executed in parallel[89–92]. Meanwhile, by coarse-graining the conformational space into metastable states, MSMs render high-dimensional dynamics more interpretable and illuminate underlying biological mechanisms.

While running short simulations in parallel reduces wall-clock time, the set of initial seed structures must be chosen judiciously to ensure both efficient sampling of metastable states and the reliability of downstream MSMs[96–98]. In this context, simulating RNAs is particularly challenging due to the mixing of competing dynamical modes across different timescales. To address this, we construct a multiscale protocol that combines hierarchical assembly of diverse structures with ML-accelerated adaptive sampling, thereby enabling parallel simulations to sufficiently explore the global conformational landscape (see Fig. 1). Specifically, before initiating MD simulations, we combine secondary and tertiary structure prediction methods to survey global and local structural heterogeneity. At the secondary structure level, an array of predictors, spanning thermodynamics-based methods such as ViennaRNA[70], RNAstructure[69], EternaFold[71], and LinearFold[72], and machine-learning-based methods such as MXfold2[78], is used to enumerate competing secondary structure topologies (Fig. 1(a)). For each candidate topology, Fragment Assembly of RNA with Full-Atom Refinement (FARFAR2)[76] is used to construct a set of all-atom tertiary models from experimentally resolved local structural motifs, followed by local sampling and refinement of the candidate structures (Fig. 1(b)). Despite neglecting environmental factors, such as solvent and ion effects, and inheriting inaccuracies from nearest-neighbor models, this workflow provides an efficient route for rapidly generating putative RNA structural ensembles. Once generation is complete, the resulting ensembles are separated into distinct classes from which representative seed structures are selected for the first round of parallel explicit-solvent MD simulations (Fig. 1(c)). An overview of the different prediction methods, together with implementation details and results, is provided in the Methods section and Supporting Information (SI).

**FIG. 1.**
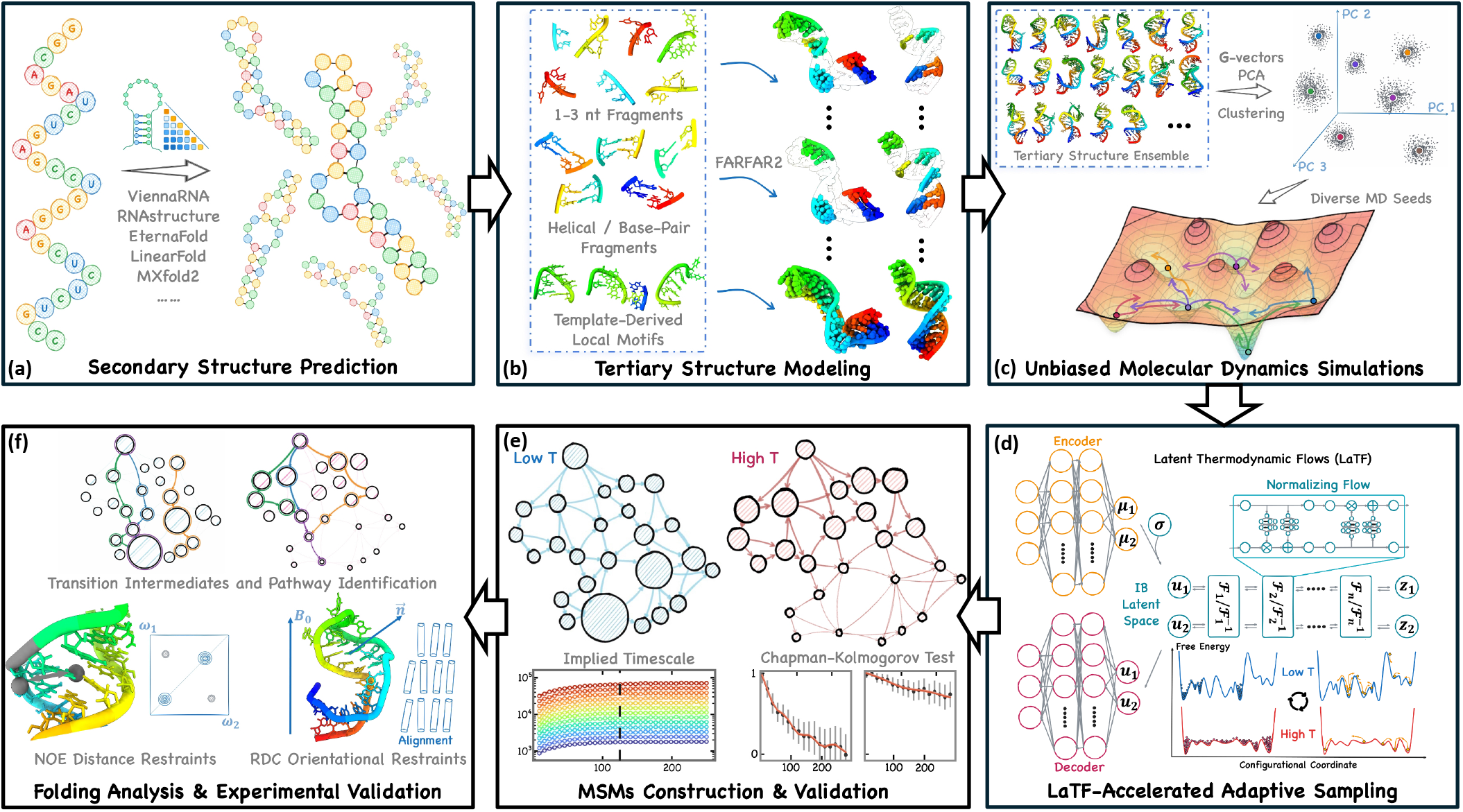
Schematic overview of the multiscale computational workflow for *de novo* modeling of RNA structural dynamics. (a) Multiple secondary structure predictors are applied to map the RNA sequence onto optimal and suboptimal secondary structures. (b) For each predicted secondary structure, an ensemble of tertiary models is assembled from structural fragments and refined using FARFAR2 scoring functions. (c) The resulted structural ensemble is partitioned into clusters. Representative cluster centroids are selected as initial seeds for unbiased MD simulations, enabling parallel sampling of local conformational transitions across the energy landscape. Independent simulations are conducted under low- and high-temperature conditions, respectively. (d) The resulting trajectories are used to train the LaTF generative model, which predicts temperature-dependent equilibrium distributions in a kinetically meaningful latent space and guides reseeding for the next round of simulations. This procedure is iterated over several rounds. (e) For each temperature, MSMs are constructed from the metastable states identified by the LaTF model and validated using implied timescale analysis and CK tests. (f) MSMs are further used to characterize transition intermediates and folding pathways, while NMR experimental observables provide additional validation of the computational results.

To further improve sampling efficiency and convergence, we perform multiple rounds of adaptive sampling guided by latent thermodynamic flows (LaTF)[99], a unified representation-learning and generative-modeling framework (Fig. 1(d)). LaTF integrates three components: a Gaussian encoder that maps high-dimensional structural descriptors of each conformation into a 2D latent information bottleneck (IB) space; a normalizing flow that transforms the encoded distribution into a prior; and a decoder that predicts the metastable state reached after a specified lag time. The LaTF encoder-decoder learns the structural features most relevant to slow, biologically significant dynamics and groups conformations by kinetic similarity into metastable states, while the flow model enables both generation and explicit likelihood evaluation in the latent space. By introducing a temperature-steerable tilted Gaussian prior, LaTF has been shown to reliably infer temperature-dependent latent equilibrium distributions from limited data[99]. For each RNA hairpin studied here, we consistently perform simulations under paired low-and high-temperature conditions. After each round of simulation, a LaTF model is trained to integrate information from both temperatures, thus allowing high-temperature data to augment inference of the low-temperature landscape. This mixing of information across temperature guides exploration of under-sampled regions and is especially crucial during the early stages of sampling. Once the LaTF model is trained, free-energy landscapes are generated at both temperatures to select new starting conformations for next round of simulations.

After multiple rounds of sampling are completed, a final LaTF model is trained exclusively on the low-temperature data to identify the latent IB space and metastable states of interest without bias from high-temperature sampling. Key training hyperparameters are selected through cross-validation using the generalized matrix Rayleigh quotient (GMRQ) score, ensuring that the LaTF model captures the dominant slow dynamical processes[100, 101]. The high-temperature simulation data is then passed through the trained LaTF encoder and decoder, enabling dynamical modeling under a shared state decomposition. Although ensembles can be generated at any temperature using the LaTF tilted prior, we choose not to perform interpolation or melting curve estimation due to a lack of suitable benchmarks and consider it beyond the scope of the current research. Instead, we analyze all simulation data using the state definitions learned at low-temperature to directly assess how temperature impacts RNA ensemble thermodynamics and kinetics. To quantify these effects, MSMs are constructed independently for low and high temperature and validated using implied timescale analysis and Chapman-Kolmogorov tests[89] (Fig. 1(e)). Additional details on the LaTF model, adaptive sampling protocol, model training, and MSM validation are presented in the Methods section and SI.

The resulting MSMs quantify equilibrium populations and transition dynamics among native and ‘excited’ states, permitting direct comparison with NMR observables such as nuclear Overhauser effect (NOE) distance restraints and residual dipolar couplings (RDCs)[102]. They also facilitate identification of folding pathways and intermediate states, as well as systematic investigations of how these pathways are reshaped by temperature (Fig. 1(f)). Finally, using the constructed energy landscapes as benchmarks, we evaluate tertiary structure prediction approaches beyond FARFAR2[76], including 3dRNA[74, 103], AlphaFold3[26], and Boltz2[80], and find that thermodynamics-based methods yield better sampling of structural ensembles as compared to purely ML methods (see Figs. S10, S22, S32).

Recently, the Tiwary group has developed RNAnneal[16], a complementary framework that integrates secondary and tertiary structure modeling with short, implicit-solvent MD and physics-informed ML to provide an efficient, but approximate, estimation of thermodynamic weights across diverse folded structures. In contrast to RNAnneal, the present pipeline incorporates adaptive sampling, extensive explicit-solvent simulations, and MSM-based kinetic modeling, enabling a far more detailed and comprehensive characterization of global RNA folding landscapes and their associated kinetics.

### B. GCAA Tetraloop

Comprised of four single-stranded nucleotides that connect the termini of a helical stem, the tetraloop (TL) is among the most prevalent loop motifs in RNA. Despite their small size, TLs exhibit diverse interaction patterns, including canonical and noncanonical base pairing and base–phosphate interactions. Furthermore, as part of larger RNAs, TLs can also undergo conformational remodeling to form long-range contacts necessary to stabilize tertiary structure [105, 106]. While extensive experimental and computational efforts have been devoted to investigating TL conformational dynamics[44, 88, 107–117], the folding mechanisms and kinetics of many TLs remain only partially understood due to their exceptional thermodynamic stability and intricate internal interaction networks. Here, we present a detailed characterization of the structural heterogeneity and folding dynamics of the GCAA tetraloop, an evolutionarily conserved motif identified through phylogenetic analyses.

#### Folding Energy Landscapes

Using our multiscale sampling pipeline, we collect 80 trajectories for the GCAA TL at both 300K and 400K, totaling ∼ 322 and ∼ 148 *µs*, respectively (Fig. S1(a)). From our adaptive unbiased simulations, we calculate a free energy surface (FES) along the heavy-atom root-mean-square deviation (RMSD) and folding compactness order parameter *Q* (see definitions in SI Sec. IV). The FES, depicted in Fig. S3, is consistent with a prior reference obtained from parallel tempering combined with biased well-tempered metadynamics under the same ff[88]. The agreement with existing simulation data obtained via enhanced sampling, as well as the robust interconnection between folded, misfolded and unfolded states, demonstrates the effectiveness of our adaptive sampling strategy.

Further training of the LaTF model on the low-temperature data with a lag time of 100 ns identifies 20 long-lived metastable states, where states with shorter lifetimes are merged into longer-lived ones (Fig. S2(a)). This rich metastability contrasts sharply with typical protein folding systems. As shown in Fig. 2(a), the resulting 300K folding landscape in the LaTF IB latent space is highly multi-funneled, with numerous kinetic traps corresponding to competing misfolded intermediates. Upon increasing the temperature to 400K, the landscape becomes substantially smoother: the native folded state remains accessible, whereas most misfolded states are depleted.

**FIG. 2.**
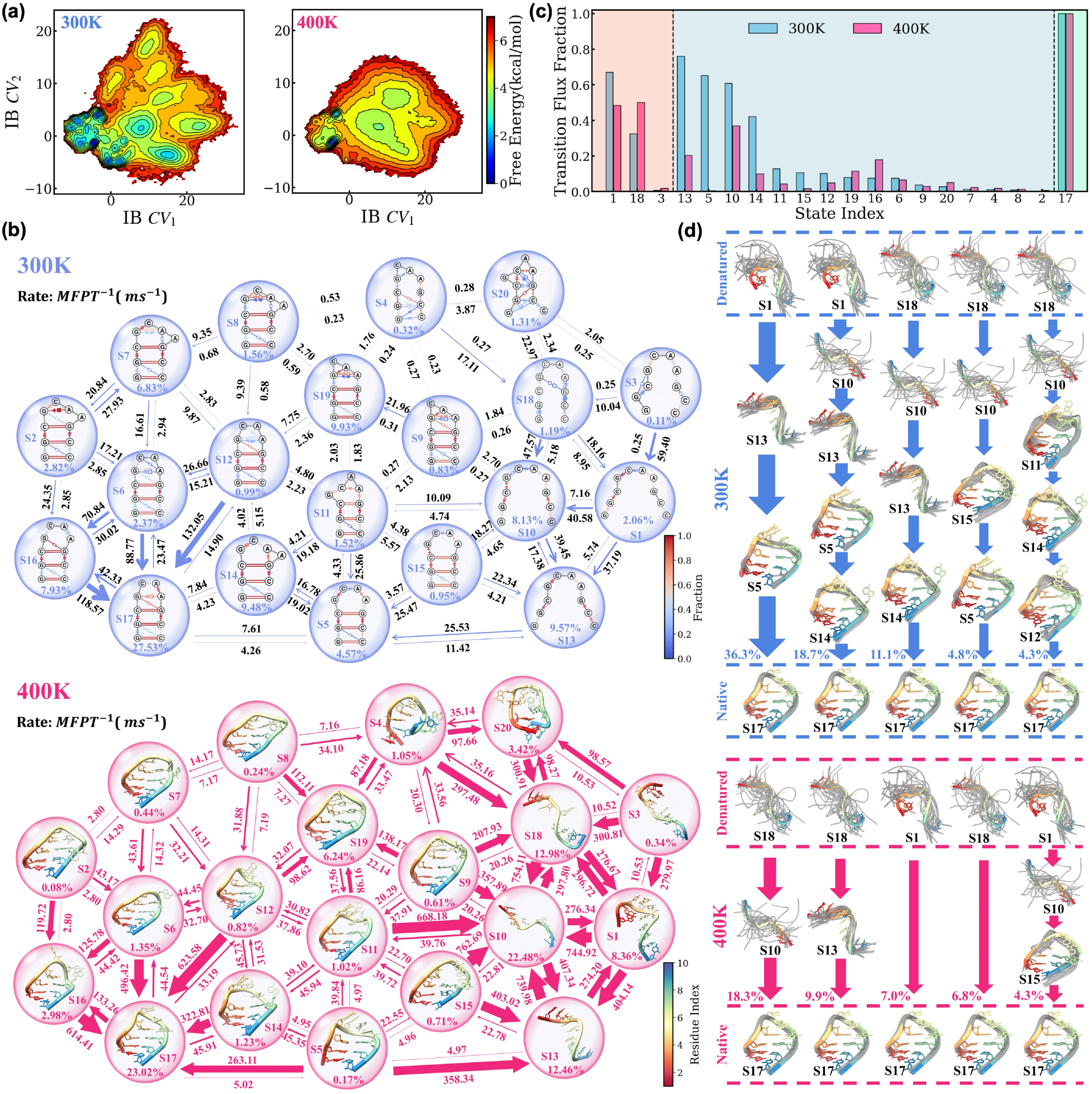
Folding free-energy landscape and kinetics of the GCAA tetraloop at 300K and 400 K. (a) Free-energy surfaces projected onto the IB space at the two temperatures. Free energies are estimated by reweighting the raw simulation data using the state stationary populations obtained from the corresponding MSMs. (b) Folding kinetic networks at the two temperatures, where transition arrow widths are proportional to the rates (inverse MFPTs) and state populations are indicated. For the 300K network, dynamic secondary structure representations are constructed from 2000 conformations randomly sampled from each metastable state, where structural annotations follow the Leontis-Westhof classification[104], and the color scheme denotes the fraction of conformations exhibiting each interaction pattern. For the 400 K network, the dominant tertiary structure of each state, ranked highest by the LaTF model, is shown. (c) Cumulative folding transition-pathway fluxes through each metastable state at the two temperatures. Folding pathways and their associated fluxes are identified and quantified using transition path theory in conjunction with the MSMs. Source unfolded states (orange), intermediate states (blue), and sink folded states (green) are categorized and shown accordingly. (d) The five most prominent folding pathways at the two temperatures, together with their associated fluxes. Each state is represented by its dominant tertiary structure (colored), together with 30 additional randomly selected conformations (transparent).

By incorporating metastable state prediction into the training scheme of LaTF, which approximates low-dimensional reaction coordinates on-the-fly, a large number of metastable states can be accommodated within a latent space of only 2D. In contrast, when the FES constructed at 300K is projected onto conventional order parameters, such as the radius of gyration, stem and loop RMSD, and *ɛ*RMSD relative to the NMR structure, many of the long-lived states identified by LaTF are not distinguished (Fig. S4). Although unable to resolve the diversity of conformational states seen in the LaTF IB space, the conventional order parameters do reveal local structural variations underlying global conformational dynamics. Notably, the loop exhibits greater flexibility and undergoes pronounced rearrangements even at low global RMSD (Fig. S4(b)), whereas the stem RMSD is closely correlated with the overall structural deviation (Fig. S4(a)), suggesting partially decoupled folding dynamics between the stem and loop regions.

#### Metastable States & Transition Kinetics

The constructed MSMs provide state-based interpretations of the GCAA folding landscape and dynamics (Fig. 2(b)). Inspection of the metastable states, together with visualization of their dynamic secondary structures and representative tertiary structures (ranked by the normalizing-flow likelihood), elucidates intra- and inter-state structural heterogeneity, particularly within the loop region. In addition to the most populated native state (S17), which faithfully recapitulates the stable interactions observed in NMR structures (PDB ID: 1ZIH)[118], twelve non-native misfolded states are identified that retain a folded stem but exhibit alternative loop conformations. The noncanonical *G_L_*_1_–*A_L_*_4_ interaction, which undergoes substantial dynamic breathing, provides an informative descriptor for distinguishing these states. When this contact is formed, the flexible *A_L_*_3_ can either become solvent-exposed (S6 and S12), or form an alternative *G_L_*_1_–*A_L_*_3_ stacking interaction (S9 and S19). When the *G_L_*_1_–*A_L_*_4_ contact is disrupted, *G_L_*_1_ can either form additional stacking interactions with the last six nucleotides (S5, S11, S14, and S15), mispair with *A_L_*_3_ (S2, S7, and S8), or leave the other three loop nucleotides solvent-exposed (S16). Apart from these stem-folded states, we also detect collapsed states (S4 and S20), in which stem nucleotides are stacked into compact structures without canonical base-pairing interactions, as well as unfolded states (S1, S3, S10, S13, and S18) distinguished by distinct partially stacked segments.

At 300K, multiple non-native misfolded states maintain appreciable populations, reflecting competition among alternative noncanonical base-pairing and stacking interactions. These interactions make important energetic contributions to TL stability, and their disruption and reformation occur on timescales as slow as milliseconds, with some persisting even longer than stem-folding transitions, as quantified by mean first-passage times (MFPTs; Fig. 2(b)). The GCAA TL shows distinct dynamical behavior across temperatures, with the lower-temperature regime characterized by highly heterogeneous and slow dynamics. At higher temperature, misfolded states become less stable and transitions accelerate, leading to a clearer separation of slow and fast processes. This reflects strongly heterogeneous dynamics consistent with a ‘glacial’ landscape picture. In the SI, we provide detailed quantitative analysis of TL dynamics via the MSM transition probability matrix (TPM; Fig. S6).

#### Folding Pathways

Transition path theory (TPT)[119, 120] is applied to the constructed MSMs to characterize folding pathways and corresponding reactive fluxes, with the most disordered unfolded states (S1, S3, and S18) defined as sources and the native state S17 as the sink (Fig. 2(c-d))). At 300K, folding follows highly heterogeneous routes that traverse multiple intermediates. The dominant pathways reveal a hierarchical folding mechanism: disordered unfolded conformations first organize into sequentially stacked unfolded structures (e.g., S10 and S13), the stem then rapidly zips through in-register intermediates, and noncanonical loop interactions subsequently emerge through a slower search among various intermediates. Misfolded states featuring *G_L_*_1_ stacking with the last six nucleotides act as major flux-carrying intermediates (e.g., S5 and S14), likely because they allow stem nucleation from stacked unfolded conformations without rearrangement of existing stacking interactions. Similar states were also preferentially sampled in the enhanced sampling study of GCAA TL folding done by Debenedetti and colleagues[88]. Further examination of transition-region structures connecting unfolded states to transient misfolded states (defined by LaTF state probabilities) reveals that stem folding can initiate through either terminal or central nucleotide-pair proximity (Fig. S7), consistent with prior experimental and computational studies[112, 121, 122]. In contrast, collapsed states (S4 and S20) and *G_L_*_1_–*A_L_*_3_ mispaired states (S2, S7, and S8) behave as off-pathway kinetic traps, carrying negligible reactive flux. Direct one-step folding pathways are detected but account for only a minor ∼ 1% of the total flux. At 400 K, the more funneled landscape significantly promotes the direct nucleation–collapse mechanism, with native stem and loop interactions forming rapidly and synchronously. Intermediates with solvent-exposed loop nucleotides become prominent (e.g., S16 and S19), potentially because of their entropic stabilization. Overall, pathway analysis shows that temperature critically modulates the folding mechanisms.

#### Experimental Validations & Other Computational Results

At the global conformational level, Xia *et al.* have reported alternative GCAA loop conformations detected using ultrafast spectroscopic measurements [123]. In one such alternative conformation, *C_L_*_2_ is flipped out while *A_L_*_3_ is partially stacked on *G_L_*_1_, which pairs with *A_L_*_4_. Upon examination of our MSM conformational states, we find that these interactions occur in the well-populated non-native state S19. Another experimentally observed alternative conformation shows *A_L_*_3_ flipped out and *C_L_*_2_ stacked on *G_L_*_1_, which is consistent with our MSM state S6. At the nucleotide level, prior NMR[118] and computational studies[111, 113, 117] have shown that *G_L_*_1_ is relatively immobile, *C_L_*_2_ is the most mobile loop nucleotide, and *A_L_*_3_ and *A_L_*_4_ are less mobile than *C_L_*_2_. Using our 300K MSM, we quantify local base-pairing and stacking interaction networks around each tetraloop nucleotide in a base-centered coordinate system, with distributions reweighted by MSM state populations (see results in Fig. S8). The results are consistent with prior findings: *C_L_*_2_ exhibits the lowest pairing frequency, whereas the other three loop nucleotides form more frequent noncanonical pairing interactions. At the atomistic level, a set of semi-quantitative short-range interproton distance restraints has been obtained from NOE spectroscopy and used in the determination of the NMR structures (PDB ID: 1ZIH)[118]. While differences between the simulation and NMR conditions may alter the relative populations of individual states, we restrict the comparison to structures sampled from the dominant S17 and evaluate this folded sub-ensemble against the NOE restraints, finding good agreement between the two (Fig. S9). A detailed analysis of all experimental validation is provided in the SI. Collectively, our findings support the effectiveness of our *de novo* computational framework: the ff favors native state stability, the hierarchical sampling strategy recovers both the native structure and its local conformational variability, and LaTF reliably identifies and prioritizes the native state.

Over the past decade, the GCAA tetraloop has been extensively studied using diverse force fields, solvent models, and sampling strategies, including unbiased MD, replica-exchange methods, and metadynamics[44, 88, 107–117]. A summary of previous simulation setups and key findings is provided in Supplementary Table S1. Simulations initiated from NMR structures have primarily characterized fluctuations near the native basin[111, 117], whereas studies of global folding remain limited by sampling: distributed unbiased MD provides uneven coverage of metastable transitions[116], while enhanced sampling methods improve exploration at the cost of unbiased kinetics and detailed pathway resolution[88, 112–115]. Here, to our knowledge, we report the longest explicit-solvent unbiased simulations of the GCAA tetraloop at two temperatures, enabling systematic characterization of its structural ensemble and folding dynamics. Our results extend and refine several observations from previous computational studies. While multiple non-native folded states had been reported in literature, their structural diversity and relative populations remained incompletely characterized[88, 108, 111, 113, 115]. Consistent with prior work, we observe that many non-native states retain a largely native-like stem but differ primarily in loop organization, occupy comparable free-energy basins, and serve as on-pathway intermediates or kinetically isolated off-pathway states[88, 108, 112]. The slower relaxation of the loop relative to stem formation is also in line with previous observations[108, 114].

Moreover, to examine whether efficient tertiary structure prediction approaches can reproduce the full conformational diversity revealed by our simulations, we project ensembles generated by different approaches including FARFAR2[76], 3dRNA[74, 103], AlphaFold3[26], and Boltz2[80] onto the LaTF latent space. Despite their substantially lower computational cost, these methods provide incomplete coverage of the equilibrium landscape (Fig. S10): thermodynamics-based modeling approaches only sample portions of the non-native folded ensemble but do not reach unfolded regions, whereas the AI models are strongly concentrated around native-like conformations. Efficient structure generation alone does not recover the structural heterogeneity of RNA, thus underscoring the need for adaptive molecular simulations in addition to hierarchical structure generation.

### C. HIV TAR Stem-loop

The trans-activation response element (TAR) is a 59-nt stem–loop located at the 5*^′^* end of HIV-1 transcripts. Its interactions with the Tat protein and the super elongation complex are essential for HIV transcription regulation, stimulating long-terminal repeat promoter activity and enabling transcription elongation[124]. TAR consists of two key structural motifs: the trinucleotide UCU bulge and hexanucleotide apical loop, both of which contribute to Tat recognition[125, 126]. Experimental and computational studies have shown that TAR samples substantial conformational heterogeneity and populates different “excited” states. However, understanding the interplay between these distinct conformational states and how they modulate Tat recognition remains an important mechanistic question[29–32, 127–130]. Since disruption of Tat recognition could suppress HIV transcription, HIV-TAR has long served as a promising RNA target for structure-based discovery of small molecules and peptides[24, 131]. Here, we focus on a 29-nt HIV-TAR stem construct that preserves the bulge and apical-loop region central to Tat recognition.

#### Folding Energy Landscapes

The simulated HIV-TAR stem-loop system contains ∼ 37, 000 atoms. Our computational protocol generates 400 trajectories at 310K and 80 trajectories at 420K, with aggregate simulation times of ∼ 394*µs* and ∼ 87*µs*, respectively (Fig. S1(b)). Using the low-temperature data, a LaTF model is trained at a lag time of 200 ns and resolves 40 long-lived metastable states. The resulting 310K FES (Fig. 3(a)) is markedly more polymorphic than that of the tetraloop, with multiple metastable sub-basins that reflect more complex and diverse conformational rearrangements. In contrast to the 310K landscape, the 420K landscape is far more funneled and shifts toward a broad unfolded basin, thus illustrating the role of temperature in determining landscape topology and complexity.

**FIG. 3.**
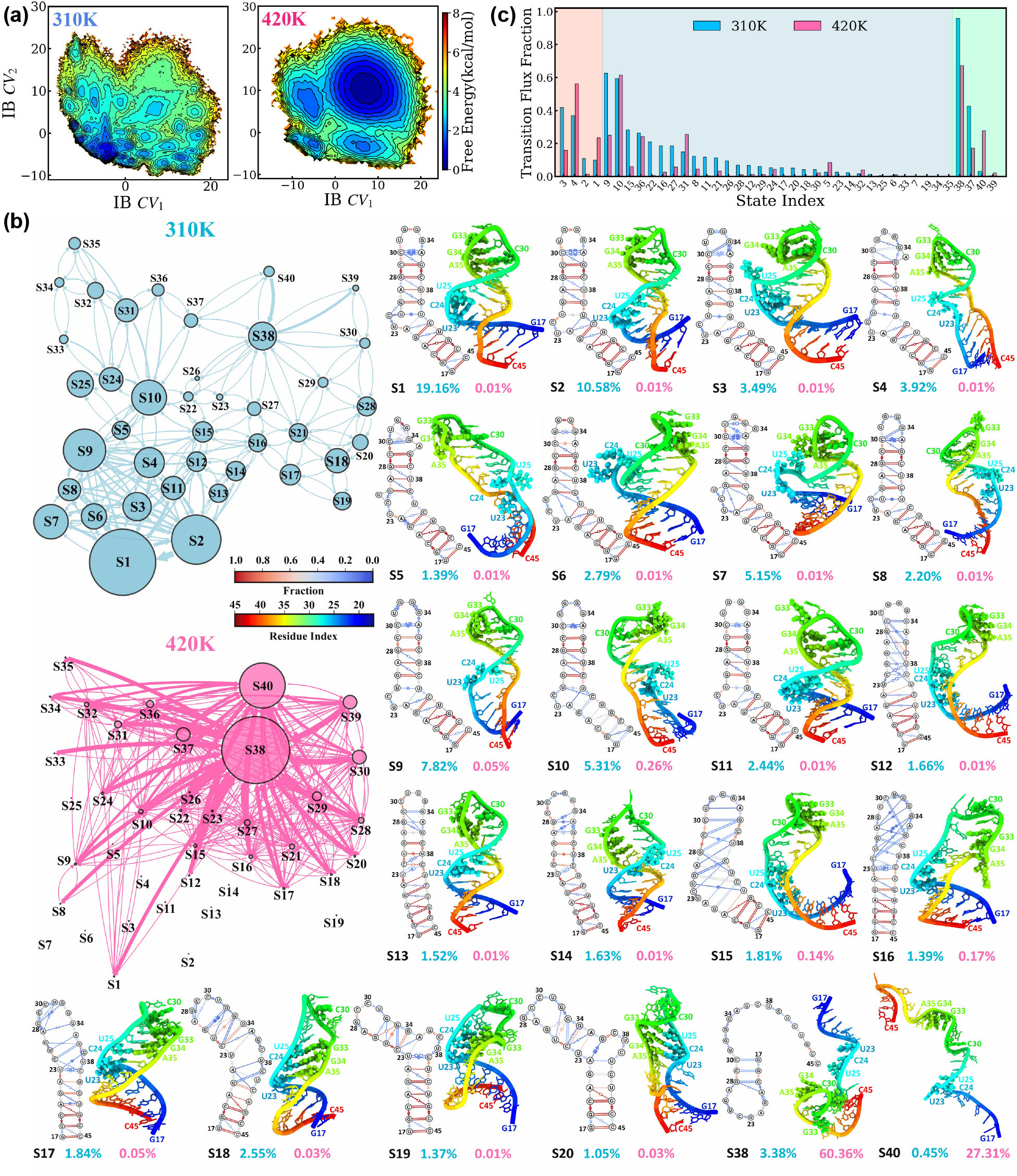
Folding free-energy landscape and kinetics of the HIV-TAR stem-loop at 310K and 420K. (a) Free-energy surfaces projected onto the LaTF IB space at the two temperatures. Free energies are estimated by reweighting the raw simulation data using the state stationary populations obtained from the corresponding MSMs. (b) Folding kinetic networks at the two temperatures, with transition arrow widths proportional to the corresponding rates (inverse MFPTs) and state circle areas proportional to their equilibrium populations. The dynamic secondary structures of the twenty-two states are shown based on 500 conformations randomly selected from each metastable state, with structural interactions annotated according to the Leontis–Westhof classification[104]. Colors indicate the fraction of conformations exhibiting each interaction pattern. The dominant tertiary structure of each state, corresponding to the conformation ranked highest by the LaTF model, is shown alongside. The apical loop and bulge regions are highlighted in sphere representation. Nucleotide numbering begins at 17 and extends to 45. State populations at the two temperatures are reported using the same color scheme as corresponding networks. (c) Cumulative folding transition-pathway fluxes through each metastable state at the two temperatures. Folding pathways and their associated fluxes are identified and quantified using TPT in conjunction with the MSMs. Source unfolded states (orange), intermediate states (blue), and sink folded states (green) are categorized and shown accordingly.

LaTF resolves numerous conformational states within a 2D latent space constructed from thousands of structural descriptors, whereas RMSD-based coordinates provide a more intuitive, albeit highly degenerate, representation of underlying structural transitions. We further analyze the RMSD distributions of the lower stem (G17–A22 and U40–C45; nucleotide numbering starts at 17), upper stem (G26–C29 and G36–C39), bulge loop, and apical loop (C30–A35) relative to the NMR structural model across all simulated conformations (Fig. S11). The resulting FESs indicate that the global conformational dynamics of HIV-TAR are dominated by rearrangements of the two helical stems, with substantially weaker coupling to motions of the bulge and apical loops. Joint analysis of the upper- and lower-stem RMSDs further supports a sequential folding mechanism in which the two stems can form in either order.

#### Metastable States & Transition Kinetics

Analysis of the transition network and associated dynamic secondary and tertiary structures reveals substantial conformational heterogeneity across the MSM states (see 3(b) and Fig. S14). The experimental native topology features two helical segments joined by a trinucleotide bulge, a global architecture preserved across states S1 to S11 in the lower-left region of the network. These states are distinguished mainly by variations in base-pairing and stacking interactions within the hexanucleotide apical loop and the UCU bulge. The highest populated states, S1 and S2, exhibit stabilized sequential stacking interactions in the apical region. However, S1 features a more flexible bulge wherein U24/U25 are frequently extrahelical, while S2 favors intrahelical bulge stacking. Some other states display a dynamic C30-G34 Watson-Crick interaction in the apical loop, accompanied by either intrahelical (e.g., S3 and S8) or extrahelical (e.g., S4 and S5) conformations of A35. In these states, the bulge remains highly flexible, with multiple nucleotides exposed to solvent or forming transient stacking interactions with residues in the upper stem. The apical loop can also adopt a more compact, zipped-up conformation stabilized by two consecutive noncanonical C30–A35 and U31–G34 interactions, as observed in S6, while all bulge nucleotides become extrahelical and solvent-exposed. Alternatively, while C30–A35 remains dynamically paired, G34 can exhibit increased flexibility, as observed in S7, where G34 is in close proximity to bulge nucleotide U25. The bulge itself can also adopt an ordered intrahelical conformation with sequential stacking while the apical-loop nucleotides become disordered and solvent-exposed, as in S11. Moreover, both the bulge and apical loop can remain simultaneously dynamic (e.g., S9 and S10), with these fluctuations occasionally extending to terminal base-pair breathing in the lower stem.

Additionally, with a stable lower stem, the upper strand can undergo a register shift that reorganizes the bulge, upper stem, and apical loop, replacing canonical Watson–Crick pairing in the upper stem with a network of U–U wobble pairs and G–A mismatches and introducing transient pairing or stacking interactions of U23 and C24 with C39 (e.g., S13 and S14). States bridging the native and shifted upper-stem registers are also identified, with highly dynamic and disordered interactions throughout the upper stem (S12, S15, and S16). More extensive register rearrangements give rise to a distinct set of metastable states featuring an alternative bulge spanning G36–C39 (e.g., S17–S21). Register shifting in the opposite direction is also accessible, producing an alternative bulge centered around A22 (e.g., S22 and S23), although these states remain only sparsely populated. A separate set of metastable conformations retains the native upper-stem register while forming a bulge around A22 accompanied by extrusion of G21 (e.g., S24 and S25).

As the RNA transitions toward less compact and unfolded conformations, structural disorder can originate from either the upper or lower stem. Upon disruption of the lower stem, the C30–G34 Watson–Crick interaction persists across a number of states (e.g., S31–S35), while the released strands can reorganize into short local hairpins, as seen in S26, S36, S38, and S39. Further loss of these residual interactions ultimately leads to fully unfolded conformations (e.g., S40).

Across the diverse metastable states, constructed MSM transition networks reveal pronounced kinetic heterogeneity at 310K (Fig. 3(b)). Interconversions among states with intact upper and lower stems but distinct apical-loop and bulge conformations are relatively rapid, whereas transitions toward register-shifted states are slower, and exchange among unfolded, misfolded, and partially folded states is slower still. These heterogeneous dynamics are further reflected in the TPM eigenvalue spectrum, which shows no clear separation of timescales (Fig. S13(b)). Increasing temperature reshapes the folding landscape and kinetic network by favoring extended, unfolded conformations and significantly accelerating transitions between unfolded and partially folded states, resulting in a more pronounced separation in the eigenvalue spectrum (Fig. S13(e)).

#### Folding Pathways

Taking the most extended unfolded states S37–S40 and the dominant folded states S1–S4 as the source and sink sub-ensembles, respectively, TPT[119, 120] is applied to the LaTF-derived MSMs to resolve the kinetic folding pathways of HIV-TAR at two temperatures and characterize their associated reactive fluxes. The dominant flux-carrying pathways and cumulative flux through each metastable state are illustrated in Figs. S15, S16, and 3(c). At 310K, TAR exhibits two major classes of sequential folding pathways, distinguished by the order of upper- and lower-stem formation. Folding can nucleate near the apical loop, producing upper-stem-first intermediates such as S31 and S36, followed by progressive helix propagation toward the lower stem. Alternatively, nucleation can initiate from the terminal region, as represented by S15, S16, S22, and S27, leading to lower-stem-first intermediates before formation of the upper helix. In both pathways, stem formation precedes organization of the flexible bulge and apical loop (e.g., S9 and S10), which subsequently relax. Our resolved pathways complement prior studies conducted under different physical conditions. Optical-tweezer experiments have previously suggested a nucleation-and-helix-propagation mechanism and implicated the three-nucleotide bulge in kinetic frustration[132, 133]. Upper-stem hairpin intermediates have also been identified by force-ramp measurements combined with secondary structure modeling[134], whereas force-free coarse-grained simulations have predicted a lower-stem-retaining intermediate during thermal unfolding[135].

Despite representing prominent alternatives near the dominant folded basin, the loop-zipped conformation (S6) and register-shifted conformations (S13 and S14) carry little folding flux and appear to function as off-pathway misfolded states. Direct folding pathways, in which the two stems form in rapid succession, are also detected. However, consistent with a kinetically partitioned folding mechanism, they contribute substantially less reactive flux at 310K. At 420K, the reactive flux becomes more concentrated along direct and upper-stem-first pathways, with S31 and S36 emerging as the predominant intermediates.

#### Experimental Validations & Other Computational Results

Over the past decades, NMR studies have determined the native HIV-TAR stem-loop structures (referred as ground state; GS), and identified two lower-populated excited states, ES1 and ES2, with distinct conformational rearrangements[29, 31]. Taken together, these data provide a useful benchmark for our computational ensemble. The NOE-refined GS structures (PDB ID: 1ANR)[136] exhibit local structural heterogeneity, and consequently map onto five LaTF metastable states (S3, S4, S8, S9, and S10). We therefore construct the folded-state ensemble from these states according to their renormalized MSM populations and calculate ensemble-averaged inter-proton distances. Of the 744 NOE-derived distance restraints, only 15 deviate from the experimental ranges by more than 1 Å(Fig. S17), with most violations localized near the bulge and none within the apical loop, indicating fair agreement with experiment.

ES1 is experimentally characterized by a zipped apical-loop conformation stabilized by the noncanonical C30–A35 and U31–G34 interactions[29], closely matching the S6 state identified in our ensemble. ES2 arises from a strand-register shift that replaces several canonical base pairs with mismatches, including U25–U38 and A27–G36[31], and corresponds closely to our states S13 and S14. Leveraging our sampled equilibrium ensemble, we further construct local coordinate systems for four nucleotide pairs to characterize their interaction patterns (Fig. S18(b)), which serve as key structural descriptors distinguishing the GS and ES conformations. The resulting free-energy landscapes (Fig. S18(c)) reveal multiple metastable interaction patterns. Al-Hashimi *et al.* [30–32] previously constructed structural ensembles for GS and ES1 (PDB IDs: 7JU1 and 8THV), as well as for ES2 using stabilizing mutations (PDB ID: 8U3M), by generating diverse conformational pools with FARFAR followed by selection and validation against NMR observables, including RDCs and chemical shifts (Fig. S18(a)). Projection of these experimentally derived sub-ensembles onto our free-energy landscapes shows good agreement: ES2 structures localize within the corresponding shallow excited-state basin, whereas the GS/ES1 ensemble spans several neighboring basins, reflecting substantial structural heterogeneity in the apical loop. Although the C30–G34 cross-loop interaction has been widely documented as a prominent GS feature, conformers lacking this contact are required to reproduce experimental observables. This necessary inclusion of heterogeneity is consistent with the conformational diversity captured by our simulation-derived landscape.

Beyond ES1 and ES2, TAR samples additional conformations with potential functional relevance. Prior experiments and simulations suggest transient proximity between the bulge and apical loop, two regions that jointly organize the Tat-recognition architecture[128, 137, 138]. Consistent with this picture, our equilibrium ensemble exhibits a clear population at short bulge–loop separations. The minimum distance between oxygen and nitrogen atoms in the two regions peaks near 3 Å, which is indicative of direct interactions that are not represented by the native NMR models (Fig. S19(a)). We also observe low-population conformations in our ensemble resembling the U23–A27–U38 base triple (Fig. S19(b)), a motif implicated in ligand- and Tat-recognition-competent states and whose disruption impairs binding and Tat-dependent transactivation[139–142].

Given that RDCs report the ensemble-averaged orientations of internuclear vectors relative to the external magnetic field, we further compare experimental RDCs with those calculated from the MSM-population-weighted simulation ensemble. To avoid overfitting, molecular alignment and RDC prediction are performed using a steric obstruction model rather than optimal singular-value-decomposition fitting; details of the PALES calculations[143] are provided in the SI. The simulated and experimental RDCs[32, 144] show moderate agreement, with a Pearson correlation of ∼ 0.63 (Fig. S20(a)). Discrepancies in the RDCs may arise from differences between the experimental liquid-crystalline environment and the aqueous simulation conditions, as well as limitations in force-field accuracy and population estimation. Nevertheless, we expect our extensively sampled atomistic ensemble to provide a broad structural basis for future experimental reweighting or ensemble refinement. We further calculate Lipari–Szabo generalized order parameters, *S*^2^[145], for C–H bond vectors across nucleotides and compare them with values derived from experimental spin-relaxation measurements[129, 146]. The resulting profiles show qualitative agreement with experiment, consistently identifying the bulge and apical loop as the most dynamically flexible regions (Fig. S20(b)). Details of the *S*^2^ calculations and MSM weighting are provided in the SI. A summary of representative prior experimental and simulation studies is provided in Table S2.

A broad range of peptides and small molecules have been developed to target distinct regions of the TAR stem loop, with several corresponding structural complexes resolved experimentally[147]. Mapping these ligand-bound TAR conformations onto the LaTF landscape places them across seven metastable states, with most ligand classes sampling more than one state. This structural diversity highlights the importance of adequately sampling the accessible TAR ensemble before pursuing structure-based ligand design. A summary of the reported binders, associated TAR conformations, and their metastable-state assignments is provided in Table S8 and Fig. S21. Several tertiary structure modeling approaches are also evaluated for TAR ensemble prediction. AlphaFold3[26] and Boltz2[80] predictions remain strongly confined near the NMR-like native basin, while FARFAR2[76] and 3dRNA[103] generate more diverse conformations, yet cover only part of the accessible structural ensemble. These limitations may arise from biases in training data and the sparse representation of relevant motifs in fragment- or template-based structural libraries.

### D. MicroROSE RNA Thermometer

RNA Thermometers (RNATs) are temperature sensing regulatory elements typically located in the 5’ untranslated region (UTR) of bacterial messenger RNAs (mRNAs). At low temperatures, the RNAT folds into a secondary structure which sequesters the ribosome binding site (RBS) and inhibits binding of the 30S ribosomal subunit. The RNAT gradually unzips with increasing temperature, thereby releasing the RBS and enabling translation initiation[148, 149]. By sensing temperature directly rather than using downstream signals such as protein aggregation, RNATs can rapidly and reversibly modulate translation efficiency in response to even 1°C temperature changes[150, 151].

As our third and final system, we examine the ROSE_1_ (Repression Of heat Shock gene Expression) thermometer of *Bradyrhizobium japonicum*[152]. ROSE elements include a conserved U-U/C-G-C-U sequence motif, where the central G nucleotide is critical for temperature sensing[153]. This conserved sequence participates in a number of noncanonical interactions which destabilize the ROSE element and nucleate unfolding at elevated temperatures[154]. ROSE elements, and RNA thermometers more broadly, have remained challenging for obtaining converged free-energy profiles from MD simulations. Here, we show that our computational pipeline can map the full dynamical ensemble of an RNAT. Specifically, we model the conformational landscape of the microROSE element, a 29-nt truncation of the 3’ adjacent stem-loop IV which contains the Shine-Dalgarno sequence.

#### Folding Energy Landscapes

After executing the hierarchical multiscale sampling protocol described in Section A., we obtain 350 unbiased, all-atom MD trajectories of the microROSE RNAT at 310K and 125 trajectories at 450K. The total cumulative simulation time is ∼385 *µs*, with ∼308 *µs* performed at 310K and ∼77 *µs* performed at 450K; details are given in Fig. S1(c).

A LaTF model is trained on the 310K simulation data using a 50 ns lag time, with key architectural and training hyperparameters selected by GMRQ score–based cross-validation (see Fig. S23 and Methods). The LaTF model learns a highly heterogeneous, rugged energy landscape consisting of 32 distinct metastable states as shown in Fig. 4(a). Three of those 32 states (S2, S10, S16) display significant similarity to the available microROSE NMR structure (PDB ID: 2GIO)[154] and are designated as the folded sub-ensemble. Throughout analysis of the micro-ROSE RNAT, structural similarity is assessed using both traditional RMSD and *ɛ*RMSD, a continuous metric defined by Bottaro *et al.*[59] that quantifies differences between the interaction networks of two structures. While S1 is the most populated state, its base-pairing patterns around the internal and apical loops deviate from those observed in the NMR-resolved native structures, hence its exclusion from the folded sub-ensemble.

**FIG. 4.**
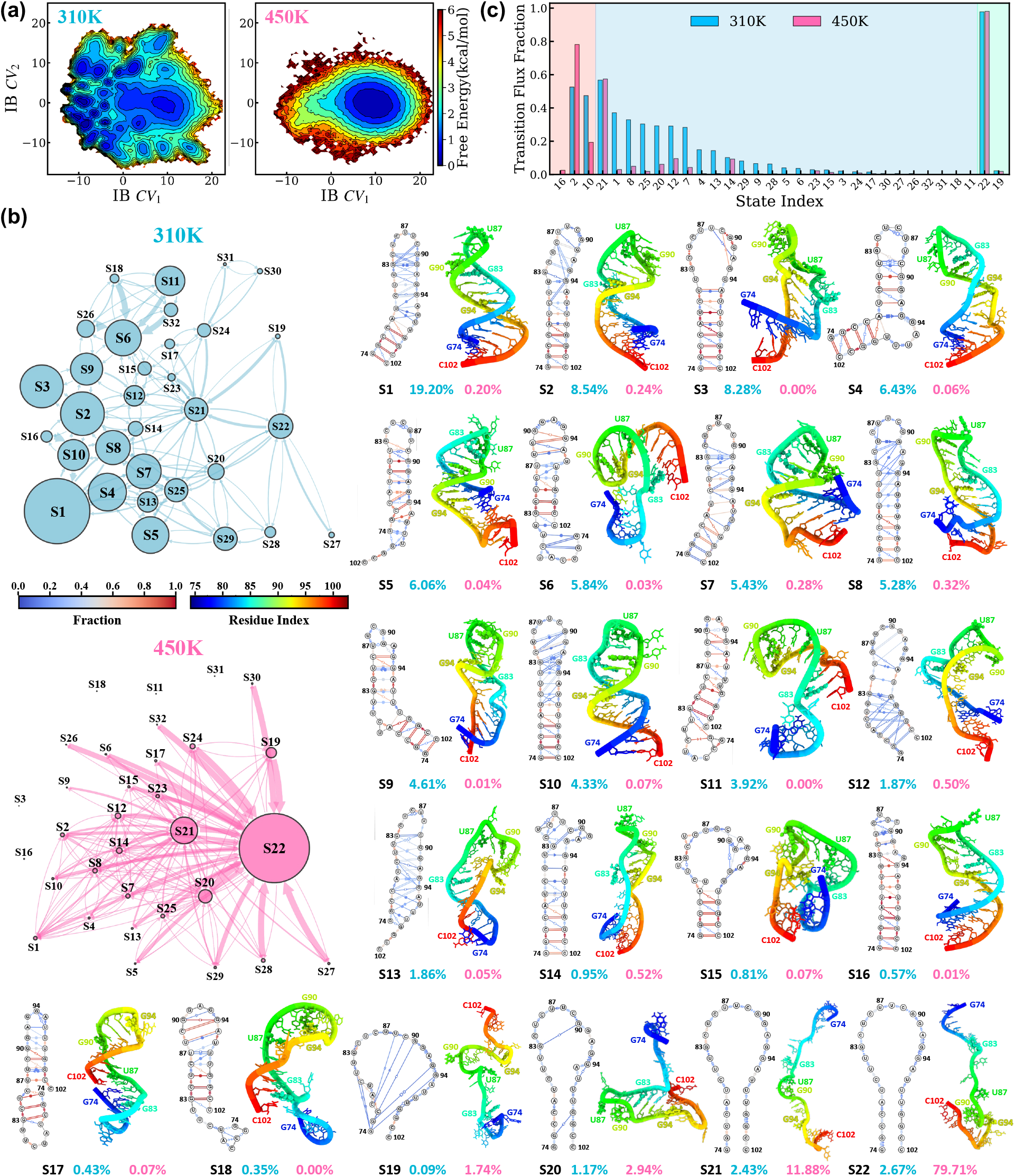
Folding free-energy landscape and kinetics of the MicroROSE RNA Thermometer at 310K and 450K. (a) Free-energy surfaces projected onto the IB space at the two temperatures. Free energies are estimated by reweighting the raw simulation data using the state stationary populations obtained from the corresponding MSMs. (b) Folding kinetic networks at the two temperatures, where transition arrow widths are proportional to the rates (inverse MFPTs) and state populations are indicated by state circle area. Dynamic secondary structure representations are constructed from 500 conformations randomly sampled from each metastable state, where structural annotations follow the Leontis-Westhof classification[104], and the color scheme denotes the fraction of conformations exhibiting each interaction pattern. The dominant tertiary structure of each state, ranked highest by the LaTF model, is also shown. (c) Cumulative folding transition-pathway fluxes through each metastable state at the two temperatures. Folding pathways and their associated fluxes are identified and quantified using TPT in conjunction with the MSMs. Source unfolded states (orange), intermediate states (blue), and sink folded states (green) are categorized and shown accordingly.

Beyond the folded sub-ensemble, we also observe extensive representation of misfolded and partially folded conformational states within our learned free-energy landscapes. Indeed, half of the basins identified from the low-temperature data can be classified as misfolded, illustrating the many thermodynamically degenerate alternate structures that are possible even for a 29nt system. At 450K, however, the rich heterogeneity of the 310K landscape collapses and only a single unfolded basin remains. Many of the misfolded regions of the landscape completely disappear at 450K, while some “native”-like partially folded regions remain—although they are still rarely visited (Fig. S28).

#### Metastable States & Transition Kinetics

We employ MSMs to gain kinetic and thermodynamic insight into the folding free-energy landscapes learned by the LaTF model. Conformations sampled at 310K and 450K are assigned to metastable states identified by the LaTF model trained on the 310K data. This state assignment is visualized for both the low and high temperature cases in Fig. S28 and enables direct comparison of state populations and transition rates across temperatures (Fig. 4(b)). While the metastable-state definitions are shared across temperatures, the Markovian lag time used to estimate transition probabilities is determined independently for each temperature. Details of MSM construction and lag-time validation are provided in the Methods, SI and Fig. S24.

The 32 states identified by the LaTF model can be classified into four categories: folded, partially folded, unfolded and misfolded. As discussed previously, states 2, 10, and 16 are designated as the folded sub-ensemble due to their low RMSD and *ɛ*RMSD to the existing NMR structures. Inspection of the dynamic secondary structures in Fig. 4(b) reveals four states (S19-S22) which lack any persistent structure and thus are chosen to comprise the unfolded sub-ensemble. For the remaining 25 states, partially folded structures are distinguished from mis-folds by correct on-register identification of the four G-C stem pairs. This definition yields 9 partially folded states (S1, S3, S4, S7, S8, S9, S12, S14, S15), with the remaining 16 states classified as misfolded intermediates (S5, S6, S13, S17, S18 shown in Fig. 4; all other misfolded states are shown in Fig. S25).

The misfolded states can be further separated into 3 classes based on the following structural features: states with an off-register stem resulting in a 5’ overhang (S11, S18, S26, S30, S31), states with an off-register stem resulting in a 3’ overhang (S5, S13, S25, S27-S29), and states containing two stacked helices (S6, S17, S24, S32). The 3’ overhang states are least depleted by temperature increase, which suggests greater stability as compared to the 5’ overhang or 2 helix classes. Conversely, the 5’ overhang states appear to be the least stable. Transitions out of states in this class are among the fastest seen at 310K (e.g., S26 to S6, S18 to S6, S11 to S6), while at 450K, they are almost entirely unpopulated. Lastly, although technically classified as misfolded, state 23 is assigned to the unfolded sub-ensemble as it displays significant disorder and appears unlikely to function as a kinetic trap. Fig. S28 displays the free energy landscapes for both low and high temperatures colored by the above categories: folded, partially folded, misfolded-3’ overhang, misfolded-5’ overhang, misfolded-2 helices, and unfolded. The organization of these conformationally distinct classes into specific regions of the latent IB space illustrates the LaTF model’s ability to integrate several short simulations into an interpretable, mechanistically informative description of microROSE structural dynamics.

The TPM for the low and high temperature MSMs both have a single eigenvalue of 1.0, indicating sufficient sampling and an ergodic, fully connected free-energy landscape (Fig. S24). While the 310K eigenspectrum lacks clear timescale separation, a spectral gap does appear in the 450K eigenspectrum between the first 2-3 slow processes and the remaining fast modes, indicating a temperature-induced softening of the landscape ruggedness. MSM analysis reveals that, at 450K, unfolded states (S19–S23) account for more than 95% of the stationary population, whereas only two partially folded states (S12 and S14) retain populations above 0.05%.

#### Folding Pathways

TPT[119, 120] is used to identify kinetic folding pathways at two temperatures from our constructed MSMs. To calculate reactive fluxes, S19 and S22, both highly disordered unfolded states, are chosen as sources and the folded sub-ensemble (S2, S10, S16) is chosen as a sink.

Figs. 4(c) & S26 reveal substantial diversity in the dominant folding pathways at 310K, with multiple partially folded states (S1, S8, S12, S7, S4, S14, S9) carrying significant flux. These states provide the correct stem scaffold needed to form native-like loop contacts and function as key folding intermediates at both low and high temperatures. Among them, S8, S12, and S14 retain substantial flux at 450K and are visited by dominant pathways at both temperatures (Figs. S26–S27), highlighting their persistent roles as on-pathway intermediates.

In addition to partially folded states, unfolded and mis-folded conformational states also appear along the dominant folding pathways. Despite their classification as unfolded, S20 and S21 organize the microROSE RNAT into a collapsed conformation and act as intermediates in nearly all dominant folding pathways regardless of temperature. Among misfolded states, S25 and S13 are most important for folding at 310K: S13 is visited in the 2nd most dominant folding pathway, while both are visited in the 3rd most dominant folding pathway (Fig. S26). Notably, S25 and S13, as well as the majority of misfolded states with non-negligible flux at 310K (S25, S13, S29, S28, S5), belong to the 3’ overhang class. The misfolded states with two stacked helices (S6, S24, S17) receive some flux at 310K, although less than the 3’ overhang states, and the misfolded states with a 5’ overhang (S11, S18, S26, S30, S31) do not carry any flux. These results suggest that at 310K, S25 and S13 act as on-pathway intermediates, while the 5’ overhang and two helix conformational classes are strongly disfavored during folding. At high temperature, the ruggedness of the 310K landscape is significantly reduced. While S25 continues to function as an intermediate at 450K, most other mis-folded states receive little flux. The depletion of mis-folded states at high temperature reduces kinetic heterogeneity and funnels the resulting folding pathways. This funneling is clearly visible when comparing panels (a) and (b) in Fig. S28 and is further illustrated by the presence of single-step folding processes within the dominant pathways at 450K (Fig. S27).

#### Experimental Validations & Other Computational Results

In 2006, Chowdhury *et al.* estimated a structure of the microROSE RNAT using NOE restraints derived from solution NMR (PDB ID: 2GIO)[154]. While others have studied this RNAT or the miniROSE construct via both simulation and experiment[155–158], the work of Chowdhury and colleagues provides the most detailed structural information and is the most applicable to our RNA-only MD simulations. We therefore perform our experimental validation using the 2GIO PDB ensemble, as well as the reported NOE restraints, as a reference. The reported NMR ensemble consists of 20 structures which display very small fluctuations. The largest pair-wise all-atom RMSD between structures in this ensemble is 1.45Å, and when projected onto the learned conformational free-energy landscape by the LaTF model in Fig. S28, the NMR structures all lie on nearly a single point in IB space within metastable state S2. To preserve this overall structural homogeneity while incorporating local conformational flexibility, structures are randomly sampled from state S2 to serve as a comparison. Measuring the correlation between the inter-proton distances derived from NMR models and the corresponding distances from the S2 sampled structures yields an *r*^2^ of 0.67, indicating fair agreement between the simulation-derived and NMR-resolved structures. Furthermore, the state S2 inter-proton distances fall outside the reported experimental uncertainty by more than 1Å for only 3/338 restraints; all 3 violations are for restraints between residues in the flexible apical loop (Fig. S31).

Beyond inter-proton distances, we also perform detailed analysis of the noncanonical interactions that are critical for microROSE temperature sensing. Specifically, we look at distributions of the displacement vector 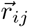 between two interacting bases expressed in local spherical coordinates (schematic shown in Fig. S8(a)). We observe excellent agreement between the NMR 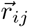 values and the corresponding 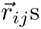 averaged over the simulation folded sub-ensemble for base pairs U79-U97 and A78-U98 (Fig. S29 (d)-(e)). In contrast, the NMR structures display more heterogenous interaction patterns for U82-A95, U81-U96, and C80-U96, which are captured by our simulations but do not always dominate (Fig. S29 (a)-(c)). Similarly, while Chowdhury *et al.* report a G83-G94 *syn*-*anti* base pair, we find that G83-G94 stacking interactions dominate the simulation folded sub-ensemble, with G83 forming the *syn*-*anti* base pair with G94 for only ∼13% of structures (Fig. S30). However, we note that two hydrogen bonds of the G83-G94 pair were enforced as constraints during refinement of the NMR ensemble and assert that is reasonable for the microROSE RNAT to occupy a larger range of conformations, as seen in our extensive simulations, without these constraints.

Finally, we evaluate multiple tertiary structure modeling approaches on the microROSE RNAT (Fig. S32). As with the GCAA TL and HIV-TAR results, AlphaFold3[26] and Boltz2[80] predictions are limited to the NMR native state and only sample local fluctuations. FARFAR2[76] provides good coverage of the folded and partially folded regions of the landscape, but rarely samples the misfolded or unfolded regions. 3dRNA[103] achieves the most extensive sampling and does predict unfolded structures, however it fails to predict many misfolded states, particularly those in the 5’-overhang class. Our results across systems clearly show that top-down computational approaches are insufficient to fully sample heterogenous RNA conformational land-scapes and demonstrate the need for integrated physics-AI approaches.

## III. CONCLUSIONS & DISCUSSION

In this work, we present a hierarchical multiscale computational framework for *de novo* mapping of RNA conformational landscapes, structural ensembles, and folding kinetics, with application to three RNA hairpins: the GCAA tetraloop, HIV-TAR stem-loop, and micro-ROSE RNA thermometer. Through a series of complementary modeling strategies, our framework combines broad conformational exploration with resolution of dynamic modes across multiple timescales. Hierarchical structure generation first provides diverse starting conformations spanning distinct regions of rugged RNA free-energy landscapes. We note that alternative states characterized by unusual noncanonical interactions may be difficult to capture in this generation stage, as such motifs are sparsely represented in structural databases and fragment libraries. Following heterogenous initial ensemble generation, subsequent unbiased MD simulations allow putative structures to relax and explore broader conformational space. Generative-AI-guided adaptive sampling combines information across temperature to promote exploration of slow folding–unfolding transitions and inter-conversion between various excited states. After multiple rounds of simulation, AI-augmented representation learning identifies low-dimensional CVs and long-lived metastable states. Finally, these state assignments are used to build Markov state models capable of integrating parallel short trajectories to resolve long-timescale kinetics and temperature-dependent folding mechanisms. Across all three RNA systems, we systematically benchmark our results against prior computational results and experimental measurements, including NMR NOEs and RDCs, and obtain overall fair agreement.

While the systematic characterization of rugged RNA conformational landscapes using unbiased, atomistic, explicit-solvent simulations remains limited, the framework developed here offers a general route to resolving RNA equilibrium ensembles and kinetics. Moreover, our results underscore several challenges that warrant further investigation. To begin, RNA structural transitions involve dynamic modes coupled across multiple spatial and temporal scales, which complicate the design of structural descriptors that accurately capture base-pairing, stacking, and higher-order conformational rearrangements. The present work employs distance-based features, however, angular descriptors and representations incorporating ion and solvent degrees of freedom may provide additional, complementary information. Increasingly rich feature spaces also complicate the identification of low-dimensional, physically interpretable CVs capable of separating numerous metastable states. This challenge is particularly consequential for CV-based enhanced-sampling approaches, for which informative coordinates such as the IB CVs identified here are difficult to derive without substantial prior sampling of the underlying conformational landscape.

RNA equilibrium ensembles and structural dynamics are also strongly influenced by environmental conditions. While we investigate the effect of temperature here, factors such as ion concentration, pH, and metabolite abundance also substantially reshape the conformational landscape and should be probed in future work. We additionally acknowledge the inaccuracies present in currently available RNA force fields—which have been well documented by previous work [41–49]—and expect that our results could change depending on the force field used for simulations. Nevertheless, we believe our current pipeline can aid future force field developers, as it provides a systematic and self-consistent protocol for generating conformational ensembles that can be compared with experimental observables to determine force field corrections. More broadly, the comprehensive structural ensembles generated by this approach provide a robust basis for subsequent experimental reweighting to obtain improved agreement with measured data.

In closing, we stress the importance of integrating physics with AI to recover Boltzmann-distributed conformational ensembles. When comparing thermodynamics-based tertiary structure prediction methods to purely AI methods, the AI methods including AlphaFold3[26]and Boltz2[80] replicate PDB structures included in their training data and are unable to sample beyond local fluctuations (Figs. S10, S22, S32). While there currently exist multiple methods to generate ensembles or alternative conformations from AI protein structure prediction models[159–163], similar work has not yet been done for RNA—despite the fact that training data memorization is a much greater issue for AI RNA structure prediction models. Our combination of hierarchical seeding, generative modeling, and adaptive MD simulations thus addresses an important gap in current ensemble generation approaches for dynamic RNA systems, while providing a foundation for further developments such as experimental reweighting, explicit treatment of cellular metabolites, and integration with downstream docking workflows.

## IV. METHODS

### Computational Modeling of RNA Secondary and Tertiary Structures

A multi-method secondary structure prediction approach is employed to maximize structural diversity. Specifically, we combine thermodynamics-based methods (e.g., RNAstructure[69], ViennaRNA[70], and mFold[164]), data-driven approaches (e.g., CONTRAfold[165] and EternaFold[71]), and deep-learning models (MXfold2)[78] to generate candidate secondary structures from a given sequence. Each secondary structure is then assembled into tertiary models using FARFAR2[76], followed by Monte Carlo sampling with a knowledge-based coarse-grained scoring function and subsequent refinement using a knowledge-based all-atom scoring function. Repeated assembly and refinement for each secondary structure yields a diverse tertiary structure ensemble. The resulting models are featurized using ***G***-vectors[58], followed by PCA for dimensionality reduction. After clustering in PC space, each cluster center is mapped to the nearest FARFAR2[76] structure to generate heterogeneous initial decoys for MD simulations. For comparison, tertiary structure predictions are also conducted using 3dRNA[74, 103], AlphaFold3[26], and Boltz-2[80]. Detailed settings and commands for all secondary and tertiary structure prediction methods are provided in the SI.

### Molecular Dynamics Simulation

Unbiased all-atom MD simulations in explicit solvent are initiated from the selected structural decoys. Each system is placed in a rhombic dodecahedral periodic box sized to maintain at least 1.2 nm separation between unfolded conformations and their periodic images. The GCAA tetraloop is solvated in 1.0 M KCl, whereas the HIV-TAR stem-loop and microROSE RNA are solvated in 0.1 M NaCl, using TIP4P-D water. RNA dynamics are simulated in GROMACS[166] with the DESRES-AMBER 2022 force field[87] and CHARMM22 ion parameters[167]. Prior to production, each system undergoes energy minimization followed by multistep *NVT* and *NPT* equilibrations with progressively released positional restraints. Production *NVT* simulations are then performed at both a lower temperature (∼300K) and an elevated temperature (*>* 400 K) for each system. Further simulation details are provided in the SI.

### LaTF-Accelerated Adaptive Sampling

Following the initial unbiased MD simulations, data accumulated across adaptive-sampling rounds at both low and high temperatures are used to iteratively train LaTF models that guide subsequent reseeding. By jointly leveraging information across temperatures, LaTF improves estimation of the temperature-dependent free-energy landscapes under limited sampling and generates latent distributions for both conditions[99]. New seeds are selected using k-means and k-centers clustering, with k-means following the inferred distribution and k-centers promoting more uniform coverage of latent space. This adaptive sampling procedure is repeated for three to four rounds, after which a final LaTF model is trained using only the low-temperature data for downstream analysis. Five-fold GMRQ score based cross-validation[100] is used to optimize initial-state labeling, model architecture, and training hyperparameters. Further details of the adaptive-sampling protocol, LaTF formulation, and training procedure are provided in the SI.

### Markov State Model Construction & Validation

MSMs are constructed separately for the low- and high-temperature data to provide quantitative descriptions of temperature-dependent RNA folding thermodynamics and kinetics. MD-sampled conformations are assigned to metastable states using the LaTF model trained at low temperature. The LaTF model takes ***r*** -vector [58] norms and pairwise 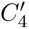 distances as input structural features and determines an appropriate state decomposition at a predefined lag time, with short-lived states merged into longer-lived metastable states. The Markovian lag time for each MSM is selected based on implied-timescale convergence, and model validity is further assessed using Chapman–Kolmogorov tests. Additional details of MSM construction and validation are provided in the SI.

## V. DATA & CODE AVAILABILITY

The simulation data associated with this research is available upon request. The source code for Latent Thermodynamic Flows model and associated documentation is available at https://github.com/tiwarylab/ LatentThermoFlows.

## VI. ACKNOWLEDGEMENTS

The authors acknowledge support from the National Science Foundation under Grant No. CHE-2044165. Y.Q. acknowledges support from startup funds provided by the University of Notre Dame. Computational resources are provided by the University of Maryland HPC cluster Zaratan, NSF ACCESS (Project CHE180027P), and the University of Notre Dame Center for Research Computing. The authors thank Dr. Lukas Herron and Dr. Da Teng for fruitful discussions during the preparation of this work. P.T. is an investigator at the University of Maryland Institute for Health Computing, which is supported by funding from Montgomery County, Maryland and The University of Maryland Strategic Partnership: MPowering the State, a formal collaboration between the University of Maryland, College Park, and the University of Maryland, Baltimore.

## Supplementary Information

### I. REVIEW & COMPARISON OF PREVIOUS STUDIES OF RNAS

**Table S1:**
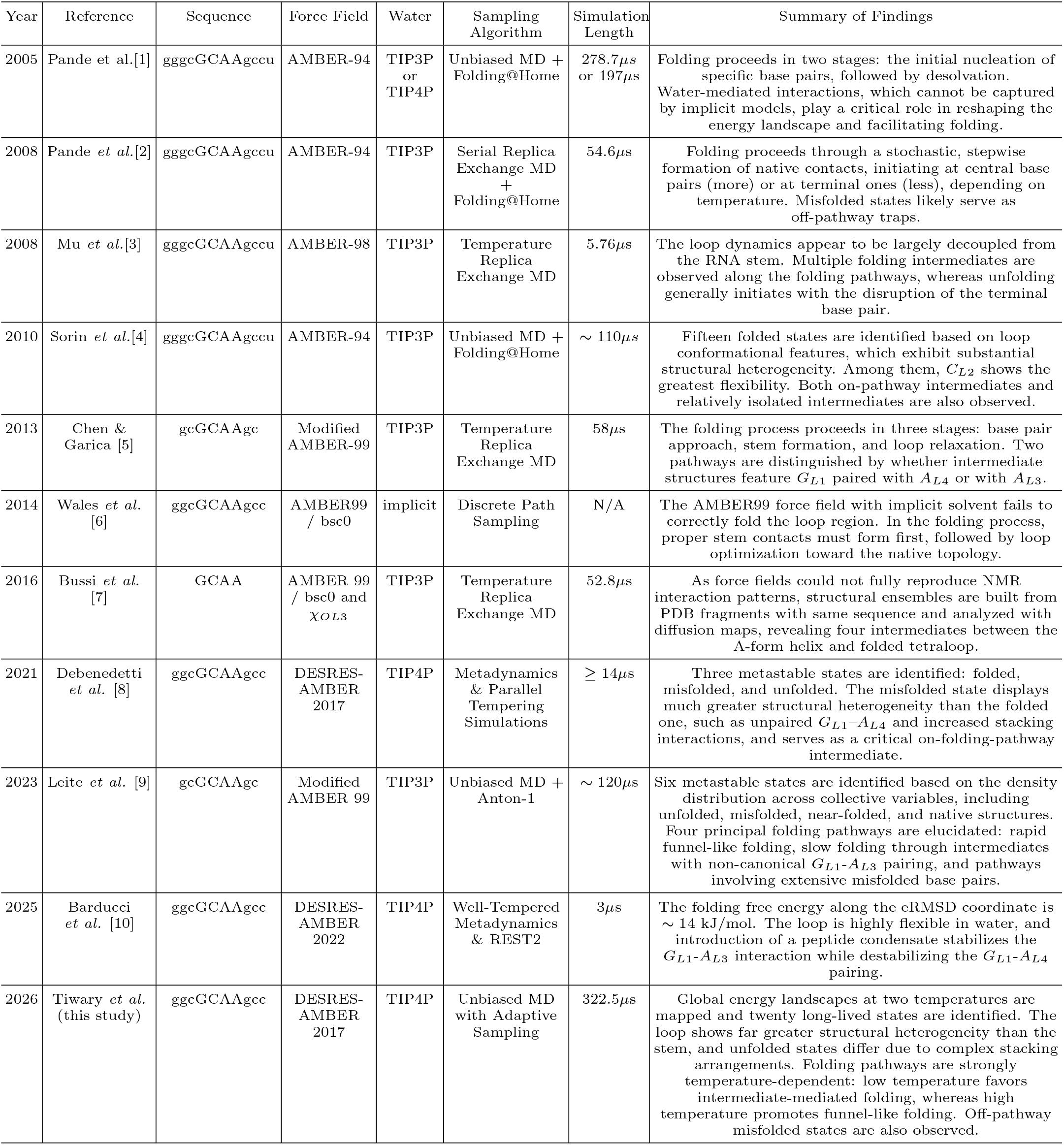
Overview of Major Computational Investigations of the GCAA RNA Tetraloop.

**Table S2:**
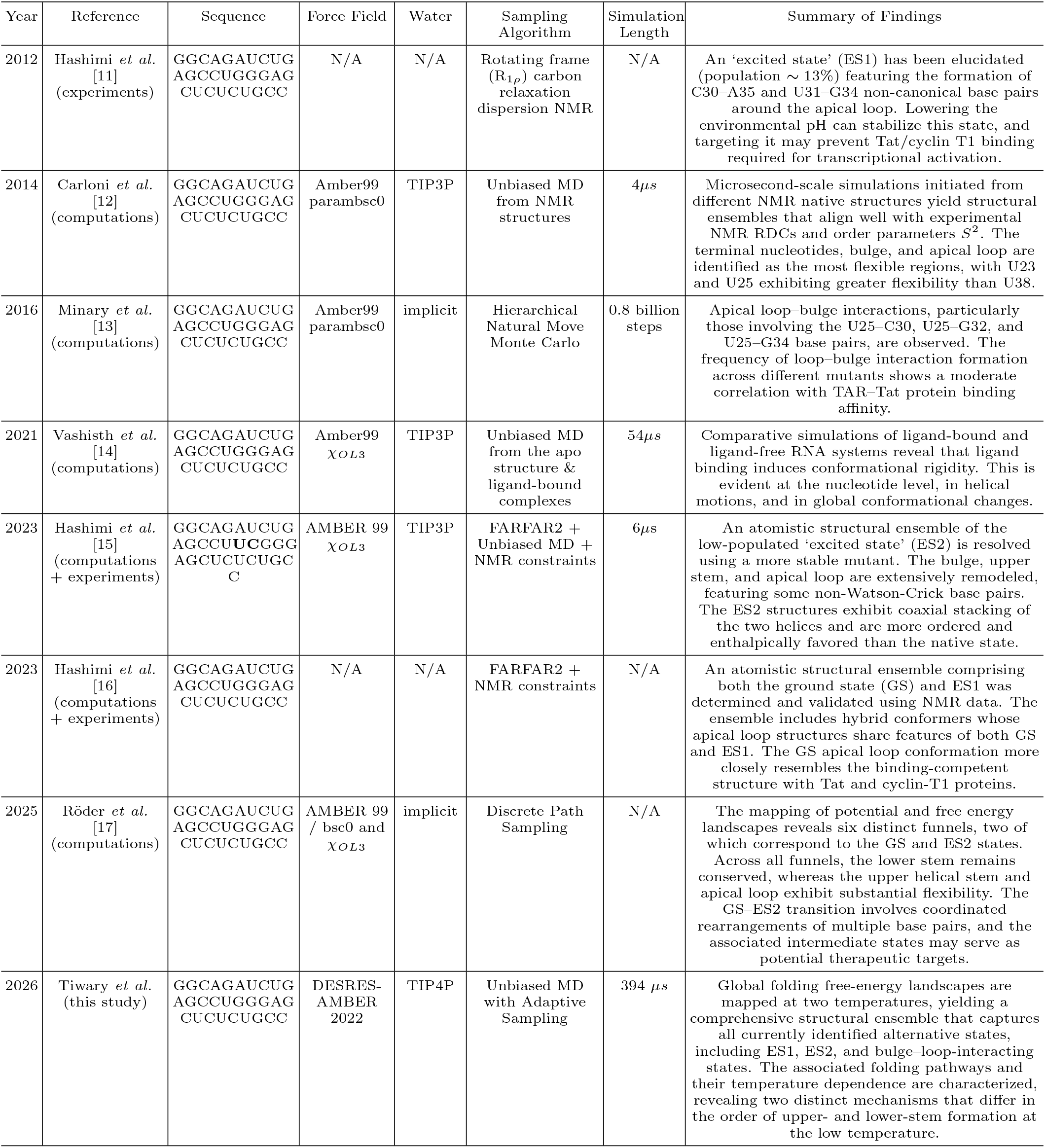
Overview of Major Computational/Experimental Investigations of the HIV-TAR RNA.

**Table S3:**
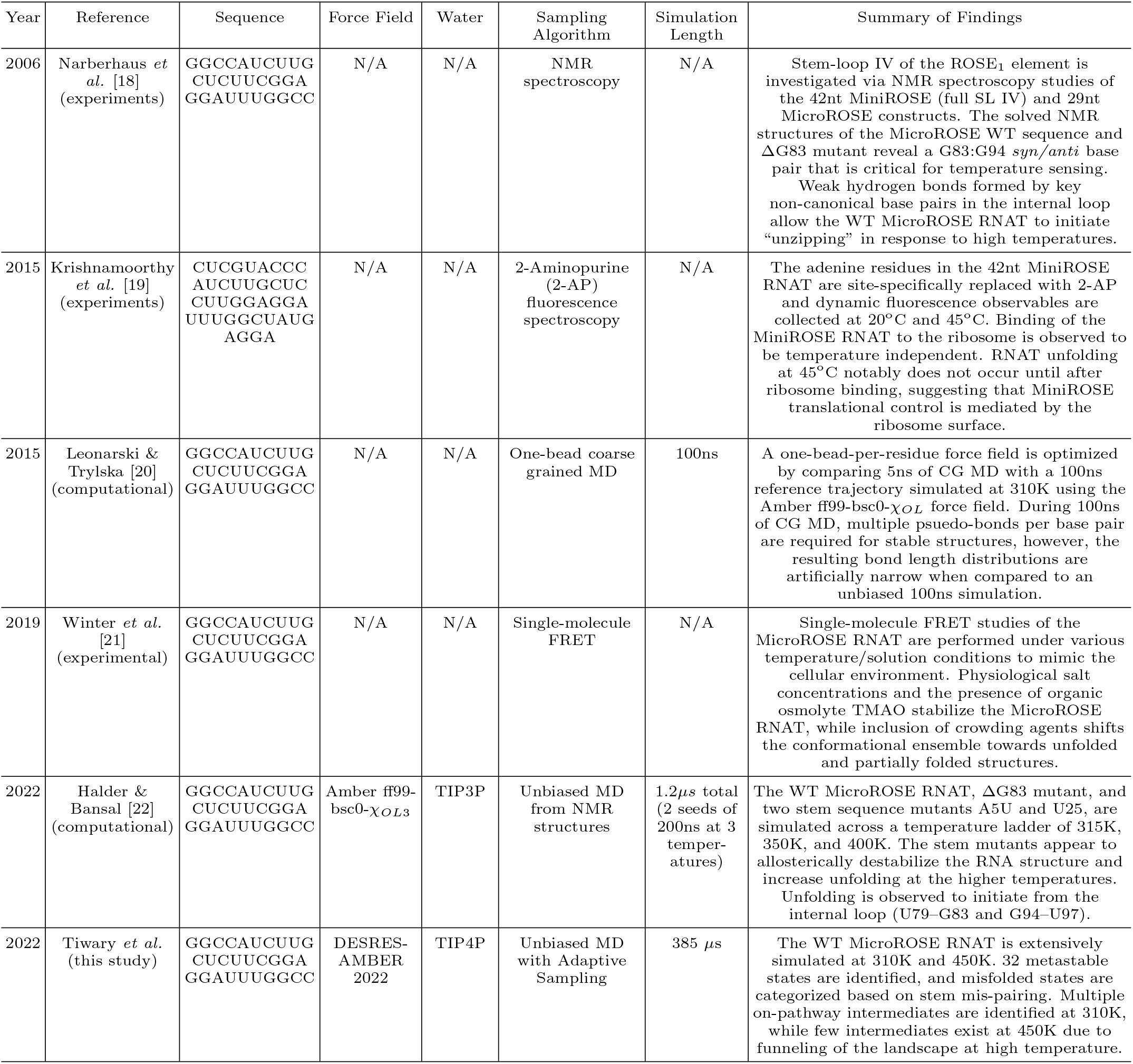
Overview of Major Computational/Experimental Investigations of the MicroROSE RNA Thermometer.

### II. COMPUTATIONAL MODELING OF RNA SECONDARY STRUCTURES

Over the past decades, numerous methods and algorithms have been developed to predict RNA secondary structures from sequence information. Broadly, these approaches can be classified into two categories: knowledge-based methods and learning-based methods. Knowledge-based methods incorporate prior information, usually derived from experimental observations or statistical analyses, into explicit search or prediction algorithms. In contrast, learning-based methods employ trainable models that directly map RNA sequences to secondary structures. These models are optimized in a supervised manner to accurately reproduce experimentally resolved structures. Knowledge-based methods can be further subdivided into two classes: (1) thermodynamics-based methods, which integrate experimentally derived free-energy parameters for individual structural motifs together with dynamic programming algorithms, and (2) covariation-based methods, which exploit structurally and functionally relevant base-pairing signals inferred from multiple sequence alignments. At present, thermodynamic approaches dominate RNA secondary structure prediction[23] due to their relatively good performance in data-scarce regimes, exploration of suboptimal structural ensembles, and ability to incorporate environmental factors such as temperature and ionic conditions. In this study, we employ a suite of primarily thermodynamics-based methods to systematically explore the RNA secondary structure space of three systems. Our strategy aims to capture the intrinsic structural heterogeneity of these RNAs and to generate sufficiently diverse initial conformations for subsequent tertiary structure modeling.

#### A. GCAA Tetraloop

Given the short sequence length and relatively limited secondary structural heterogeneity of the GCAA tetraloop, as detailed in our previous work[24], we only employ the *ViennaRNA-2.7.0* package [25] (thermodynamic-based method) with following command to construct its secondary structure ensemble from sequence:

`RNAsubopt -s -T 37 -d 3 -e 4 --salt 1.021 -N -i gcaa_tetraloop_fasta.fasta`

This approach yields the minimum free-energy structure along with two suboptimal conformations, which primarily differ in their base-pairing patterns within the helical stem region.

#### B. HIV-TAR RNA Stemloop

A comprehensive integration of six methods, including both thermodynamics-based and learning–based approaches, is employed to generate a ensemble of 365 distinct secondary structures for the HIV-TAR stemloop. The implementation details, including algorithmic settings, execution commands, and the number of structures produced by each method, are summarized as below and in Table S4:

(1&2). The following commands are used with the *ViennaRNA-2.7.0* package[25], a purely thermodynamics-based approach, to identify both optimal and suboptimal secondary structures, as well as structures containing pseudoknots. This prediction yields 288 suboptimal structures and 20 structures with pseudoknots:

`RNAsubopt -s -T 37 -d 3 -e 10 --salt 1.021 -N -i hiv_tar_1ANR_fasta.fasta`

`echo “GGCAGAUCUGAGCCUGGGAGCUCUCUGCC” | RNAPKplex -T 37 --salt 1.021 -c 1e-7 -s 15 -e -10`

We manually set a relatively large energy score for pseudoknot formation to exhaustively explore possible structures. It is worth noting, however, that the *PKplex* algorithm can predict at most one pseudoknot per sequence. We expect structures containing multiple pseudoknots to be relatively rare given the length of the RNAs studied here (29 nts). Thus, we posit that our implementation of *ViennaRNA-2.7.0* provides comprehensive and reasonable structural ensembles.

(3). Additionally, *RNAstructure-6.5* [26], another thermodynamics-based method built upon slightly different nearest-neighbor thermodynamic parameters[27–29], is employed to validate and complement the secondary structure predictions obtained using *ViennaRNA-2.7.0*. We execute the following command:

`RNAstructure/exe/AllSubjupyter-notebook --no-browser --ip=0.0.0.0 --port=8890 hiv_tar_1ANR_fasta.txt output.ct -a 10 --alphabet rna -p 100 -t 310.15`

where the two thresholds constraining the maximum absolute and relative energy deviations from the minimum free energy structure are set to 10 kcal/mol and 100% respectively. This choice ensures that we are able to comprehensively explore suboptimal structures. In total, we obtain 364 structures from *RNAstructure-6.5*.

(4). To further explore suboptimal structural possibilities, the *UNAFold (mfold)* package[30, 31], an alternative thermodynamics-based method, is also utilized. Folding predictions are performed at 310 K under ionic conditions of 1 M NaCl without divalent cations. The percent suboptimality threshold is set to 100%, allowing prediction of structures with free energies up to 100% above the minimum free energy. This yields a total of 40 suboptimal structures.

(5). Given that many RNAs may not predominantly adopt the minimum free energy or maximum likelihood structures predicted by standard models[32–34], we also utilize *CentroidFold* to reassess the ensemble of secondary structures[35, 36]. *CentroidFold* applies the *γ* centroid estimator, which ranks structures based on their expected accuracy derived from base-pairing probabilities predicted by thermodynamic[37] or machine-learning models (e.g., *CONTRAfold* [38]). By varying *γ* across a broad range and incorporating all available base-pair probability predictors, we identify a total of 7 representative candidate structures.

(6). While deep learning has achieved remarkable advances in RNA secondary structure prediction, we adopt a hybrid approach, *MXfold2* [39], which integrates thermodynamic-based and deep learning–based approaches. Specifically, *MXfold2* combines physics-based free-energy terms with neural network–derived scores within a dynamic programming framework to identify the optimal folded structure. Its max-margin training strategy enables the model to capture intrinsic correlations in RNA sequences by leveraging the large sequence–structure dataset. This approach offers a single candidate secondary structure.

**Table S4:** Number of secondary structures for HIV-TAR RNA predicted by methods based on different principles.

| Methods | Thermodynamic |  |  |  |  | Machine Learning |
| --- | --- | --- | --- | --- | --- | --- |
|  | ViennaRNA<br>RNAsubopt | ViennaRNA<br>RNApkplex | RNAstructure<br>Allsub | mFold | CentroidFold | MXfold2 |
| # of predicted secondary structures | 288 | 20 | 364 | 40 | 7 | 1 |
| # of accumulated non-redundant<br>secondary structures | 288 | 308 | 344 | 363 | 365 | 365 |

#### C. MicroROSE RNA Thermometer

Secondary structure prediction is performed using a multi-method approach to generate a diverse ensemble of 1221 distinct secondary structures for the MicroROSE RNA thermometer. Three methods are used in this analysis: *Eternafold* [40], *LinearFold* [41] and *LinearAlifold* [42]. All implementation details, including explicit execution commands, are provided below. Table S5 details the number of structures produced by each method and the cumulative number of unique structures generated.

(1) *Eternafold* is a learning-based method that is created by extending the loss function of *CONTRAfold* [38], also a learning-based method, to optimize for multiple ensemble property prediction tasks. To be explicit, while *CONTRAfold* is trained to correctly predict individual secondary structures, *Eternafold* extends this framework to include accurate prediction of chemical mapping signals and riboswitch protein binding affinities both with and without ligand. We run the following command to generate 5000 stochastic samples from the learned partition function:

`contrafold sample 2gio.fasta --params parameters/EternaFoldParams.v1 --nsamples 5000`

(2) *LinearFold* modifies the usual dynamic programming algorithms used by nearest-neighbor models, which scale cubically with sequence length, to provide linear run-time secondary structure prediction. *LinearFold* contains implementations of the thermodynamics-based *ViennaRNA*[25] and the learning-based *CONTRAfold* models. Both energy models are run using the following commands:

`echo GGCCAUCUUGCUCUUCGGAGGAUUUGGCC | ./linearfold --zuker --delta 1000 -V`

`echo GGCCAUCUUGCUCUUCGGAGGAUUUGGCC | ./linearfold --zuker --delta 1000`

*Linearfold* by default runs using the *CONTRAfold* parameters, and the addition of the -V flag switches the implementation to the *ViennaRNA* parameters. We set the -delta flag to 1000 to return suboptimal structures within 1000 kcal/mol of the optimal structure. This essentially includes all generated suboptimal structures in our ensemble, thereby maximizing exploration of possible secondary structures.

(3) *LinearAlifold* uses the dynamic programming algorithm reported in *Linearfold* to predict consensus secondary structures from multiple aligned homologous RNA sequences. It also estimates a partition function which can be stochastically sampled. *LinearAlifold* contains implementations of the thermodynamics-based *ViennaRNA* and hybrid *BL\** energy models[43], both of which are run in this analysis. The loss function used to find the *BL\** energy parameters is similar to that of *CONTRAfold*, however both thermodynamic and structural data are used in the likelihood maximization. Thus, we term *BL\** a hybrid thermodynamics- and learning-based method. We run *LinearAlifold* using the following commands:

`cat 2gio.fasta | ./linearalifold -s 20000 -e 1`

`cat 2gio.fasta | ./linearalifold -s 20000 -e 2`

The -s 20000 flag returns 20,000 stochastic samples of the learned partition function and -e toggles the parameterization between *ViennaRNA* (1) and *BL\** (2). We also run the following commands to compute the ThreshKnot[44], Centroid and Maximum Expected Accuracy[32] structures. However, these structures are not unique and have already been generated during the stochastic sampling step, so the commands are omitted from Table S5.

`cat 2gio.fasta | ./linearalifold -p -e 1`

`cat 2gio.fasta | ./linearalifold -p -e 2`

**Table S5:**
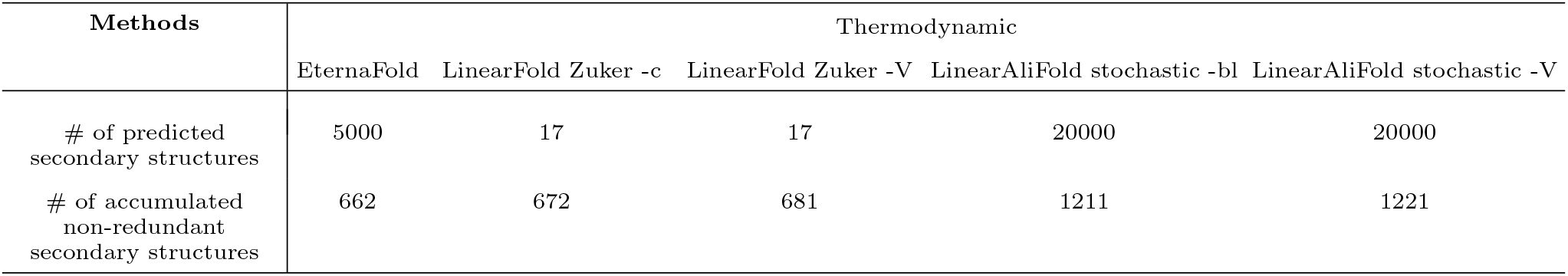
Number of secondary structures predicted for MicroRose RNA Thermometer by multi-method approach.

### III. COMPUTATIONAL MODELING OF RNA TERTIARY STRUCTURES

#### A. FARFAR2

FARFAR2[45], a fragment-assembly structural modeling protocol implemented in the Rosetta `rna_denovo` pro-gram, is employed to generate tertiary structures from the given RNA sequence and its corresponding ensemble of predicted secondary structures. For each secondary structure, the sequence is decomposed into a series of overlapping fragment windows, and candidate structural fragments are identified from a library of experimentally determined RNA structures. These fragments are subsequently assembled to construct full-length tertiary structure models. The assembled structures are then subjected to Monte Carlo sampling using a low-resolution, coarse-grained, knowledge-based scoring function to refine local conformations and explore structural flexibility. Finally, the resulting models are further optimized using the high-resolution, all-atom Rosetta `rna_hires` scoring function. In our implementation, hundreds of independent Monte Carlo trajectories, each consisting of thousands of sampling cycles, are performed for every secondary structure to extensively explore the RNA conformational landscapes. This approach yields 2,250, 27,375, and 12,210 structures for the tetraloop, HIV-TAR stem–loop, and MicroROSE RNA thermometer systems, respectively. The generated structures are subsequently clustered and used as the only source of initial decoys for downstream unbiased molecular dynamics simulations. In addition, several alternative tertiary structure modeling approaches are executed, as described below, to assess their ability to capture the conformational heterogeneity of different RNAs.

#### B. 3dRNA

Apart from FARFAR2, we employ another fragment-assembly approach, 3dRNA[46–49], to predict RNA tertiary structures and sample conformational flexibility. After decomposing a secondary structure into its smallest constituent secondary structure elements (SSEs), 3dRNA searches a template library for the corresponding structural fragments and assembles them sequentially. Templates with varying degrees of sequence homology are randomly sampled to assemble five structural models for each secondary structure. Subsequent optimization using simulated annealing Monte Carlo enables comprehensive sampling of local conformational fluctuations in the resulting tertiary structures. For each system, sampling is conducted using a knowledge-based scoring function named 3dRNAscore[47] at three Monte Carlo temperatures (i.e., 10, 50, and 100). Structures are recorded every 2,000 Monte Carlo steps, resulting in 50 configurations per temperature. The following commands are executed:

`nsp assemble -name 3dRNA_assembly -seq <SEQUENCE> -ss “<2D STRUCTURE>” -num 5`

`nsp opt -name 3dRNA_opt -seq <SEQUENCE> -ss “<2D STRUCTURE>” -init 3dRNA_assembly.pred.pdb`

`nsp traj cluster opt_structure.pdb -prefix opt_structure.cluster -k 1 -aa`

`nsp fit opt_structure.cluster.1.center.pdb -o opt_fit_structure.pdb`

Each assembled structure is subjected to further optimization and relaxation using the all-atom AMBER14 force field[50] with explicit TIP3P solvent. This protocol consists of 2,000 steps of local energy minimization using the L-BFGS algorithm, followed by 1 ps of *NVT* equilibration at 300 K. Consequently, 2,250, 129,910, and 440,862 structures are generated for the tetraloop, HIV-TAR stemloop, and MicroROSE thermometer systems, respectively. For the latter two systems, structures containing chain breaks or exhibiting extremely poor stability, as indicated by high scoring function values, are excluded during sampling.

#### C. Boltz2 & AlphaFold3

RNA structures are also predicted using the Boltz2[51] and AlphaFold3[52] models, thereby enabling direct comparison between our molecular dynamics results and other state-of-the-art approaches. For the Boltz2 model, we adopt the Boltz-steering inference-time strategy, which incorporates physics-based potentials to enhance the physical plausibility of predicted structures. In parallel, we utilize the generative diffusion model with 200 uniformly spaced step-scale values between 0.8 and 2.0, resulting in 10 structures for each setting to systematically sample RNA structural heterogeneity. The command used for structure generation is as follows:

`boltz predict RNA_boltz2_prediction.yaml --use_msa_server --use_potentials --diffusion_samples 10--recycling_steps 3 --step_scale $scale`

For the AlphaFold3 model, we vary the random seed from 0 to 400 and generate 5 structures per seed, yielding a total of 2,000 structures for each system. All other model settings are kept at their default values.

The resulting structural ensembles generated by the different prediction approaches are featurized using **r**-vectors and pairwise distance descriptors (as detailed in the following section) and subsequently represented in the latent information bottleneck (IB) space encoded by the corresponding trained LaTF models. These visualizations illustrate each method’s ability to capture structural heterogeneity across different RNA systems. In parallel, structure prediction accuracy is assessed using RMSD and ɛRMSD metrics with respect to the corresponding NMR reference structures. The results are shown in Fig. S10, S22 and S32. Because the NMR structures for all three systems were included in the training of the Boltz2 and AlphaFold3 models, the observed low RMSD and ɛRMSD values, and the correspondingly localized and limited structural diversity, are not unexpected.

### IV. DETAILED MOLECULAR DYNAMICS SIMULATION PROTOCOLS

#### A. GCAA Tetraloop

Following the initial structure construction and generative AI-augmented simulation protocol described in our previous work[24], we extend our prior molecular dynamics (MD) simulations to roughly twice their original duration to improve sampling of transitions among metastable states. The system consists of a single GCAA tetraloop solvated in ~ 3900 TIP4P water molecules with 1 M KCl, yielding a total of 15, 000 atoms (Fig.S1(a)). Simulations are performed using the DESRES AMBER 2017[53] force field in combination with CHARMM22 ion parameters. The final dataset comprises 80 trajectories at 300 K with a cumulative simulation time of ~ 322*µ*s and 80 trajectories at 400 K with a cumulative time of ~ 148*µ*s, as shown in Fig.S1(a).

We evaluate the convergence of our adaptive MD simulations using a previous study which employed enhanced sampling with the same force field to explore the folding free energy landscape of the GCAA tetraloop[8]. In this previous work, parallel tempering in the well-tempered ensemble combined with well-tempered metadynamics (PTWTE-WTM) is adopted to sample the free energy surface (FES) as a function of two order parameters, *i.e.*, the root-mean-square deviation (RMSD) and Q, a parameter which quantifies the compactness of inter-strand contacts in the stem region. Q is defined by the following expression:

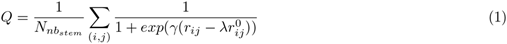

where (*i, j*) denotes a contacting heavy-atom pair in which atom *i* belongs to the strand terminating at the 3*^′^* end and atom *j* belongs to the strand terminating at the 5*^′^* end. 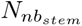 represents the total number of such atom pairs whose separation 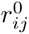 is less than 5^°^*A* in the reference native structure. *r_ij_* is the distance between *i* and *j* in any given instantaneous structure. The smoothing parameter *γ* is set to 50*nm^−^*^1^ , and the hyperparameter *λ* is chosen to be 1.5.

A comparison of the results obtained via PTWTE-WTM and those directly estimated from raw unbiased MD simulations (Fig. S3) demonstrates that our parallel adaptive MD simulations clearly and smoothly sample transitions between folded, misfolded, and unfolded states. Unbiased MD simulations at 300 K also capture many metastable basins in the transition regions, highlighting the ability of our generative AI-augmented unbiased sampling strategy to efficiently explore the global energy landscape.

#### B. HIV-TAR RNA Stemloop

The FARFAR2-generated HIV-TAR tertiary structure ensemble is represented using **G**-vector descriptors and projected onto the first ten principal components obtained from principal component analysis (PCA). The projected structures are subsequently partitioned into 50 clusters using the k-means and k-centers algorithms, respectively. For each cluster, the structure closest to the cluster centroid in the PC space is selected as a representative seed structure for simulations. Unbiased MD trajectories are then initiated from these representative conformations. The HIV-TAR is solvated in a rhombic dodecahedral periodic box with a vector length of 7.4 nm, which is chosen to ensure that even the most extended RNA conformations remain at least 1.2 nm from the box boundaries. To neutralize the negatively charged RNA, Na^+^ ions are added, and additional Cl*^−^* ions are included to achieve a NaCl concentration of 0.1 M. Approximately 9,000 TIP4P-D water molecules are added, resulting in a system comprising 37, 000 atoms in total (Fig.S1(b)). The most recent DESRES AMBER 2022 force field[54] is used to model the RNA, in combination with the CHARMM22 ion parameters[55].

Prior to the production simulation, we perform a multi-step equilibration protocol. Initially, energy minimization is performed for 5,000 steps using the steepest descent algorithm. Subsequently, a 2ns *NVT* relaxation is conducted, during which all RNA atoms are restrained to their energy-minimized positions using a harmonic potential with a force constant of 1000*kJ/*(*mol · nm*^2^). This is followed by four sequential 2ns *NPT* relaxations (total length of 8ns) with progressively reduced restraint force constants of 1000, 100, 10 and 0 *kJ/*(*mol·nm*^2^). Pressure is regulated using the Berendsen barostat[56] during the initial three stages, after which the Parrinello–Rahman coupling method[57] is used for for the final phase of equilibration. Final *NVT* production simulations are performed starting from the equilibrated configurations. Throughout the relaxation and simulation steps, the system temperature is consistently controlled using a velocity-rescaled Berendsen thermostat.[56]

The cutoff for both Coulombic and van der Waals interactions is set to 1.2 nm, and long-range electrostatic interactions are treated using the particle mesh Ewald (PME) method[58]. Simulations are performed using a 2 fs integration time step, while all bonds involving hydrogen atoms are constrained using the LINCS algorithm[59]. The coordinates of the HIV-TAR RNA are recorded every 0.2 ns and are used during analysis. All the MD simulations are conducted using the GROMACS package[60].

Given the pronounced ruggedness of the HIV-TAR RNA energy landscape, rapid exploration of its global features using unbiased MD simulations can be challenging. Here, we implement an adaptive sampling protocol augmented with a generative AI model to accelerate exploration of the rugged energy landscape and enable efficient sampling of transition dynamics between metastable states. The flow-based generative model within the Latent Thermodynamic Flows (LaTF) framework has been shown to accurately infer temperature-dependent free energy surfaces (FES) from limited and sparse simulation data[24]. As such, we perform simulations at two temperatures and train a generative model to guide reseeding of subsequent simulation rounds on-the-fly, thus improving overall sampling efficiency.

Our detailed structure generation protocol provides an initial heterogeneous structural ensemble. Subsequent clustering of these structures yields 100 representative decoys to serve as starting points for simulations. All 100 decoys are employed to initiate independent simulations at 310 K, each run for approximately 1*µs*. To exploit the faster convergence seen at higher temperatures, only the first 50 decoys are used to perform simulations at 420 K, with each replica run for ~ 0.5*µs*. A LaTF model is then trained using the simulation data collected at two temperatures. This model learns a two-dimensional information bottleneck that separates metastable states and predicts the equilibrium distributions at different temperatures. Samples generated at 310 K are separated into 100 clusters in the information bottleneck space using both the k-means and k-centers algorithms; both methods are used to generate 50 clusters for a total of 100. Use of k-means ensures that seed structures will approximately follow the equilibrium distribution, while use of k-centers provides coverage of boundary and less-explored regions of the landscape. Cluster centers are reconstructed as all-atom structures by mapping each center to the nearest simulated data-point in the information bottleneck space. An additional round of parallel simulations is then initiated from these 100 seed structures at 310 K. This adaptive sampling protocol is repeated for three iterations at 310 K. Meanwhile, the existing 50 trajectories at 420 K are continuously extended through three iterations rather than being reseeded.

In total, we obtain 400 trajectories at 310 K with a cumulative simulation time of ∼ 394*µ*s, corresponding to an average trajectory length of ∼ 0.98*µ*s. In addition, we obtain 50 trajectories at 420 K, yielding a cumulative simulation time of ∼ 87*µ*s and an average trajectory length of *∼* 1.75*µ*s (Fig.S1(b)).

#### C. MicroROSE RNA Thermometer

The following protocol is used to maximize the diversity of the initial seeds selected for all-atom, explicit solvent MD simulations. First, the **G**-vector representations[61] of all FARFAR2 assembled tertiary structures are computed and projected onto nP C = 2, 3, …, 10 principal components (PCs). For each number of PCs, the projected data is clustered using advanced density peaks clustering (ADP), as implemented in dadapy[62]. The only parameter for ADP is Z, or the merging factor, which is set to 1. Z has a default value of 1.65 and determines how the aggressively peaks are merged: a higher Z results in more merging/fewer clusters, while a lower Z results in less merging/more clusters. To ensure maximal conformational diversity in the initial MD ensemble, the number of PCs which yields the largest number of clusters via ADP is used to select seeds. For this analysis, nP C = 8 yields the largest number of clusters, which was 11. The clusters are then ranked based on the average Rosetta score over all conformations assigned to that cluster. To assemble the initial seeds, the clusters are accessed in reverse order of their average Rosetta scores and the top scoring conformer from each cluster is selected as a seed for MD and removed. The clusters are accessed in a round robin until 50 initial seed structures are assembled.

Unbiased molecular dynamics simulations are initiated from these 50 initial seed structures. The MicroROSE RNA thermometer is solvated in a rhombic dodecahedral periodic box with a vector length of 8.03 nm, which is chosen to ensure that even the most extended RNA conformations remain at least 1.2 nm from the box boundaries. To neutralize the negatively charged RNA, Na^+^ ions are added, and additional Cl*^−^* ions are included to achieve a NaCl concentration of 0.1 M. Approximately 14,000 TIP4P-D water molecules are added, resulting in a system comprising 43, 000 atoms in total (Fig.S1(c)). The most recent DESRES AMBER 2022 force field[54] is used to model the RNA, in combination with the CHARMM22 ion parameters[55]. We use the same equilibrium protocol described above to prepare the simulations. All simulation details, (*e.g.*, thermostat, interaction cutoffs, integration time step, etc.) are kept the same as the HIV-TAR system.

The initial 50 seeds are simulated at 310 K for an average of 478 ns, and at 450 K for an average of 256 ns. An adaptive seeding protocol similar to the protocol used in the HIV-TAR system is followed. First, a LaTF[24] model is trained to learn a two-dimensional information bottleneck which integrates information from both the high-temperature and low-temperature simulations. To generate new seeds for the 310 K simulations, samples are generated by LaTF at 310 K and clustered using the k-means (k=50) and k-centers (k=50) algorithms to yield 100 cluster centers. K-means is used to generate seed structures that approximately follow the equilibrium distribution, while k-centers is used to ensure exploration of boundary and lower probability regions of the conformational landscape. The cluster centers are reconstructed as all-atom structures by mapping each center to the nearest simulation data-point in the information bottleneck space. To generate new seeds for the 450 K simulations, samples are generated by LaTF at 450 K and clustered using only k-centers (k=25) to focus the high-temperature simulations on exploration of the conformational landscape. Each round of this LaTF-based adaptive seeding procedure generates 100 seeds to be simulated at 310 K and 25 seeds to be simulated at 450 K. Re-seeding is repeated for 3 rounds to yield a total of 350 trajectories at 310 K with average length of ~ 0.88*µ*s, and 125 trajectories at 450 K with an average length of ~ 0.62*µ*s. Our final dataset cumulatively contains ~ 308*µ*s of simulation time at 310 K and ~ 77*µ*s of simulation time at 450 K (Fig.S1(c)).

### V. TRAINING OF LATENT THERMODYNAMIC FLOWS

#### A. Featurization & Generation of Initial Labels

Two types of internal distances are employed to construct structural descriptors for the RNA molecules, i.e., the pairwise distances between ribose 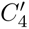 carbon atoms and the norm values of **r**-vectors[61]. The **r**-vectors are defined by introducing a local coordinate frame centered at the geometric center of atoms C2, C4, and C6 of very nucleobase. The x-axis is oriented toward atom C2, while the y-axis points toward atom C4 for pyrimidines (C/U) or atom C6 for purines (A/G) (see the illustration of the coordinate frame in Fig. S8(a)). Within this reference coordinate framework, the position of another nucleotide j relative to base i can be represented by the inter-nucleobase **r**-vector **r** = *x_ij_*, *y_ij_*, *z_ij_* . Distance information characterized by the norms of the **r**-vectors has been shown to effectively distinguish distinct RNA interaction types, including base stacking as well as Watson-Crick and non-Watson-Crick base pairing[61]. Therefore, we rely on **r**-vectors in this study to represent the complex interaction patterns underlying diverse RNA conformations. Each GCAA tetraloop RNA structure is encoded using a 180-dimensional descriptor, while the HIV-TAR RNA stemloop and the MicroROSE RNA thermometer are represented by 1624-dimensional descriptors, respectively. The quantification of **r**-vectors is conducted using the Python package *Barnaba*[63].

To train the LaTF models, initial state labels must first be assigned to each RNA conformation. Following our previous work[64], we generate fine-grained labels by combining time-lagged independent component analysis[65] (tICA, with kinetic mapping[66]) and k-means clustering. The initial labeling procedure involves three key hyper-parameters: the number of tICs, the tICA lag time, and the number of k-means clusters. To balance systematic and statistical errors, we select the optimal hyperparameter combination based on the generalized matrix Rayleigh quotient (GMRQ) score evaluated via cross-validation. The input features for tICA are the structural descriptors described above and the cross-validation is performed by partitioning the full trajectory ensemble uniformly into five folds. Four folds are used for Markov state model (MSM) construction and the remaining fold is reserved for validating the model’s ability to capture the dominant slow dynamical modes. Cross-validation is repeated over five independent trials, each using a different random partitioning of the dataset. Three hyperparameters are screened and optimized in an sequential and iterative manner, whereby two parameters are held fixed while the third is varied until convergence to the optimal combination is achieved. The optimal parameter set is chosen such that the training score converges and the validation score shows minimal deviation from the training score. The whole protocol is performed exclusively using the low-temperature simulation data for each system, and the results are presented in Fig.S5(a), S12(a) and S23(a). The summary of optimal hyperparameter choices for each system are reported in Table S6.

**Table S6:** Summary of Optimal Hyperparameters for Initial Label Generation.

| Systems | # of tICs | tICA Lag Time | # of k-means Clusters |
| --- | --- | --- | --- |
| GCAA Tetraloop | 4 | 50 ns | 100 |
| HIV-TAR RNA Stemloop | 2 | 100 ns | 300 |
| MicroROSE RNA Thermometer | 6 | 100 ns | 50 |

#### B. LaTF Model Architecture & Training Protocol

The LaTF model unifies representation learning and generative modeling within a single framework[24], and can both (1) construct low-dimensional descriptors that extract the dominant slow conformational dynamics from high-dimensional data, and (2) generate temperature-aware structural ensembles. Such unification of representation learning and generative modeling demonstrates two key capabilities. First, representation learning and generative modeling mutually reinforce one another: the flow-based generative model improves the alignment between the encoded posterior distribution and an arbitrary prior distribution, whereas the learned low-dimensional representation enhances generative accuracy while reducing computational cost. Second, through the incorporation of a flexible, temperature-steerable tilted Gaussian prior, LaTF enables physically reasonable interpolation of temperature-dependent equilibrium distributions from limited training data. In particular, LaTF has been shown to infer the melting curve of the GCAA tetraloop in qualitative agreement with experimental observations and extensive simulation results, despite being trained using data from only two temperatures[24].

Here, we employ LaTF for both adaptive sampling and final dynamical model construction. Leveraging its capability to infer temperature-dependent equilibrium distributions, LaTF is trained on-the-fly during data collection to generate seed structures for simulation reseeding, thereby promoting efficient exploration of the energy landscape while improving convergence of equilibrium sampling. After construction of the final dataset, LaTF is retrained exclusively using the low-temperature data to obtain a two-dimensional representation of the RNA conformational landscape and define the corresponding metastable states, which subsequently serve as the foundation for Markov state modeling.

The LaTF architecture comprises three main components: an encoder, a normalizing flow (NF) module, and a decoder. To reduce the computational cost associated with NF training, the LaTF model is optimized in two stages. In the first stage, the parameters of the NF module are frozen, and only the encoder and decoder are trained. This step promotes the merging of the initially labeled short-lived states into kinetically meaningful metastable states while establishing a physically reasonable latent representation. The training procedure follows that of the State Predictive Information Bottleneck (SPIB) and is based on the following loss function:

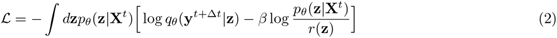

where the latent posterior distribution is modeled via a Gaussian encoder, *p_θ_*(**z X***^t^*), while the future-state label after lag time Δ*t* is reconstructed through a decoder with a softmax output layer, log *q_θ_*(**y***^t^*^+Δ*t*^ **z**). The prior distribution, *r*(**z**) is chosen as a variational mixture-of-posteriors prior (VampPrior). The objective function is designed to extract informative structural features for accurate prediction of future-state labels, while simultaneously regularizing the latent space toward a multimodal Gaussian distribution. In particular, the training is conducted in an iterative and self-consistent manner. Once the loss function converges below a predefined threshold (i.e., convergence threshold) for a specified number of training epochs (i.e., convergence patience), each input conformation is relabeled according to its most probable future-state assignment and states with limited populations are discarded. This refinement procedure is then repeated for a given number of iterations (i.e., refinement number). The resulting model is expected to identify metastable states with lifetimes much longer than the lag time Δ*t*.

After completion of the first training stage, the NF module is incorporated and the entire model is optimized jointly. In this second stage, the encoded latent distribution undergoes an additional flexible bijective transformation before being regularized toward the prior distribution. Improved alignment between the posterior and prior distributions via the generative model enhances both the quality of the learned representation and the refinement of metastable-state boundaries. Specifically, the following loss function is employed for the unified training procedure:

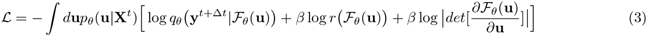

where *F_θ_* denotes the forward transformation of the NF module, and **u** represents the latent random variable generated by the Gaussian encode 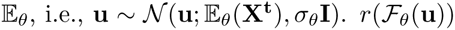 is the prior distribution, which, owing to the flexibility of the NF module, is not restricted to a Gaussian form as in SPIB and can instead adopt an arbitrary distribution. Here, we choose the prior to be a tilted Gaussian distribution of the form:

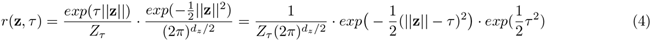

here, 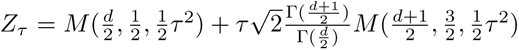 is the normalization constant, where *M* denotes the Kummer confluent hypergeometric function, and d*_z_* is the dimensionality of the latent variable **z**. A standard Gaussian is not chosen as the prior since it concentrates its maximum probability density at a single point, which makes it difficult for the NF model to distinguish multimodal distributions from subtle variations in the latent space. However, the tilted Gaussian distribution distributes its maximum probability density over a hyperspherical shell with radius ||**z||** = *τ* , allowing improved accommodation and separation of multimodal data in the latent space. Furthermore, by introducing the temperature-steerable parameter *T* into the tilted Gaussian prior, as defined below:

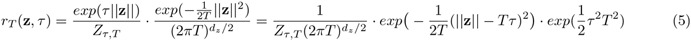

both the radius corresponding to the maximum probability density and the variance of the distribution become proportional to the temperature parameter *T* . Here, 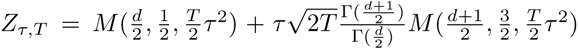 is the temperature-dependent normalization constant. Such a formulation enables the prior distribution to capture entropic effects that reshape and shift free-energy basins across temperatures, facilitating the inference of temperature-dependent latent distributions using training data collected at only a limited number of temperatures. Setting the tilting factor *τ* = 0 reduces the tilted Gaussian prior to a standard Gaussian distribution, where only the variance remains temperature dependent.

#### C. Training Hyperparameter

We perform cross-validation to determine the optimal LaTF training hyperparameters, specifically the learning rate, the *β* weight in the loss function, and the convergence threshold. We employ the GMRQ score as the evaluation metric in the same manner as the protocol used for selecting the initial labeling parameters. The MD trajectories are partitioned into five folds, of which four are used to train a SPIB model (without NF module) and construct an MSM based on the identified metastable states, while the remaining one fold serves as the validation set. We employ SPIB for hyperparameter optimization because of its lower training cost and its role as the baseline for the second-stage LaTF training. Furthermore, the introduction of the NF module does not significantly alter the metastable state definitions. The three hyperparameters are optimized iteratively and sequentially, i.e., two parameters are fixed while the third is tuned to achieve a converged training score and the smallest discrepancy between the training and validation scores, until overall convergence is reached. For each RNA system, cross-validation is performed using different data partitions from five different random seeds to ensure that the resulting model captures the dominant slow dynamical processes without overfitting the limited MD dataset. In parallel, the tilting factor of the prior distribution is screened and optimized to minimize generative deviations, as quantified by the KL divergence. The parameter-selection results are presented in Fig. S5(b), S12(b) & S23(b), and all hyperparameters used for LaTF training are summarized in Table S7.

**Table S7:** Neural network architectures and training hyperparameters used for training LaTF models.

| System | Latent Thermodynamic Flows Parameters |  |  |  |  |  |  |  |  |  |
| --- | --- | --- | --- | --- | --- | --- | --- | --- | --- | --- |
| | Learning Rate | $\beta$ Weight | Enc-Dec Neurons | Coupling Layers | $\tau$ Value | Flow Neurons | Batch Size | Convergence Threshold | Convergence Patience | Refinement Number |
| GCAA Tetraloop (1st stage) | $10^{-4}$ | $5 \times 10^{-4}$ | 64/64 | N/A | N/A | N/A | 1024 | 0.005 | 5 | 15 |
| GCAA Tetraloop (2nd stage) | $10^{-4}$ | $10^{-3}$ | 64/64 | 50 | 2 | 16 | 1024 | 0.01 | 250 | 1 |
| HIV-TAR RNA Stemloop (1st stage) | $5 \times 10^{-4}$ | $10^{-3}$ | 512/128 | N/A | N/A | N/A | 1024 | 0.002 | 5 | 15 |
| HIV-TAR RNA Stemloop (2nd stage) | $5 \times 10^{-4}$ | $10^{-3}$ | 512/128 | 50 | 4 | 16 | 1024 | 0.01 | 250 | 1 |
| MicroRose RNA Thermometer (1st stage) | $5 \times 10^{-4}$ | $5 \times 10^{-4}$ | 512/128 | N/A | N/A | N/A | 1024 | 0.001 | 5 | 15 |
| MicroRose RNA Thermometer (2nd stage) | $5 \times 10^{-4}$ | $5 \times 10^{-4}$ | 512/128 | 50 | 4 | 16 | 1024 | 0.01 | 250 | 1 |

### VI. MARKOV STATE MODELING & EXPERIMENTAL VALIDATION

#### A. GCAA Tetraloop

##### Markov State Models Construction & Validation

Self-supervised training of the LaTF model on the 300 K dataset resolves 20 metastable states at a lag time of 100 ns. Each MD conformation is assigned to the most probable state as inferred by the trained encoder–decoder. The state definitions obtained at 300 K are subsequently transferred to the 400 K simulations, enabling a consistent comparison of state-resolved thermodynamic and kinetic properties across temperatures. The transition dynamics among these metastable states are modeled independently at each temperature using a Markov process. While the RNA energy landscape is often described as ‘glassy’ at low temperatures, we first examine whether the unbiased MD simulations adequately sample reversible transitions. By directly counting transition events between each pair of metastable states at a lag time of 300 ns, we obtain a nearly symmetric transition count matrix (TCM) for 300K, as shown in Fig. S6(a), indicating that our extensive adaptive sampling protocol successfully captures reversible transitions among distinct conformational basins. The eigenvalue spectrum of the corresponding normalized transition probability matrix (TPM) is shown in Fig. S6(b). Lacking clear timescale separation, the continuous spectrum reveals substantial dynamical heterogeneity even for this simple RNA tetraloop. Furthermore, the presence of a single dominant eigenvalue equal to unity indicates the absence of disconnected components in the TPM. At 400 K, we observe the disappearance of many metastable states (Fig. S6(d)), which results in a significantly smoother free-energy landscape and clear separation of dynamical timescales (Fig. S6(e)).

Analysis of the dominant implied timescales (ITS) derived from transition probability matrices constructed at varying lag times shows that the timescales converge beyond 300 ns at 300 K and 150 ns at 400 K, thereby indicating appropriate Markovian lag times for MSM construction (Fig. S6(c,f)). Chapman–Kolmogorov (CK) tests further confirm that MSMs built using a lag time of 300 ns at 300 K and 150 ns at 400 K accurately reproduce long-time dynamics and transition probabilities, in agreement with estimates obtained directly from the raw MD trajectories (Fig. S6(g,h)).

##### Nuclear Overhauser Effect (NOE) Derived Distance Calculation

To validate the accuracy of the force field and computational framework in capturing the tetraloop structural heterogeneity, we evaluate the simulation-derived folded-state structural ensemble with the reference of proton distance constraints obtained from nuclear Overhauser effect (NOE) spectroscopy[67]. Specifically, the trained LaTF model enables identification of the most highly populated native state and extraction of its representative atomistic structures. Based on the modeled folded conformers, the average inter-proton distances can be computed as follows to enable direct comparison with experimental measurements:

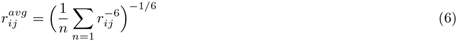

where the r*_ij_* is the pairwise distance and n is the number of folded structures employed for the calculation. An analogous quantification can also be carried out using the refined NMR structural models, as illustrated in Fig. S9.

#### B. HIV-TAR RNA Stemloop

##### Markov State Models Construction & Validation

Following the similar procedure described above for GCAA tetraloop, the LaTF model trained on the 310 K dataset identifies 40 states for the HIV TAR stem–loop using a lag time of 200 ns. The resulting metastable state definition (i.e., the trained encoder and decoder) is then transferred to analyze data obtained at 420 K, enabling the characterization and comparison of temperature-dependent thermodynamic and kinetic properties of the RNA. The raw TCMs are constructed, and the eigenvalues of the corresponding TPMs are quantified to evaluate the connectivity of metastable states sampled by the adaptive and parallel simulations (Fig. S13 (a-b & d-e)). As the ITS become invariant with respect to the lag time beyond 300 ns and 100 ns for the 310 K and 420 K cases, respectively, the corresponding Markovian lag times are selected accordingly (Fig. S13 (c & f)). The resulting MSMs are subsequently validated through CK tests, where the agreement between model-predicted transition probabilities and those directly estimated from the raw trajectory data demonstrates the models’ ability to accurately capture long-timescale dynamics (Fig. S13 (g & h)).

To benchmark the simulated folded ensemble against the NOE-derived distance restraints, we applied the same distance-evaluation protocol described above for the GCAA tetraloop.

##### Residual Dipolar Couplings (RDCs) Calculation

Residual Dipole Coupling (RDC) measurements have emerged as a standard tool for characterizing biomolecular structure and site-specific dynamics. These couplings originate from the direct nuclear dipole-dipole interactions between pairs of magnetic moments. In an isotropic solution, these couplings typically average to zero because the molecules lack a preferential orientation. However, by introducing an anisotropic solvent environment, such as an oriented liquid crystalline phase, as an alignment medium, the molecule’s rotational distribution is slightly perturbed. This induced weak alignment allows for the extraction of RDCs, which provide critical information regarding the orientational distribution of specific bond vectors relative to the molecule-fixed alignment tensor.

In the laboratory coordinate system, the RDC between a nuclear spin pair, i and j, can be quantified as:

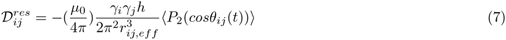

where *r_ij,eff_* represents the effective internuclear distance after accounting for internal vibrational corrections, and *γ_i_* and *γ_j_* denote the gyromagnetic ratios of the respective nuclei. The angular dependence is described by the second-rank Legendre polynomial, *P*_2_(*cosθ_ij_*(*t*)), where angle *θ_ij_* denotes the angle between the internuclear vector and the external magnetic field.

In general, molecular alignment is not tightly restricted under experimental controls, and consequently both RDC data and a set of structural coordinates must be known *a priori* to determine the alignment (order) tensor. For a rigid molecular structure, the time-averaged second Legendre polynomial can be expressed in the molecule-fixed coordinate system as:

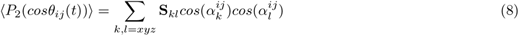

where *α^ij^* denotes the angle subtended between the *ij*th internuclear vector and a given axis of the molecular coordinate system, and **S***_kl_* represents the symmetric and traceless Saupe order tensor, containing five independent elements, that describes the orientation of the magnetic field relative to the molecular coordinate frame. The determination of the alignment (order) tensor **S** therefore requires a set of structural coordinates together with at least five independent experimental RDC measurements. The alignment tensor can then be obtained by solving the resulting system of linear equations using singular value decomposition (SVD) or iterative least-squares minimization. This approach may also be employed to assess the independence of RDC datasets measured under different alignment media. A comprehensive review of RDC techniques can be found in Ref. [68].

Herein, an alternative approach based on a steric obstruction model is employed to predict molecular alignment and RDCs. In this framework, the nematogen (Pf1 bacteriophage) is approximated as an infinite cylinder, and the input RNA structure undergoes 1D translational grid searching together with uniform orientational sampling[69]. Orientations resulting in incompatibility with the liquid crystal environment are excluded, and the overall alignment tensor is subsequently computed as the linear average over all sterically permissible configurations. This algorithm may not achieve the optimal fitting consistency obtained using SVD, but it effectively prevents overfitting. Practically, the program named Prediction of AlignmEnt from Structure (PALES) [70] is employed, and the calculations are performed using the following command:

`pales-linux -stPales -pdb structure.pdb -inD E0_wt_rdc.tab -outD output.txt -H -pf1 -wv 0.022`

where an infinite-cylinder liquid crystal model is adopted with an effective concentration specified by *-pf1 -wv 0.022*. Starting from the input RNA structure, *structure.pdb*, all translational and rotational configurations are systematically explored. Based on the constructed and validated MSM at 310 K, RDCs are computed for conformations assigned to each metastable state, and the corresponding stationary populations are used as weights to obtain ensemble-averaged RDCs. The experimental RDC data, taken from Ref.[16, 71], serve as a reference for comparison with the computational predictions, as shown in Fig. S20(a). We conduct our analysis only with the stem–loop structures directly sampled from MD simulations without including any *in silico* helix elongation using ideal A-form geometry.

##### Spin Relaxation Order Parameter Calculation

The square of Lipari-Szabo generalized order parameter (OP) *S*^2^ [72] is widely employed to quantify the spatial restriction of fast internal motions for internuclear vectors. Specifically, *S*^2^ characterizes the amplitude of orientational fluctuations for individual bond vectors, thus providing a direct link to dynamics derived from NMR spin relaxation measurements. A brief summary of its mathematical definition and physical interpretation is presented here.

The NMR relaxation contribution from the dipolar-dipolar interaction in a nuclear pair can be evaluated via the following correlation function:

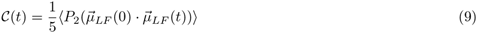

where 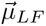 is directed along the internuclear axis between the two nuclei, defined within the laboratory coordinate system and 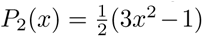 is the second Legendre polynomial. Under the assumption that the global tumbling of the molecule is characterized by a single isotropic correlation time and can be adiabatically decoupled from internal dynamics, the total correlation function can be decomposed into a product of overall and internal components:

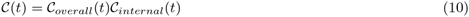

Within a truncated observation window of picoseconds to nanoseconds, the overall correlation function remains approximately unchanged, given that its correlation time is much longer than the that of the internal dynamics. The internal term of the correlation function can therefore be defined by the internuclear vector 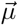within a molecular-fixed reference frame:

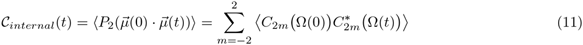

The correlation function is further reformulated using modified spherical harmonics, i.e., 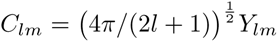, expressed as a function of the polar angles Ω = (*θ, ϕ*) of the internuclear vector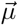. Within this framework, the OP *S*^2^ is defined as the value of the internal correlation function in the limit of infinite time:

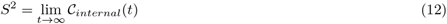

By invoking the following property of the correlation function in the limit of infinite time, *i.e.*, lim*_t→∞_ A*(0)*B*(*t*) = *A B* , the OP *S*^2^ can be reformulated as a summation of the squared ensemble averages of the individual modified spherical harmonic components:

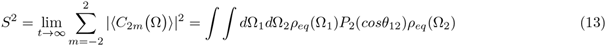

where *θ*_12_ represents the angle between the vectors 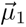 and 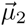, associated with the polar angles Ω_1_ and Ω_2_. The equilibrium distribution of these orientations is described by the density function *ρ_eq_*(Ω).

In practice, the order parameter *S*^2^ can be calculated from simulations and compared to experimental values derived from NMR spin relaxation measurements to evaluate how accurately the simulations capture local orientational fluctuations of bond vectors. In line with prior studies[16, 73], *S*^2^ values can be computed from the three-dimensional coordinates of the ensemble of *in silico* generated structures using the following equation:

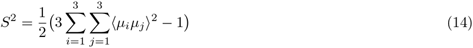

where *µ_i_* and *µ_j_* correspond to the Cartesian components of the normalized internuclear vector. In this study, we examine the relaxation of C8–H8 dipolar interactions for adenine and guanine, and C6–H6 dipolar interactions for cytosine and uracil. To eliminate the overall rational diffusion, all structures from MD simulations are aligned to a reference configuration using backbone atoms before computing the internuclear vectors. Since spin relaxation measurements probe motions on the picosecond-to-nanosecond timescale, the ensemble average in Eq.14 is first evaluated over conformations within each metastable state identified by the LaTF model, respectively. The order parameter is subsequently obtained by averaging across states, weighted by their equilibrium populations obtained from the validated MSM. This procedure relies on certain approximations, as the lifetimes of individual states may vary and can exceed the spin relaxation timescale. The OP values are further normalized to the largest corresponding value among helical residues for each resonance type (C6 and C8) independently. These normalized values are then compared with experimental results reported in previous studies[74**?**]. Results are illustrated in Fig. S20(b).

##### Conformational Heterogeneity Induced by Ligand/Peptide Recognition

**Table S8:**
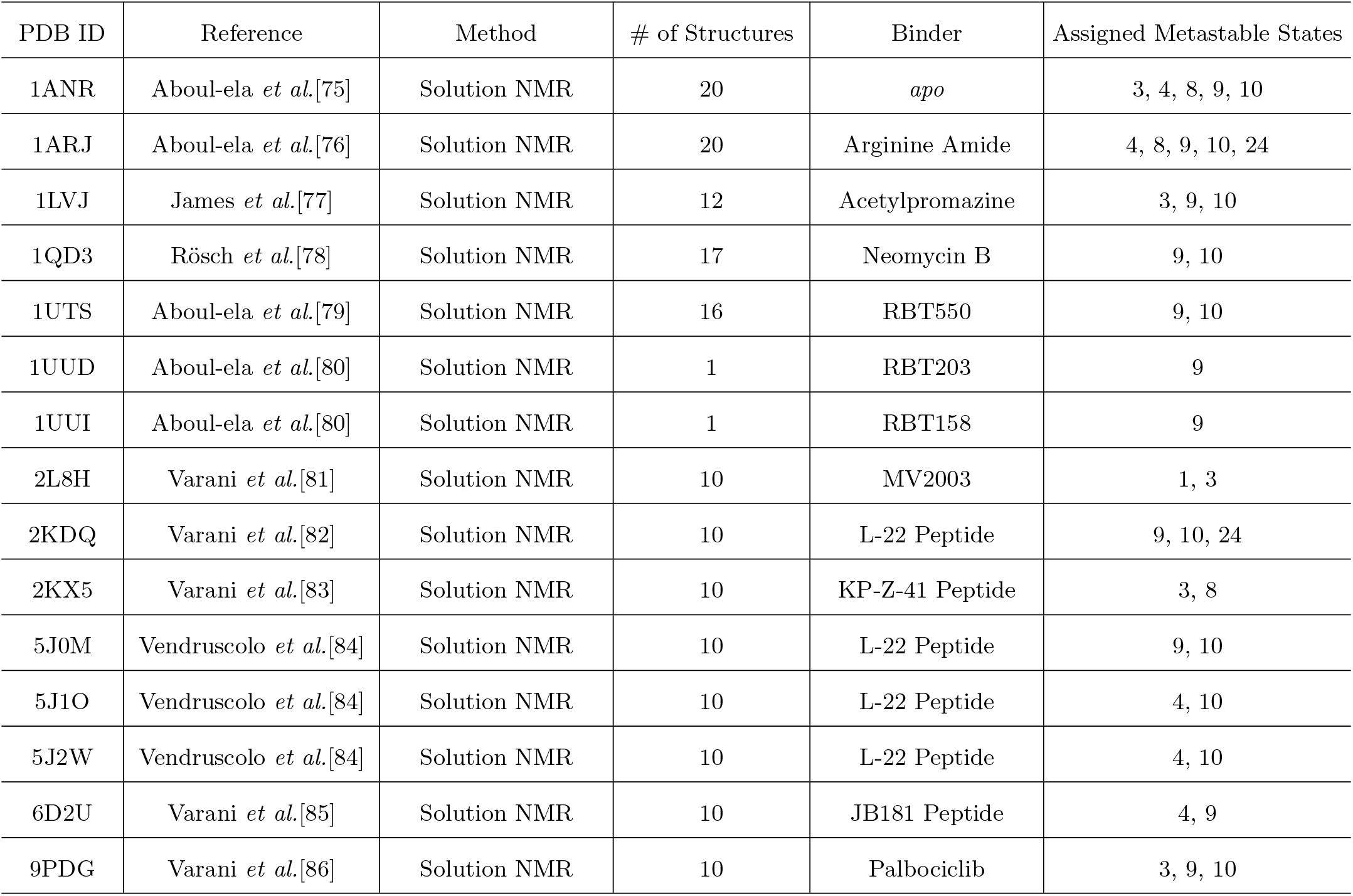
Summary of Experimental HIV TAR Conformations With and Without Binder.

| PDB ID | Reference | Method | # of Structures | Binder | Assigned Metastable States |
| --- | --- | --- | --- | --- | --- |
| 1ANR | Aboul-ela *et al.*[75] | Solution NMR | 20 | *apo* | 3, 4, 8, 9, 10 |
| 1ARJ | Aboul-ela *et al.*[76] | Solution NMR | 20 | Arginine Amide | 4, 8, 9, 10, 24 |
| 1LVJ | James *et al.*[77] | Solution NMR | 12 | Acetylpromazine | 3, 9, 10 |
| 1QD3 | Rösch *et al.*[78] | Solution NMR | 17 | Neomycin B | 9, 10 |
| 1UTS | Aboul-ela *et al.*[79] | Solution NMR | 16 | RBT550 | 9, 10 |
| 1UUD | Aboul-ela *et al.*[80] | Solution NMR | 1 | RBT203 | 9 |
| 1UUI | Aboul-ela *et al.*[80] | Solution NMR | 1 | RBT158 | 9 |
| 2L8H | Varani *et al.*[81] | Solution NMR | 10 | MV2003 | 1, 3 |
| 2KDQ | Varani *et al.*[82] | Solution NMR | 10 | L-22 Peptide | 9, 10, 24 |
| 2KX5 | Varani *et al.*[83] | Solution NMR | 10 | KP-Z-41 Peptide | 3, 8 |
| 5J0M | Vendruscolo *et al.*[84] | Solution NMR | 10 | L-22 Peptide | 9, 10 |
| 5J1O | Vendruscolo *et al.*[84] | Solution NMR | 10 | L-22 Peptide | 4, 10 |
| 5J2W | Vendruscolo *et al.*[84] | Solution NMR | 10 | L-22 Peptide | 4, 10 |
| 6D2U | Varani *et al.*[85] | Solution NMR | 10 | JB181 Peptide | 4, 9 |
| 9PDG | Varani *et al.*[86] | Solution NMR | 10 | Palbociclib | 3, 9, 10 |

Over the years, a diverse range of binders, spanning peptide mimics to small molecules, have been developed to specifically recognize HIV-TAR. Through distinct binding modes, these binary complexes may modulate the conformational preferences of HIV-TAR. Solution NMR studies have resolved the structures of numerous HIV-TAR–binder complexes, providing valuable opportunities to examine whether our MD simulations capture the structural heterogeneity induced by ligand or peptide recognition. Here, we analyze HIV-TAR conformations induced by fourteen distinct binders, as summarized in Table S8, together with the corresponding *apo* structures. Specifically, each experimentally resolved structure is represented using **r**-vector and pairwise-distance features and subsequently projected onto the information bottleneck latent space and assigned to metastable states using the trained LaTF model. As shown in Fig. S21 and Table S8, the binder recognition partially restricts the conformational flexibility of HIV-TAR, while the experimentally resolved conformations collectively span seven distinct metastable states across different binders. Notably, all of these states are captured by the MD simulations and the resulting metastable state model, with states 9 and 10 appearing particularly promising for future binder design. The conformational sub-ensembles sampled within these states, together with their varying degrees of local structural flexibility, may provide a valuable foundation for future small-molecule docking and rational ligand design.

#### C. MicroROSE RNA Thermometer

##### Markov State Models Construction & Validation

Following the procedures described above for the GCAA tetraloop and HIV-TAR stemloop systems, we train a LaTF model using the 310 K simulation data for the MicroROSE RNAT system. The LaTF model is trained with a lag time of 50 ns and identifies 32 metastable states. The resulting metastable state definitions (i.e., the trained encoder and decoder) are used to analyze simulation data obtained at 450 K, thereby enabling direct comparison of thermodynamic and kinetic quantities across both temperatures. The raw TCMs are calculated and the eigenvalues of the corresponding TPMs are quantified to evaluate the connectivity of metastable states visited by our extensive adaptive sampling protocol (Fig. S24 (a-b & d-e)). The 310 K eigenspectrum does not display any clear timescale separation, indicating heterogeneous conformational dynamics. In contrast, the TPM estimated at 450 K exhibits timescale separation as more than half of the metastable states have eigenvalues very close to zero, indicating that they decay very quickly. Both TPMs have a single eigenvalue equal to 1.0 which demonstrates that our estimated MSM is ergodic.

The ITS plateau as a function of lag time beyond 300 ns for the TPMs estimated from 310 K simulation data. For the 450 K case, most of the ITS never become independent of lag time. However, one timescale does begin to plateau at 25ns. From the ITS analysis, 300ns and 25ns are selected for MSM construction at 310 K and 450 K, respectively (Fig. S24 (c & f)). The resulting MSMs are subsequently validated through CK tests, which assess the predictive power of MSMs by investigating the agreement between MSM-derived transition probabilities and those directly estimated from the raw trajectory data (Fig. S24 (g & h)).

##### Nuclear Overhauser Effect (NOE) Derived Distance Calculation

To benchmark the simulated folded ensemble against the NOE-derived distance restraints, we compute the ensemble averaged inter-proton distance for each of the 338 proton pairs identified in the reported NOE restraints using the following power law weighting: 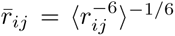. In Fig. S31(a), we compute the correlation between the set of 338 PDB ensemble averaged *r̅_ij_*s and the corresponding set of state 2 ensemble averaged *r̅_ij_*s. This analysis yields an *r*^2^ of 0.67, indicating good agreement between the 500 randomly-sampled state 2 structures and the PDB ensemble obtained via NMR. Fig. S31(b) shows a plot of the first 75 reported NOE restraints, where the experimental uncertainties are visualized as specified in [18]. The S2 generated *r̅_ij_*s lie within the experimental uncertainty for 77.2% (261/338) of restraints. Across 77 violations, the S2 *r̅_ij_*s are on average 0.37^°^A outside the experimental uncertainty. Deviations greater than 1^°^A from experimental uncertainty are only observed for 3 restraints: U88-H6:U87-H4’, U88-H5:U87-H4’, G90-H8:G91-H1’. This result is quite good as 3/4 of the nucleotides involved in large deviations (U87, U88, and G90) are all located in the flexible apical loop. Furthermore, for 42 of the 77 violations, the corresponding PDB averaged *r̅_ij_*s also lie outside the experimental uncertainty, and otherwise appear towards its upper bound.

##### Base-Base Interactions

To compare base-base interactions observed in the NMR structures to those observed via simulation, we calculate displacements 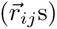 between the two bases under investigation for all structures within the folded sub-ensemble (S2, S10, S16) and extract the *ρ* and *z* components for each 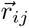; see Fig. S8(a) for a visualization. By examining 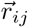 distributions in *ρ*-*z* space, we can conclude whether bases *i* and *j* exhibit preferences for Watson-Crick base pairing, base stacking, or non-canonical interactions. The scatterplots in Figs. S29, S30 are colored based on the Barnaba annotations as described in [63]. ‘WCc’, ‘WCt’ and ‘WWc’ are all colored as Watson-Crick, hence why U-U base pairs have ‘Watson-Crick’ interactions.

Specifically, we investigate the *ρ*-*z* plots of non-canonical pairs within the internal loop (U79-U97, C80-U96, U81-U96), as well as the Watson-Crick pairs flanking it (A78-U98, U82-A95) and the key G83-G94 *syn-anti* base pair. The NMR 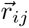 values lie within a single dominant basin for U79-U97 and A78-U98, thereby demonstrating excellent agreement between the folded sub-ensemble and NMR data (Fig. S29 (d)-(e)). In contrast, U82-A95, U81-U96, and C80-U96 all display more flexibility than U79-U97 and A78-U98, as shown by the larger spread in projected NMR 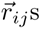 across the 20 conformers and the multiple wells visualized from simulation data (Fig. S29 (a)-(c)). For these three pairs, the interaction patterns observed via NMR are present within our simulation derived folded structures. However, they fall in regions of intermediate free-energy, indicating that these interactions are not dominant within the folded sub-ensemble.

For the microROSE RNAT, G83 is the central guanine found in the universally conserved sequence motif U-U/C-GC-U. When G83 is removed, the microROSE element forms a stable hairpin, losing its internal loop and consequently the ability to unfold in response to temperature changes. Although G83 was originally predicted to exist as a bulged nucleotide, the authors in [18] observe a strong NOE between the G83 H1’ and H8 protons, in addition to G94 chemical shifts indicating that G83 and G94 actually form a *syn-anti* base pair. To investigate whether our simulations recapitulate this key interaction, we inspect the *χ* torsion angle distributions for all 29 microROSE nucleotides (Fig. S30(a)). From this analysis, we observe that only G83 and G90 non-negligibly populate the *syn* conformation (90° *< χ <* 90°): G83 exists in a *syn* conformation for ~13% of structures in the folded sub-ensemble, while G90, an apical loop nucleotide, exists in a *syn* conformation for ~15% of structures over the same ensemble. In Fig. S30(b), we see that over the entire folded sub-ensemble, G83 and G94 most often interact via base stacking. However, by examining only the structures in which G83 exists in the *syn* conformation, we can see that the G83-G94 *syn-anti* base pair is indeed sampled, although not as frequently as G83-G94 stacking interactions.

**Figure S1:**
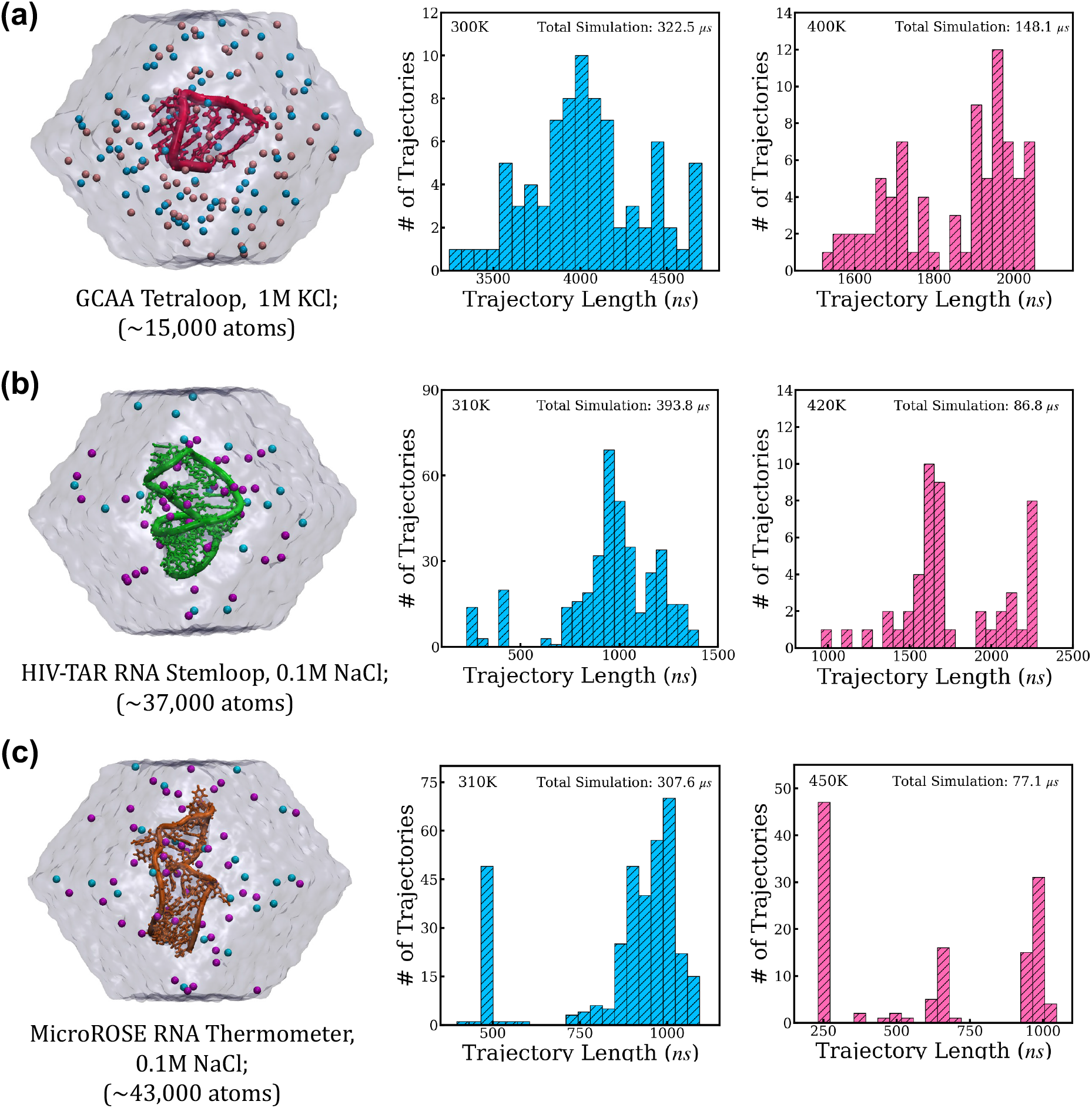
Visualization of the simulated RNA systems together with the distributions of molecular dynamics trajectory lengths obtained at different temperatures. The first column depicts the dodecahedral simulation boxes constructed for the three RNA systems: (a) the GCAA tetraloop, (b) the HIV-TAR stem–loop, and (c) the MicroROSE RNA thermometer. Potassium ions (K^+^) are shown in pink, sodium ions (Na^+^) in magenta, and chloride ions (Cl*^−^*) in cyan. The second and third columns summarize the distributions of MD trajectory lengths obtained at low and high temperatures, respectively. For all three systems, the cumulative simulation time at low temperature exceeds 300 *µ*s.

**Figure S2:**
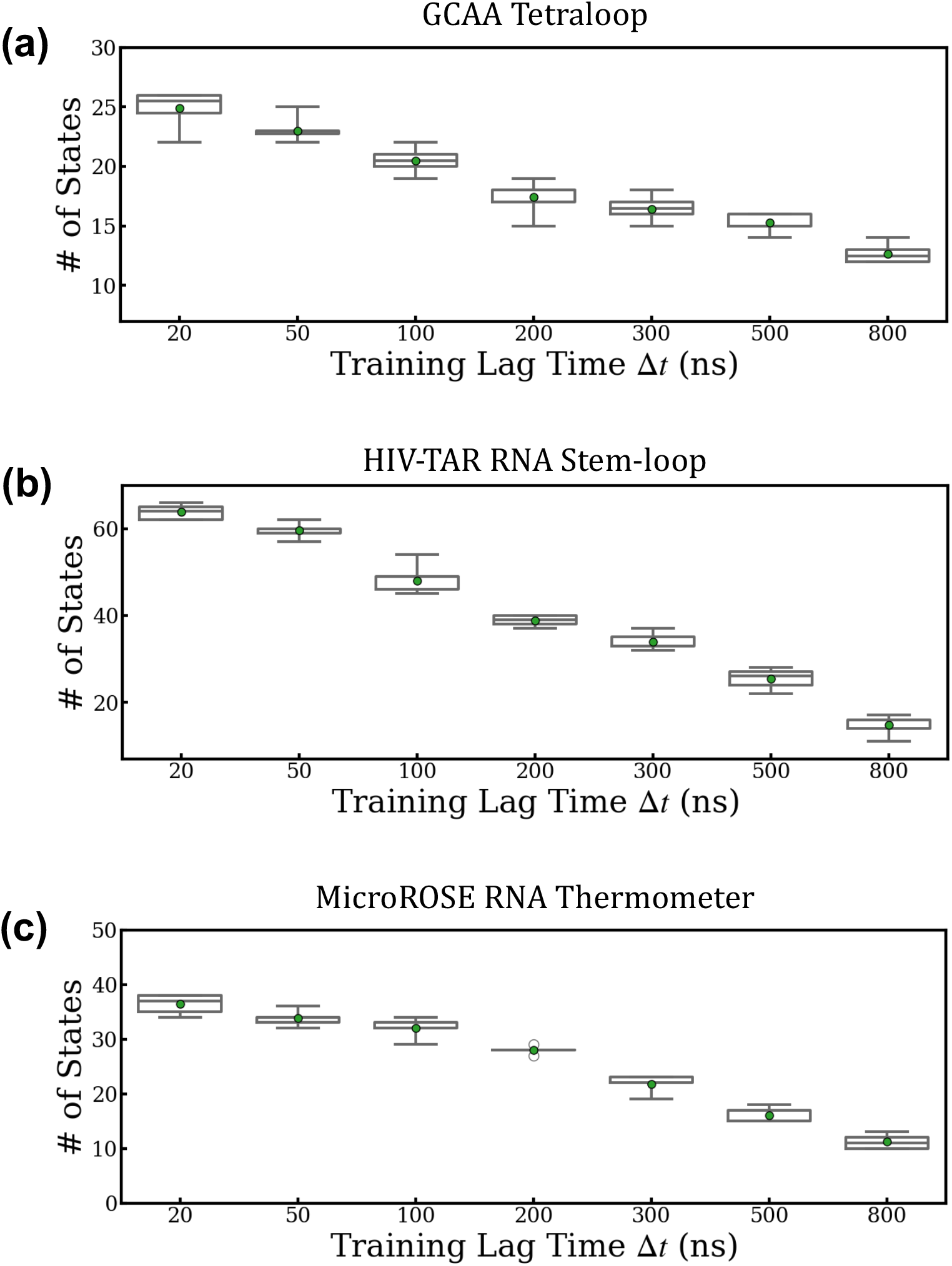
Dependence of the number of resulting metastable states on the lag time Δ*t* of the state predictive information bottleneck (SPIB) model for three RNA systems. Seven lag times, ranging from 20 ns to 800 ns, are used to train SPIB models with r-vectors and pairwise-distance structural descriptors as inputs. For each lag time, ten independent models are trained with different random initializations. The resulting numbers of identified metastable states are reported. The SPIB results are used as a computationally efficient approximation to those of LaTF, as the incorporation of the normalizing flow model typically does not substantially alter the state boundaries.

**Figure S3:**
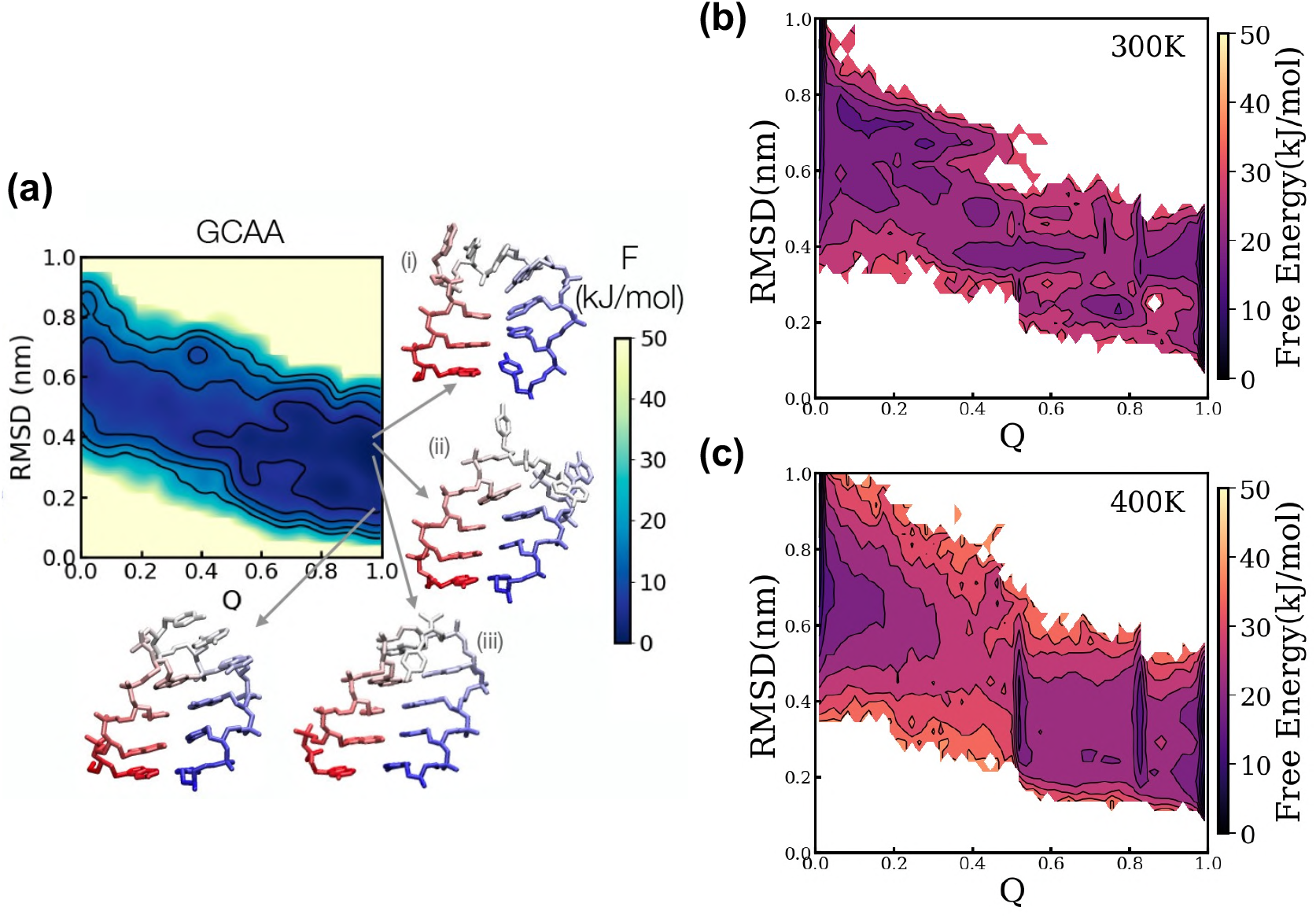
Comparison of the free energy surfaces (FES) of the GCAA RNA tetraloop projected onto two order parameters, obtained using parallel tempering in the well-tempered ensemble combined with well-tempered metadynamics (PTWTE-WTM) and adaptive unbiased MD simulations. (a) Results from previous work by Debenedetti *et al.*[8] are shown, in which the DESRES AMBER 2017 force field[53] is combined with PTWTE-WTM to map the FES of the GCAA tetraloop as a function of the root-mean-square deviation (RMSD) and the order parameter Q at 300 K. The order parameter Q quantifies the folding compactness of the stem region based on distances between contacting heavy atoms. Representative structures are extracted from the major free energy basins and are indicated by arrows. The figure is reprinted from Reference [8] @ 2021 American Chemical Society. (b-c) The FES of the same tetraloop obtained in this work using the same DESRES AMBER 2017 force field[53] are compared. These surfaces are directly estimated from projected raw unbiased MD data at 300 K and 400 K, respectively.

**Figure S4:**
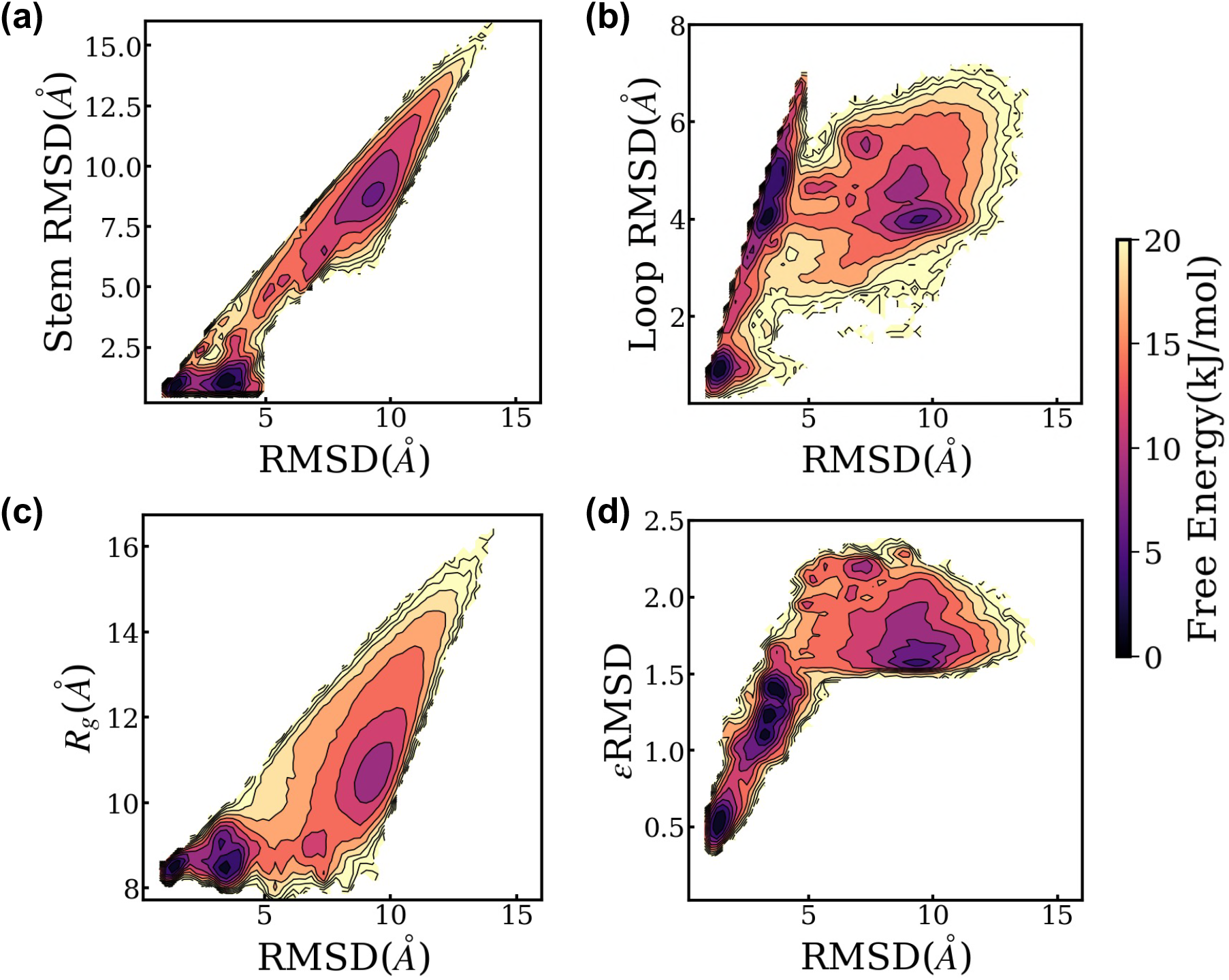
Visualization of unbiased MD sampling of the GCAA tetraloop at 300 K projected onto different structural order parameters. The heavy-atom root-mean-square deviation (RMSD) relative to the native NMR structure (PDB ID: 1ZIH) is computed for each MD-sampled conformation, separately for the entire tetraloop, the stem region, and the loop region, and the corresponding raw distributions from simulation are shown in panels (a–b). In addition, the radius of gyration *R_g_* and the *ɛ*RMSD[61] are evaluated, and their joint raw distributions together with the whole structural RMSD are mapped.

**Figure S5:**
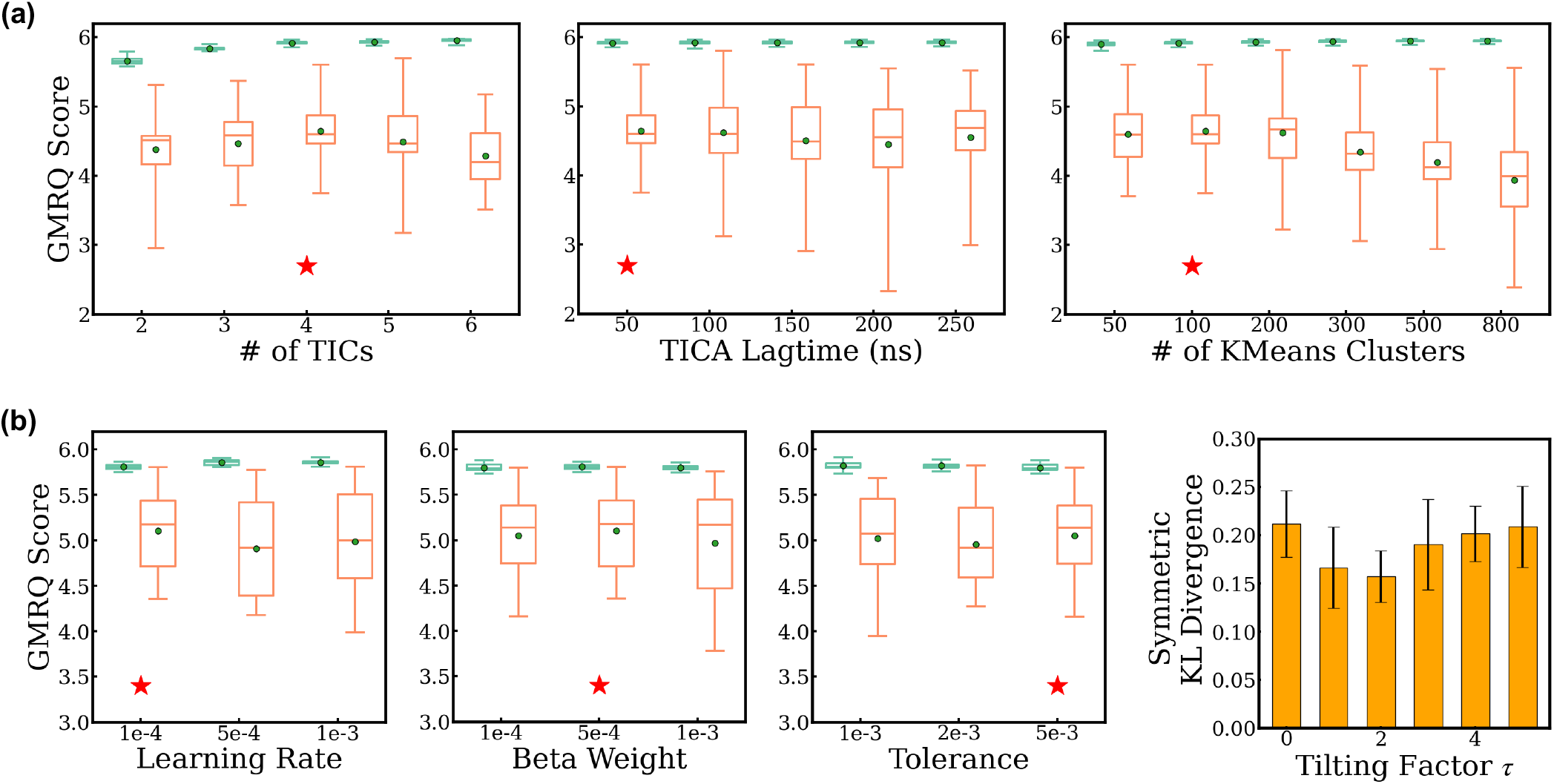
Selection of optimal hyperparameters for training the Latent Thermodynamic Flows model on the GCAA RNA tetraloop system at 300K. (a) Choice of hyperparameters for initial labeling via cross-validation based on the GMRQ score. The tICA method followed by k-means clustering is applied to generate initial labels for LaTF training. Three key hyperparameters, the number of tICs, the tICA lag time, and the number of clusters, are systematically varied to construct Markov state models, whose ability to capture the five dominant slowest dynamical modes is evaluated using the GMRQ score. The cross-validation procedure is performed sequentially and iteratively, whereby two parameters are fixed at their optimal values while the third is scanned, until all three parameters simultaneously yield optimal GMRQ scores. The hyperparameter set that achieves the highest validation score with the smallest discrepancy between training and validation performance is selected as optimal (indicated by red stars). The whole trajectory ensemble is randomly partitioned into five equal folds for cross-validation, with four folds used for training and one for validation. (b) Cross-validation using the same protocol is performed to select three LaTF model hyperparameters: the learning rate, the *β* weight, and the tolerance. Markov state models used for GMRQ score evaluation are constructed based on state definitions derived from the LaTF encoder-decoder assignment. The optimal tilting factor *τ* for the tilted Gaussian prior is chosen as 2 to minimize the symmetric KL divergence between the generated latent distribution and that derived from the MD simulations. Uncertainties are estimated by training the LaTF model five times with different random initializations.

**Figure S6:**
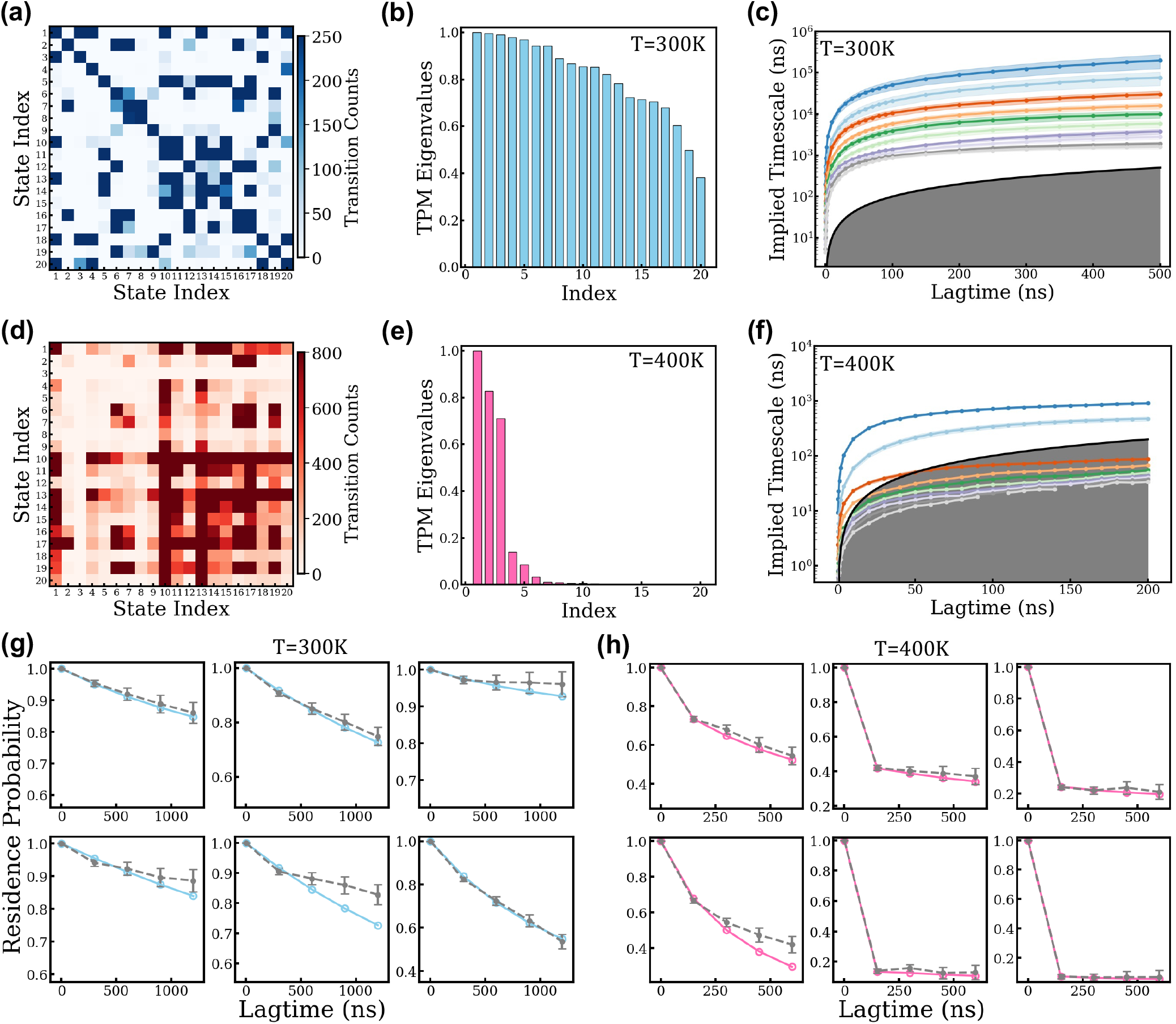
Markov State Models (MSMs) construction, characterization, and validation for the GCAA RNA tetraloop at 300 K and 400K. The Latent Thermodynamic Flows (LaTF) model is trained on molecular dynamics (MD) data collected at 300 K with a lag time of 100 ns. Metastable state definitions are constructed using the trained encoder and decoder, and MD data at 400 K are analyzed using the same state definitions. (a) Raw unsymmetrized transition count matrix (TCM) directly estimated from simulation data at 300 K with a lag time of 300 ns. (b) Ordered eigenvalues of the transition probability matrix (TPM) at 300 K for a lag time of 300 ns. (c) The ten leading implied timescales as a function of transition lag time at 300 K. (d–f) Corresponding analyses for simulations performed at 400 K. The lag times for the TCM and TPM are set to 150 ns. (g) Chapman–Kolmogorov (CK) test of the MSM built with a lag time of 300 ns for the six most populated metastable states at 300 K. (h) Corresponding CK test for the MSM constructed with a lag time of 150 ns at 400 K. The gray error bars denote residence probabilities estimated directly from the raw MD simulation data, whereas the blue and red markers represent the corresponding probabilities obtained from MSM propagations. Uncertainties in both the implied timescales and CK tests indicate the standard derivations estimated via bootstrap resampling of the trajectory ensemble fifty times with replacement.

**Figure S7:**
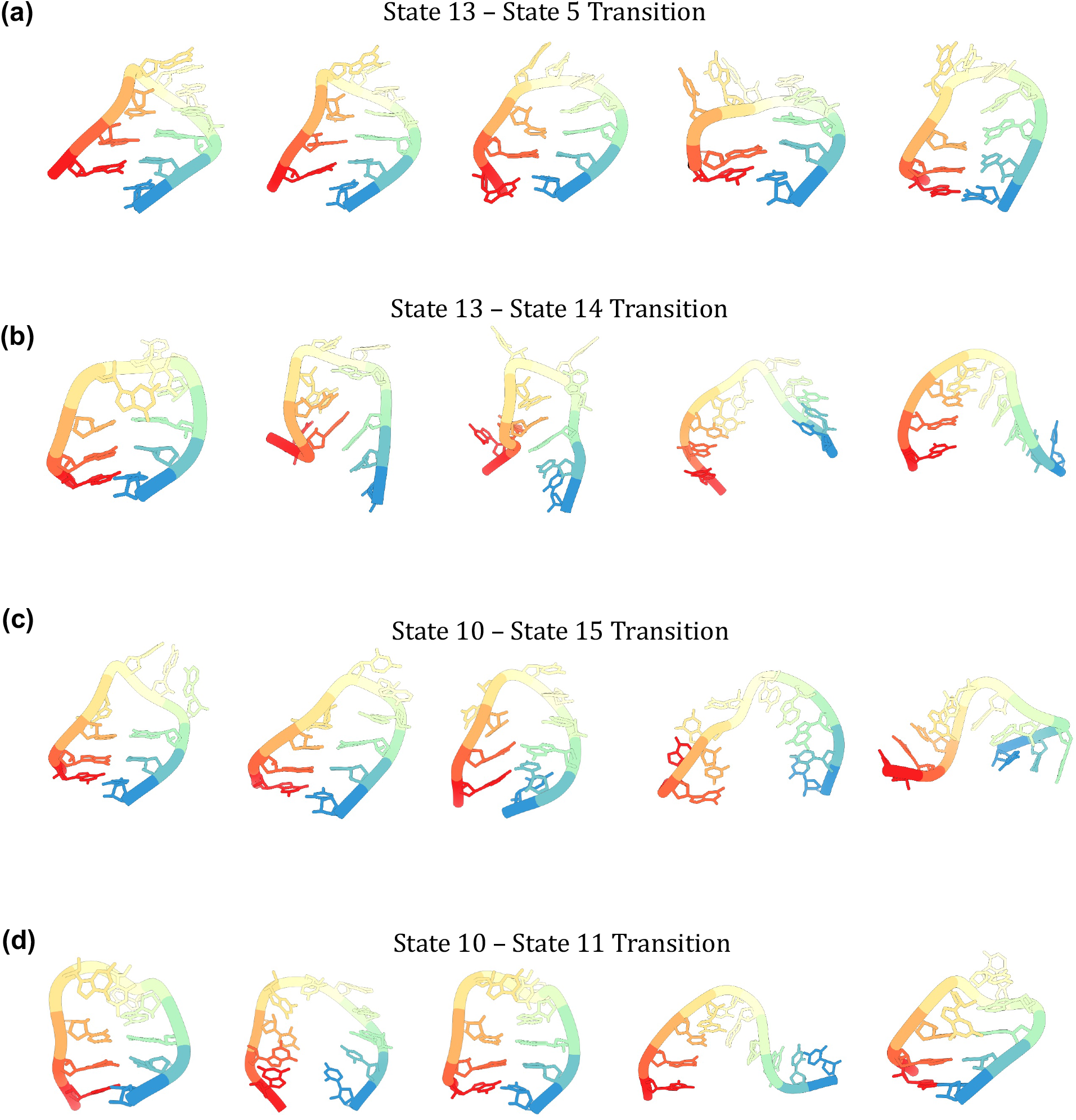
Representative transition-region structures between two selected metastable states in the GCAA tetraloop system. The probabilities of assigning each tetraloop conformation to the metastable states are derived using the trained LaTF decoder model. To identify transition-region structures between states i and j, we compute the assignment entropy s = p*_i_* ln p*_i_* p*_j_* ln p*_j_* and select conformations with the largest entropy values, which correspond to comparable, non-zero assignment probabilities to both states. Along different folding pathways, the top five highest-entropy structures are extracted and visualized between four pairs of metastable states. Structures are displayed in the spectral color scheme, with the 5*^′^* terminus colored red and the 3*^′^* terminus colored

**Figure S8:**
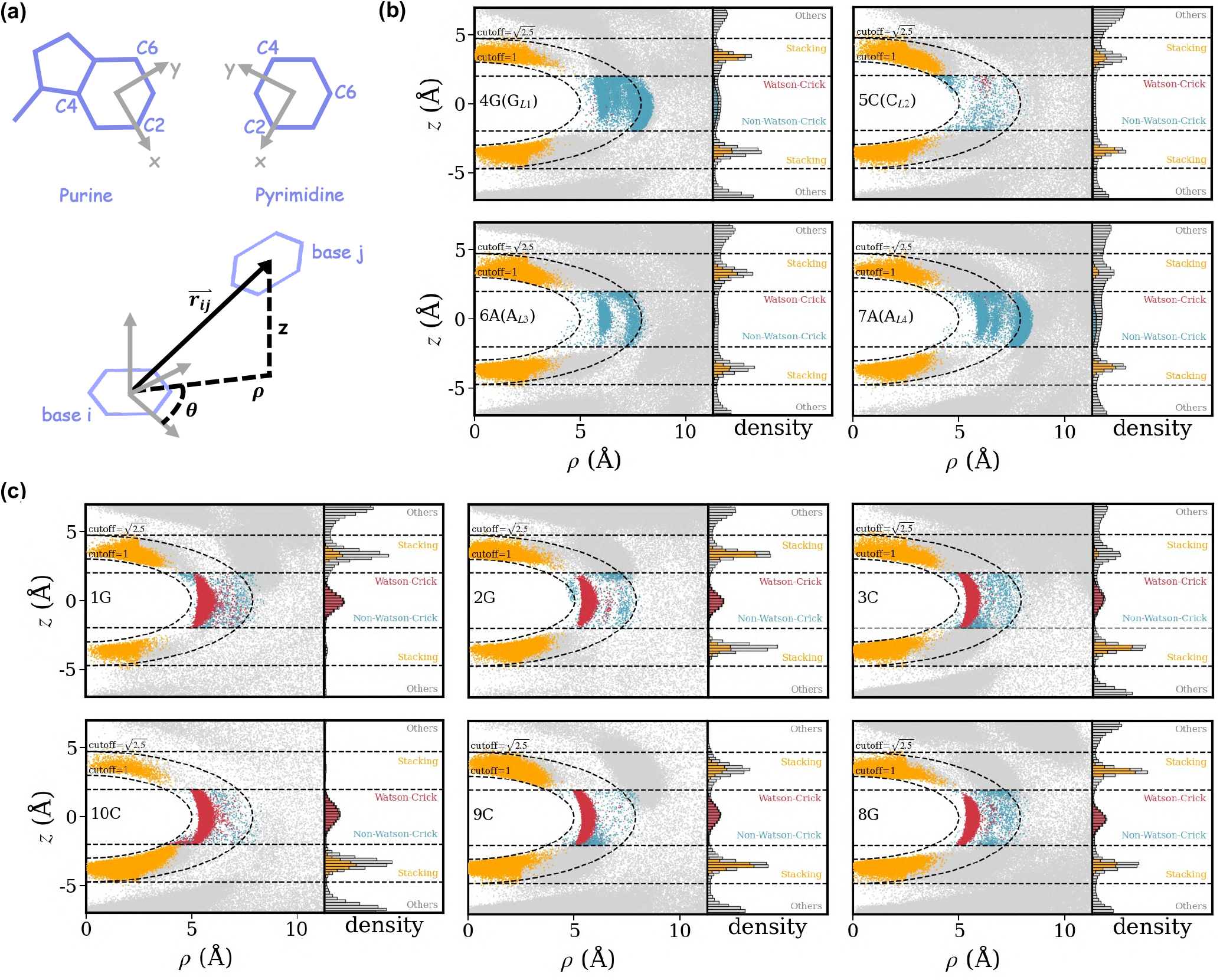
Illustration of the equilibrium local interaction patterns associated with each nucleotide in the GCAA tetraloop at 300K. (a) Schematic definition of the local coordinate systems for purine and pyrimidine bases and the corresponding **r** vector. The **r***_ij_* vector encodes the relative spatial arrangement of base *j* with respect to base *i* and is represented using cylindrical coordinates *ρ*, *θ* and *z*. (b) Equilibrium local interaction pattern distributions associated with each nucleotide in the loop region of the GCAA tetraloop. For each MD-sampled conformation, interactions between a given nucleotide and its neighboring bases are represented by the corresponding **r***_ij_* vectors and their distributions are projected onto the *ρ*-*z* plane. Scatter plots and one-dimensional histograms along the *z* coordinate are constructed by reweighting conformations according to the stationary populations obtained from the validated Markov state model. Histograms are normalized independently for each nucleotide. Interaction types are classified using the criteria implemented in the *Barnaba* package[63] and are color-coded as follows: Watson-Crick base pairing (red), non-Watson-Crick pairing (light blue), base stacking (orange), and other interactions (gray). An ellipsoidal shell defined by anisotropic position vectors with norms of 1.0°Aand 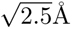[61] is overlaid to highlight regions preferentially occupied by base-pairing and base-stacking interactions. (c) Corresponding equilibrium interaction distributions for the six nucleotides in the stem region of the tetraloop, weighted by the metastable state populations derived from the Markov state model.

**Figure S9:**
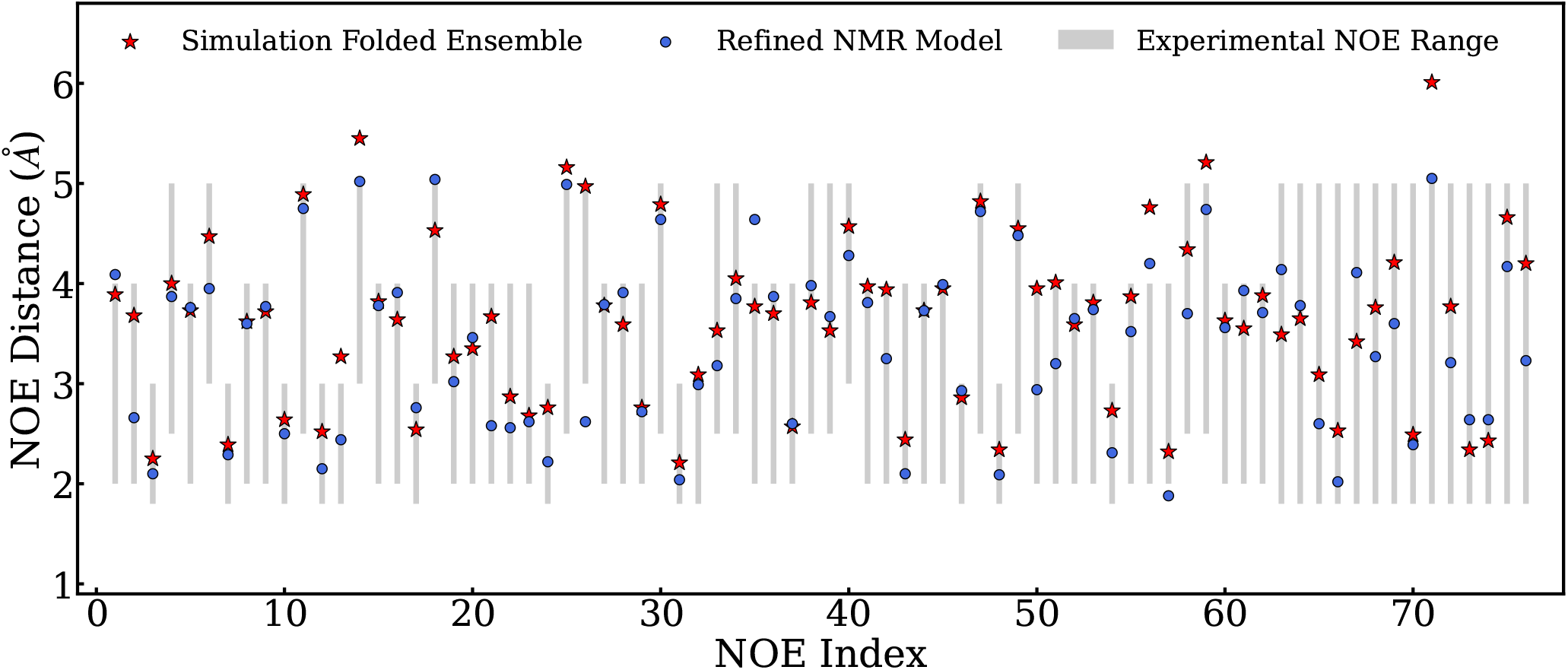
Comparison and validation of the local structural heterogeneity of the GCAA tetraloop in the simulation-derived folded-state ensemble against NMR measurements. The ranges of 76 inter-residue proton-proton distances in the GCAA RNA tetraloop derived from nuclear Overhauser effect (NOE) measurements are shown as gray bars and used as reference values. A Latent Thermodynamic Flows model trained on simulation data at 300 K identifies a dominant metastable state that exhibits a substantially higher population and significantly smaller structural deviation from the native NMR structure than all other states. Five hundred structures are randomly sampled from this state to represent the folded ensemble. The average inter-hydrogen distances quantified from these conformers are reported and indicated by red stars. Corresponding distances calculated from the refined NMR structural models (PDB ID: 1ZIH) are shown as blue circles for comparison.

**Figure S10:**
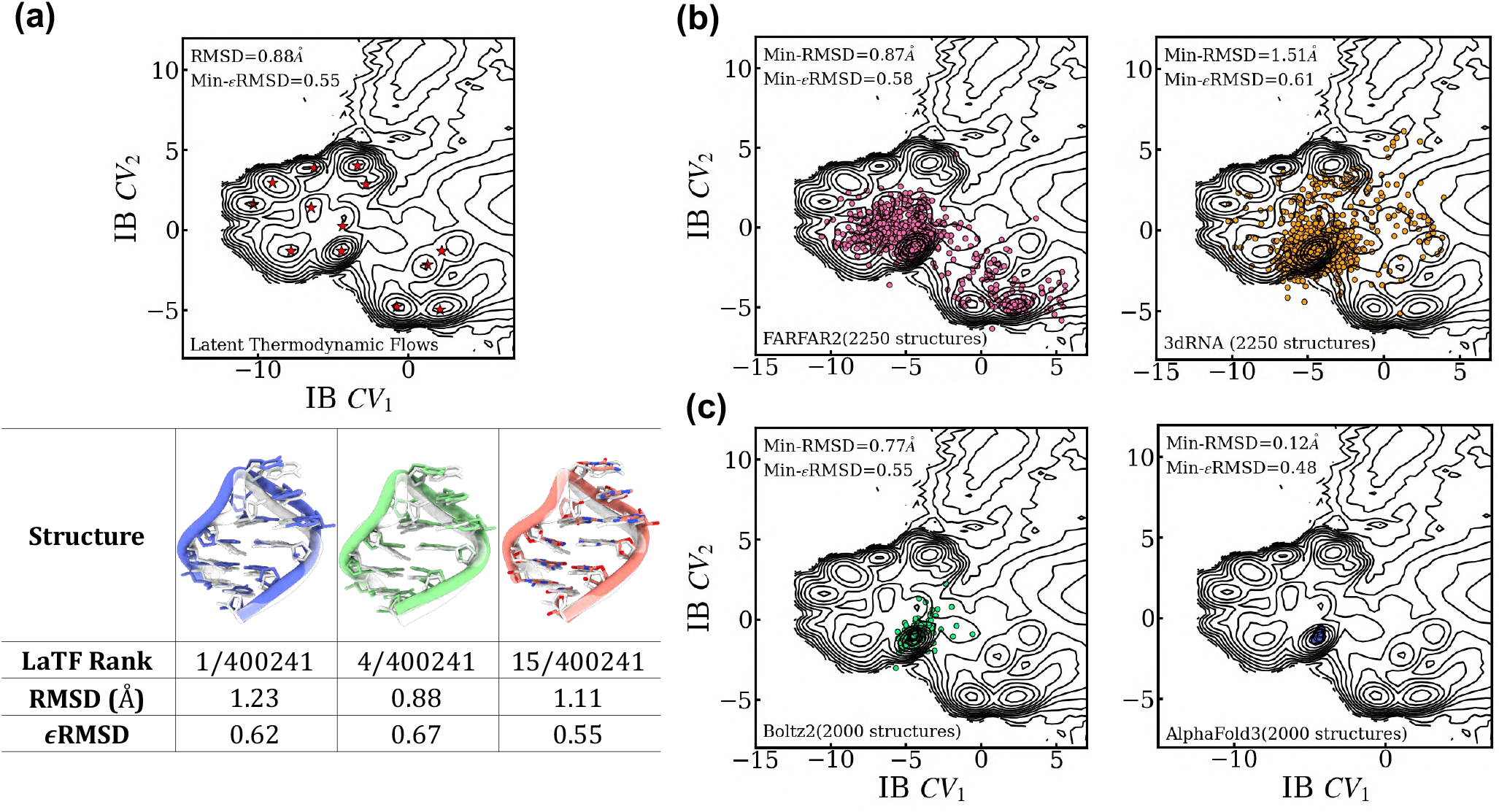
Performance evaluation of tertiary structure modeling approaches in prediction accuracy and structural heterogeneity sampling for the GCAA tetraloop system. (a) The free energy landscape sampled by adaptive molecular dynamics simulations at 300 K and projected onto the latent information bottleneck (IB) space via the trained LaTF encoder is visualized. For each metastable state identified by the LaTF decoder, the top-ranked structure (scored by the LaTF normalizing flow component) is marked by a red star. For the most populated folded state, three representative structures are analyzed: the top-ranked structure, the structure with the smallest RMSD, and the structure with the smallest *ɛ*RMSD (in reference of NMR structure). The corresponding structures, along with their ranks and RMSD and *ɛ*RMSD values, are summarized in the table. MD conformations are shown in color, while the NMR structure (PDB: 1ZIH) is displayed in a transparent representation for comparison. (b) Two thermodynamics-based fragment assembly methods, FARFAR2 and 3dRNA, are applied to predict and sample tertiary structures of the GCAA tetraloop using three different secondary structure models. Structural heterogeneity is explored through Monte Carlo simulations performed at multiple temperatures with a knowledge-based scoring function. The resulting 2, 000 conformations are featurized using the **r**-vectors and pairwise distance descriptors and subsequently embedded into the IB space using the trained LaTF encoder. The minimum RMSD and *ɛ*RMSD values across all sampled structures, computed relative to the NMR structure, are reported. (c) Two deep learning–based methods, i.e., Boltz2 and AlphaFold3, are employed to predict tetraloop tertiary structures directly from sequence information. Structural diversity is explored by varying random seeds or diffusion step scales within the generative model. The minimum RMSD and *ɛ*RMSD values across all sampled 2,000 structures with reference of the NMR structure are also reported.

**Figure S11:**
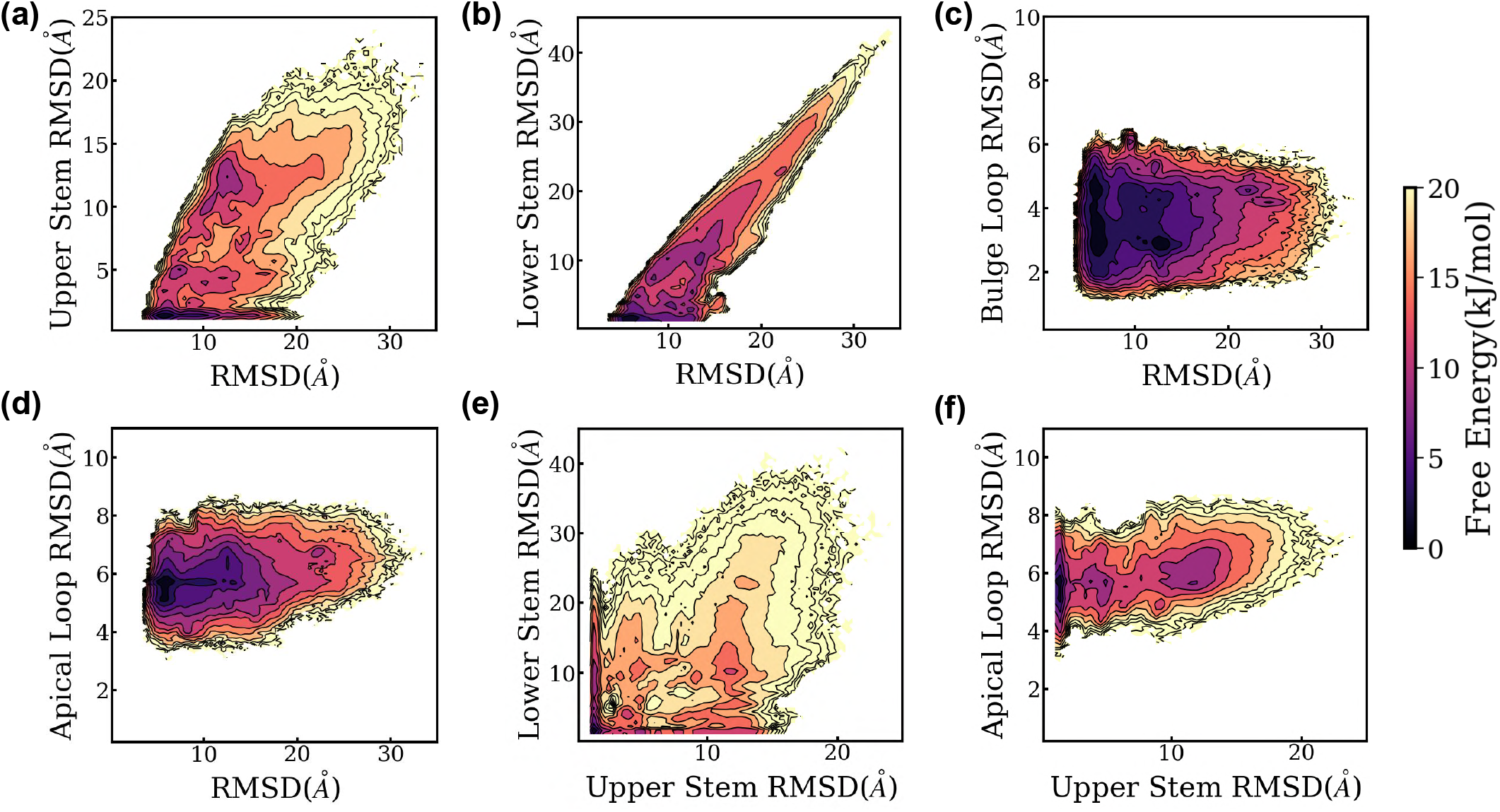
Visualization of the conformational distributions sampled by unbiased MD simulations of the HIV-TAR stem loop at 310 K across different structural order parameters. Heavy-atom RMSDs relative to the NMR structure (PDB ID: 1ANR) are quantified for the entire stem loop and for different individual segments, including the upper stem, lower stem, bulge loop, and apical loop. The resulting free energy surfaces are shown for six pairwise combinations of these RMSD-based coordinates.

**Figure S12:**
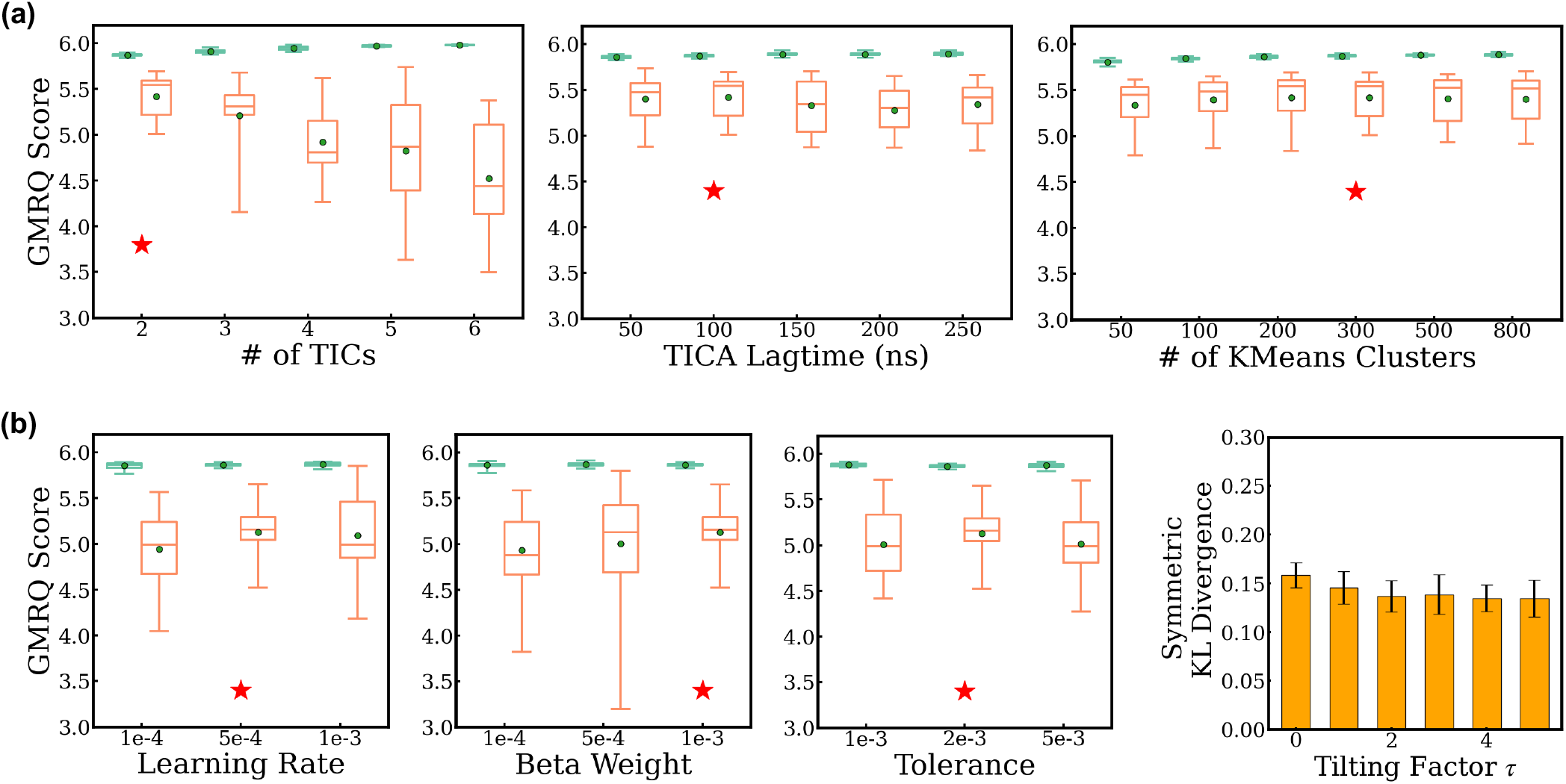
Selection of optimal hyperparameters for training the Latent Thermodynamic Flows model on the HIV-TAR RNA stemloop system at 310K. (a) Choice of hyperparameters for initial labeling with tICA and k-means clustering via cross-validation based on the GMRQ score. (b) Selection of the optimal set of LaTF training hyperparameters. All selection protocols follow the same procedure as that described in Fig. S5.

**Figure S13:**
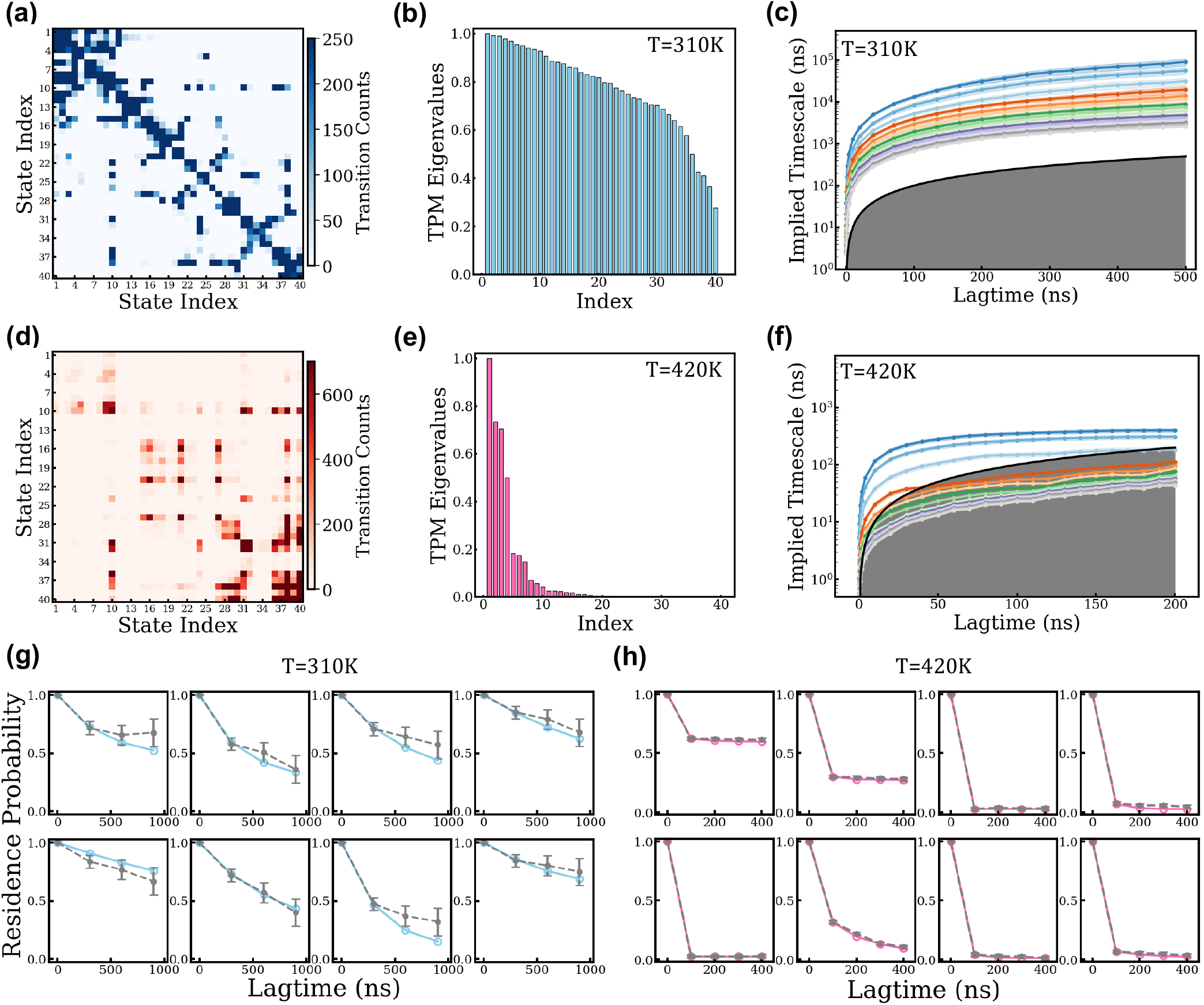
MSMs construction, characterization, and validation for the HIV TAR RNA Stemloop at 310 K and 420K. The LaTF model is trained on MD data collected at 310 K with a lag time of 200 ns. Metastable state definitions are constructed using the trained encoder and decoder, and MD data at 420 K are analyzed using the same state definitions. (a) Raw unsymmetrized TCM directly estimated from simulation data at 310 K with a lag time of 300 ns. (b) Ordered eigenvalues of the TPM at 310 K for a lag time of 300 ns. (c) The fifteen leading implied timescales as a function of transition lag time at 310 K. (d–f) Corresponding analyses for simulations performed at 420 K. The lag times for the TCM and TPM are set to 100 ns. (g) CK test of the MSM built with a lag time of 300 ns for the eight most populated metastable states at 310 K. (h) Corresponding CK test for the MSM constructed with a lag time of 100 ns at 420 K. The gray error bars denote residence probabilities estimated directly from the raw MD simulation data, whereas the blue and red markers represent the corresponding probabilities obtained from MSM propagations. Uncertainties in both the implied timescales and CK tests indicate the standard derivations estimated via bootstrap resampling of the trajectory ensemble fifty times with replacement.

**Figure S14:**
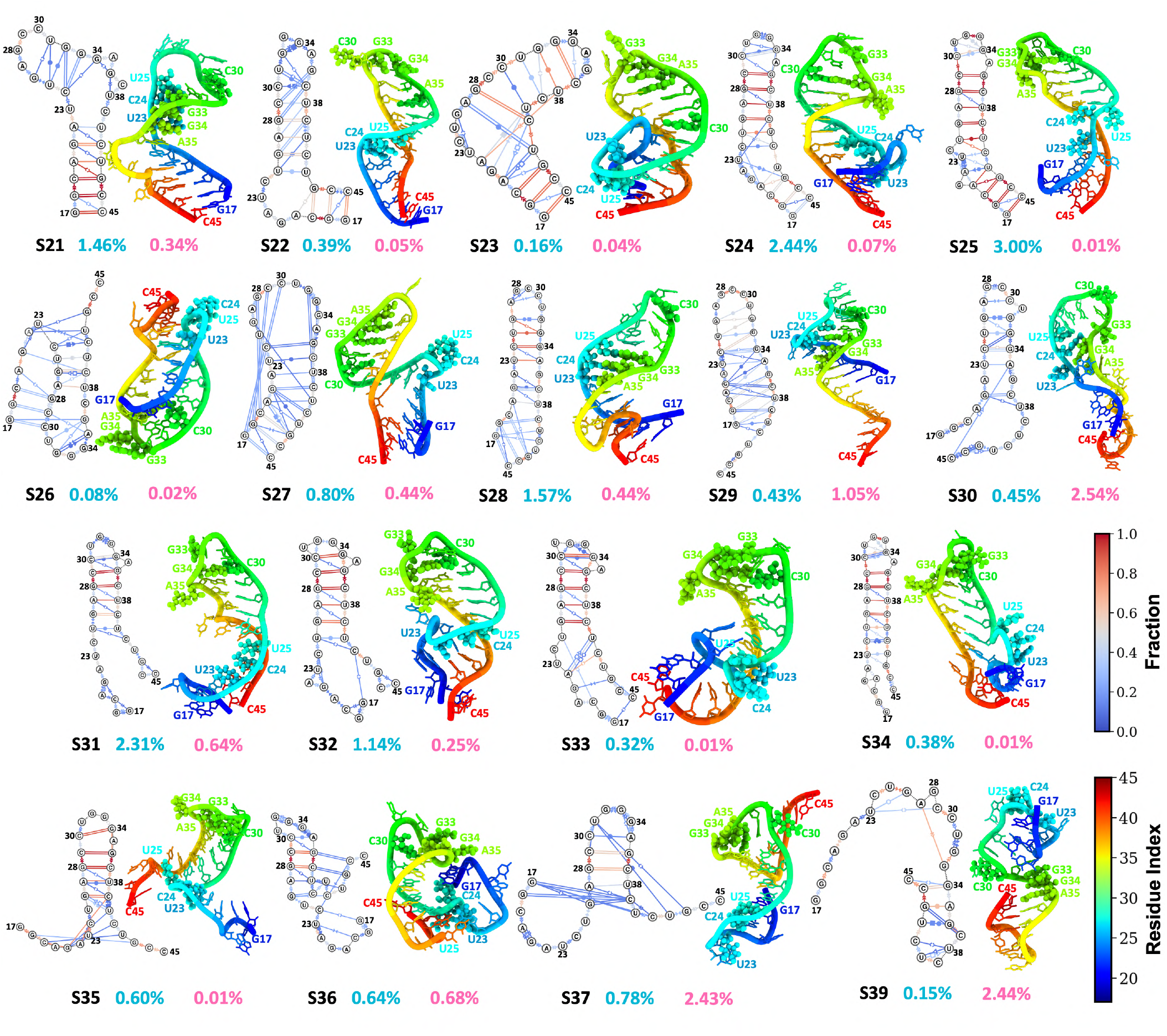
Representative dynamic secondary structures and tertiary structures of the partially folded and unfolded metastable states of the HIV TAR RNA stem-loop. The structural subensembles within different metastable states are visualized as dynamic secondary structures, with distinct interaction patterns colored according to their probabilities, estimated from 500 randomly selected structures sampled from the corresponding states. Structural annotations follow the Leontis–Westhof classification[87]. The representative tertiary structure is chosen as the conformation with the top likelihood ranked by the LaTF normalizing flow model. The apical loop and bulge regions are highlighted in sphere representation. Nucleotide numbering begins at 17 and extends to 45.

**Figure S15:**
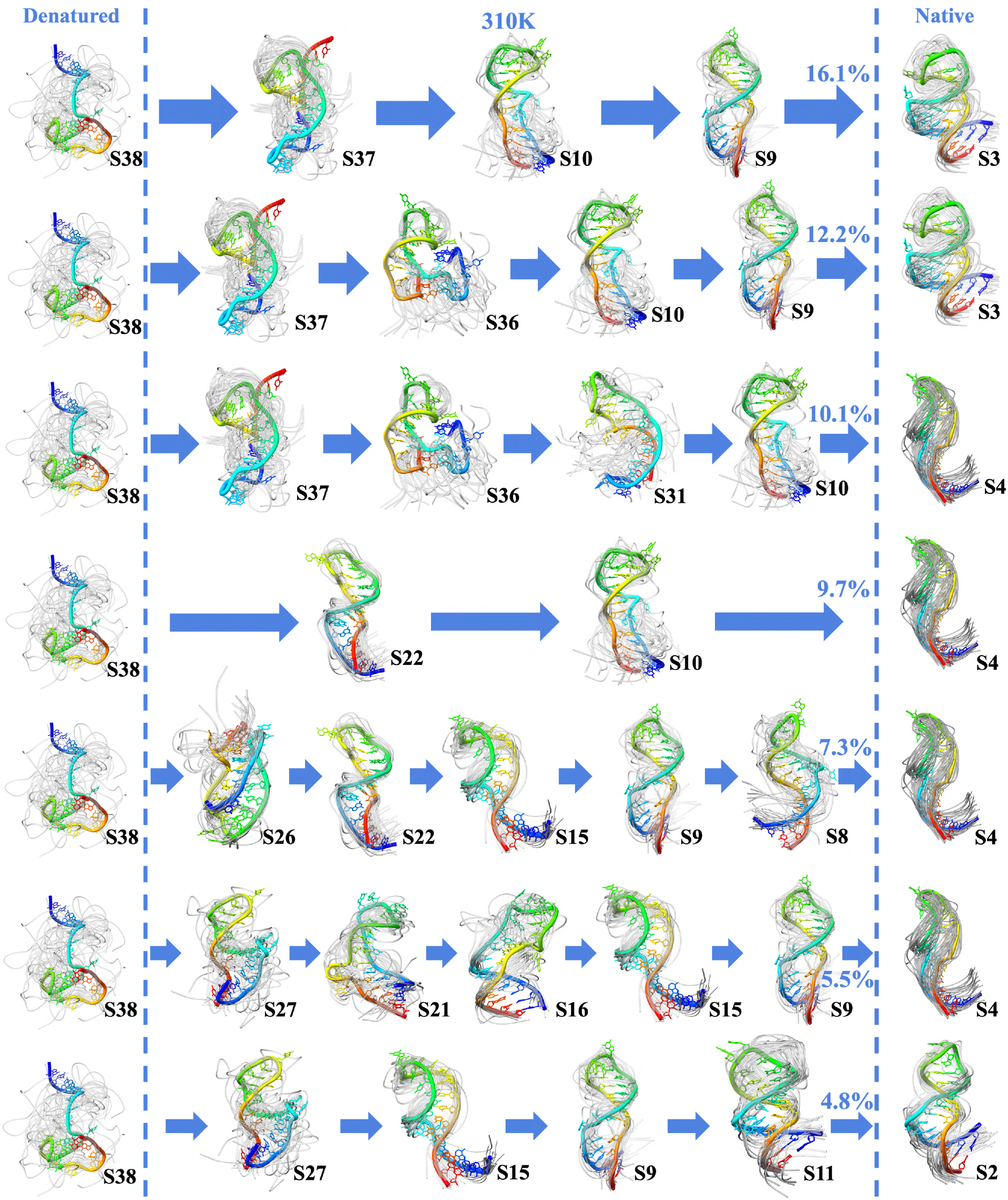
Representative kinetic folding pathways of the HIV TAR stem–loop at 310 K that contribute to the dominant flux. The seven pathways with the highest reactive fluxes are shown. The denatured (unfolded) and native (folded) states are displayed in the leftmost and rightmost columns, respectively, separated by dashed lines. Each state is represented by its 30 highest-likelihood structures as scored by the trained normalizing flow model, with the top-ranked structure highlighted in color and the remaining structures shown in gray. The flux associated with each pathway is indicated, and arrow widths are scaled proportionally to the corresponding flux.

**Figure S16:**
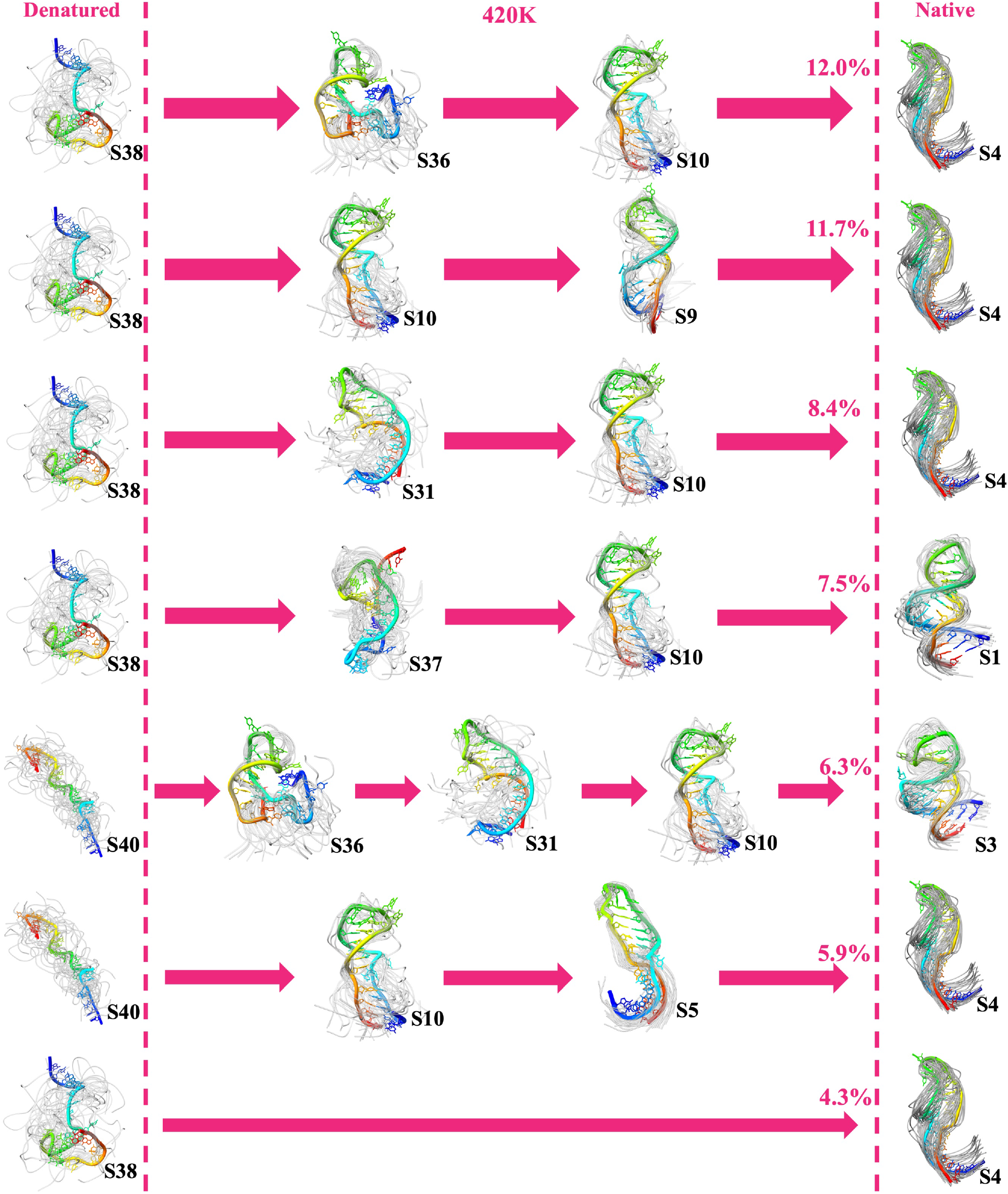
Representative kinetic folding pathways of the HIV TAR stem–loop at 420 K that contribute to the dominant flux. The seven pathways with the highest reactive fluxes are presented in the same manner as in Fig. S16.

**Figure S17:**
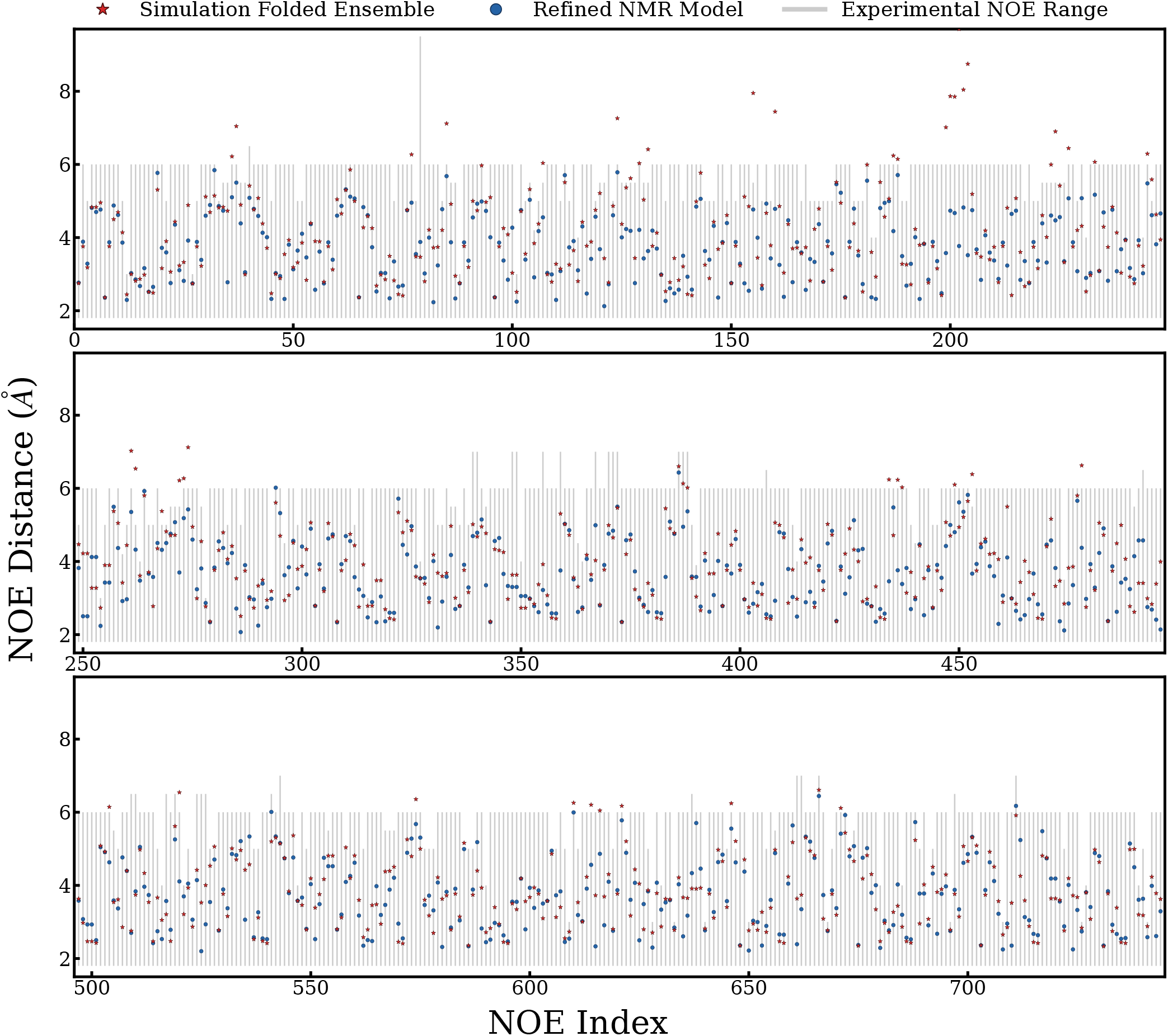
Comparison and validation of the local structural heterogeneity of the HIV TAR stem-loop in the simulation-derived folded-state ensemble against NMR measurements. The ranges of 744 inter-residue proton-proton distances in the HIV TAR stem-loop derived from NOE measurements are shown as gray bars and used as reference values. Five metastable states (S3, S4, S8, S9, and S10) identified by the Latent Thermodynamic Flows model trained on the 310 K simulation data are found to exhibit close structural similarity to the NMR structures and are therefore collectively regarded as the folded ensemble. Ensemble-averaged interproton distances are calculated from conformations sampled across these five states, with sampling weighted according to their normalized MSM state populations; the resulting distances are shown as red stars. For comparison, the corresponding distances calculated from the refined NMR structural models (PDB ID: 1ANR) are shown as blue circles.

**Figure S18:**
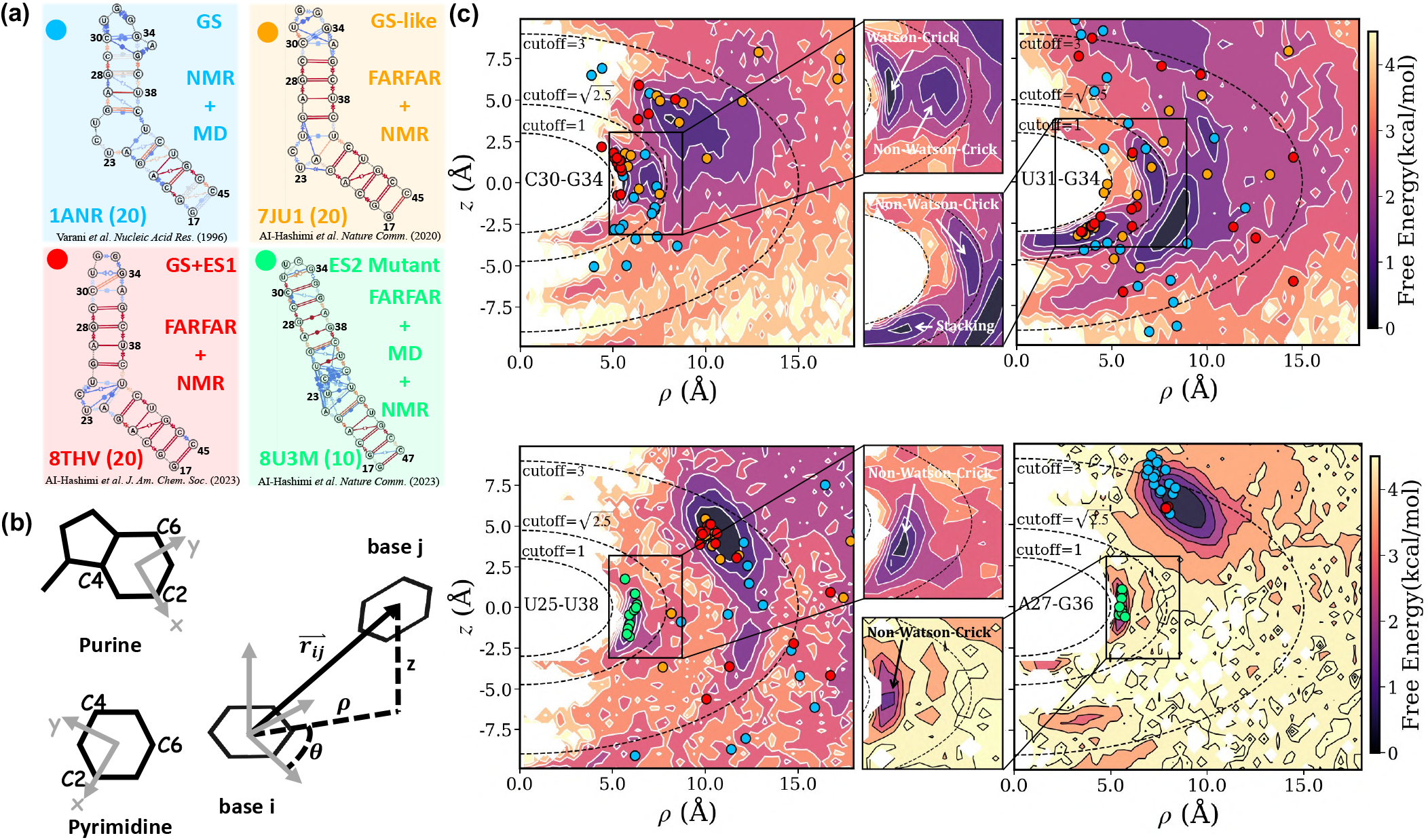
Comparison of experimentally resolved ground- and excited-state structures of the HIV-TAR stem loop with the simulation-derived structural ensemble. (a) Four experimentally derived structural subensembles, with PDB IDs and numbers of structures indicated. The 1ANR ground-state (GS) ensemble (blue) is determined using NOE-derived distance restraints and restrained MD simulations. The 7JU1 (orange; GS-like) and 8THV (red; GS+excited state 1, ES1) ensembles are selected from FARFAR-generated conformational pools to best reproduce NMR residual dipolar coupling (RDC) observables. The 8U3M (green; excited state 2, ES2) ensemble is constructed using a similar strategy, with additional MD sampling incorporated into conformational-pool generation and apical-loop mutations introduced to stabilize the low-populated ES2. (b) Illustration of the local coordinate system used to describe nucleotide-pair geometries. The left panel defines the origin and *x* and *y* axes for purine and pyrimidine bases, while the right panel illustrates the relative position of base *j* within the coordinate frame defined by base *i*. (c) Free-energy landscapes describing the relative positions of the C30(*i*)–G34(*j*), U31(*i*)–G34(*j*), U25(*i*)–U38(*j*), and A27(*i*)–G36(*j*) nucleotide pairs. Landscapes are constructed from the full simulation ensemble and weighted by the corresponding MSM populations. Insets show enlarged regions of the local landscapes, with pairing and stacking interactions annotated. Structural descriptors calculated from the four experimentally derived subensembles are projected onto the corresponding landscapes using the same color scheme as in (a).

**Figure S19:**
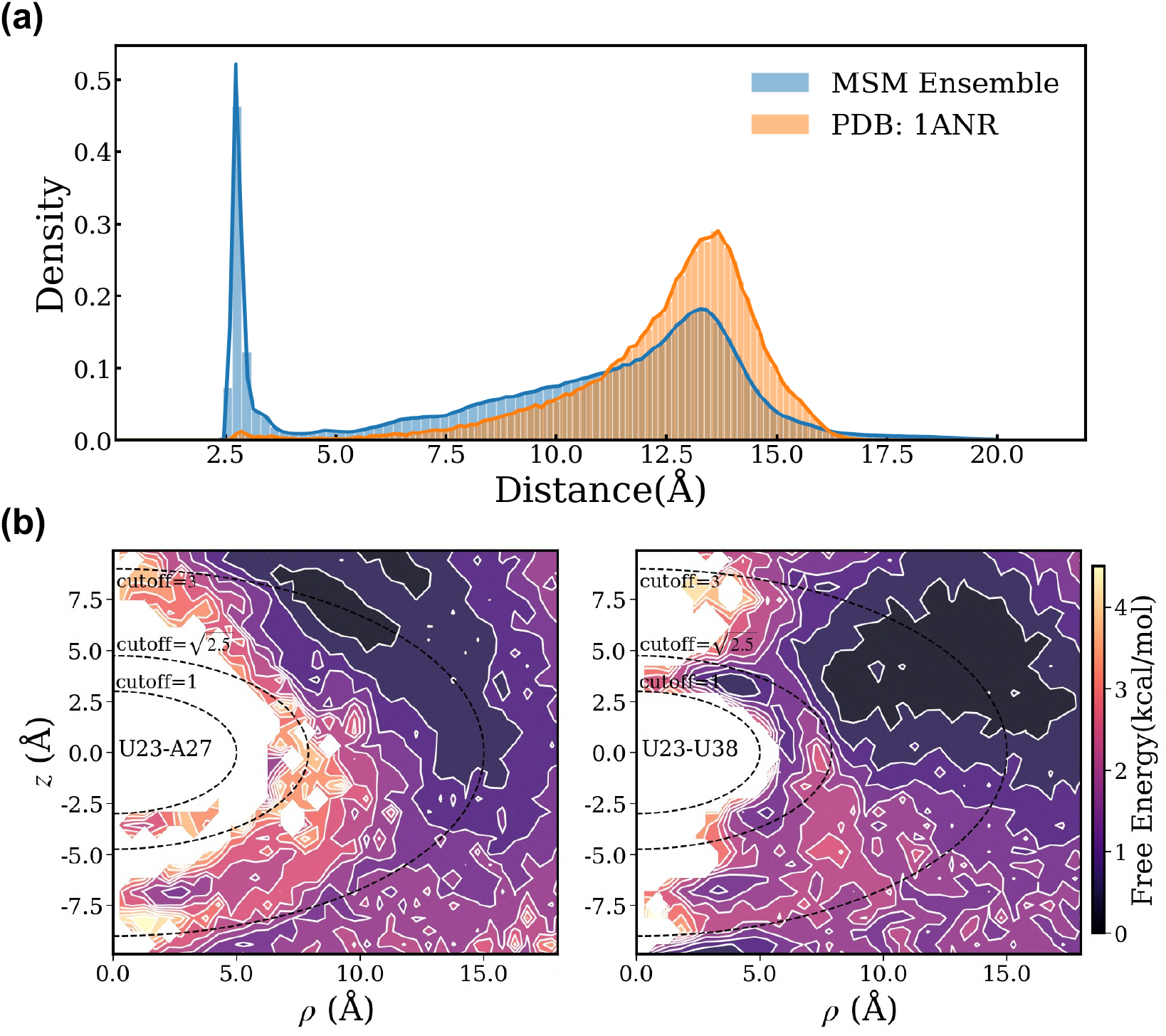
Characterization of interactions between the HIV-TAR bulge and nucleotides in the apical loop and upper stem. (a) Equilibrium distribution of the minimum distance between the bulge (U23–C24–U25) and apical loop (C30–U31-G32-G33-G34-A35), calculated using oxygen and nitrogen atoms only. The simulation-derived distribution is weighted by MSM equilibrium populations, with the corresponding distribution from the 1ANR structures shown for comparison. (b) Distribution of the relative positions of A27 and U38 within a U23-centered local coordinate system across the simulated structural ensemble (see definition in Fig. S18(b)). The corresponding free-energy landscape is also constructed using MSM population weighting.

**Figure S20:**
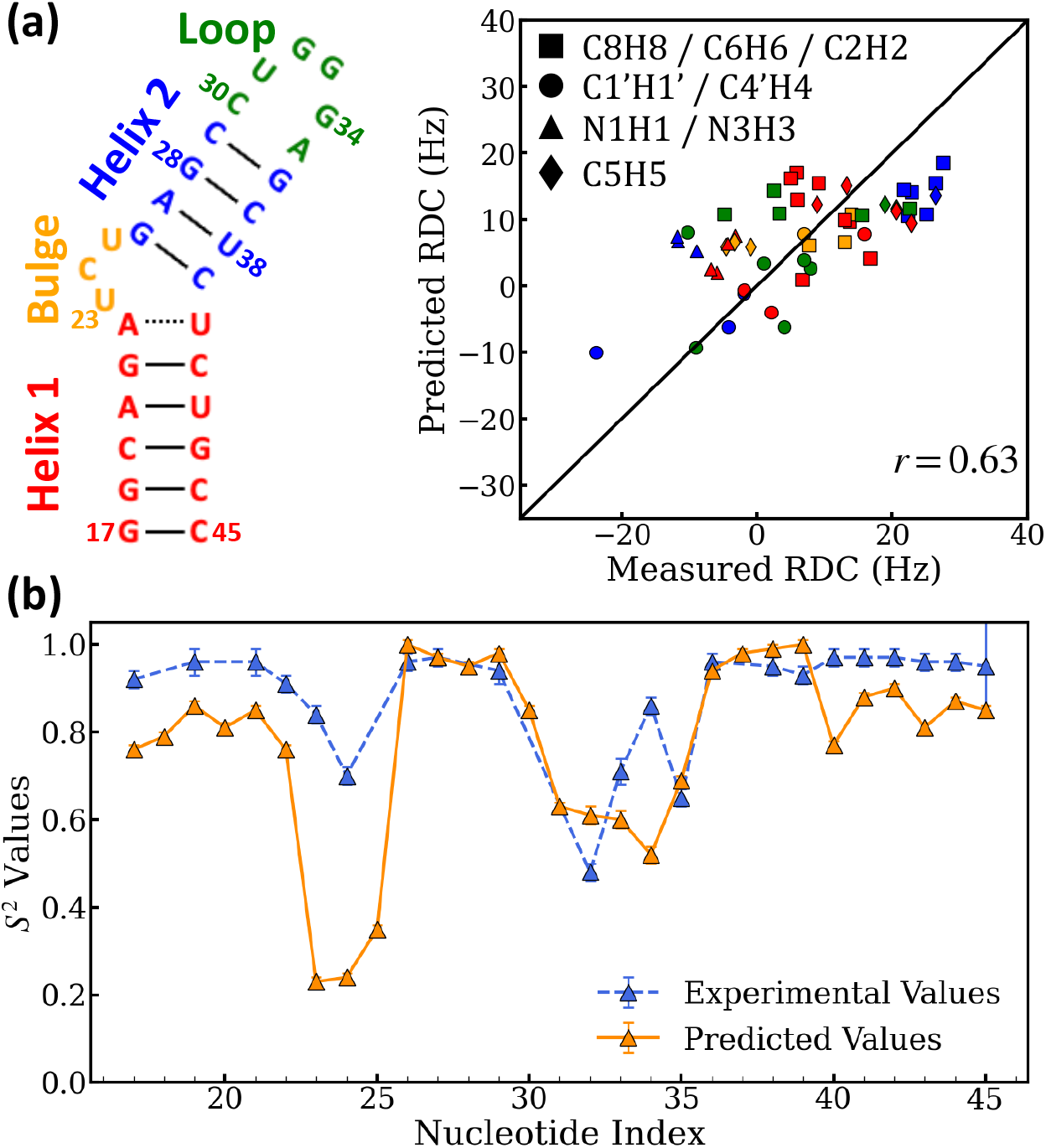
Comparison of NMR measurements for HIV TAR RNA with corresponding computational predictions derived from MD simulations and MSMs. (a) (Left) Secondary structure representation of the TAR ground state, with different structural motifs color-coded for clarity. (Right) Correlation between experimental and computationally predicted residual dipolar couplings (RDCs) values for various bond vectors (of directly bonded nuclei). Predicted values are derived from the MD-sampled structural ensemble and reweighted according to MSM-derived state populations. Data points are categorized and visualized by bond type (markers) and structural motif (color). The Pearson correlation coefficient *r* is reported. (b) Comparison between experimental (blue) and computationally predicted (orange) Lipari-Szabo generalized order parameter *S*^2^ of different bond vectors across the RNA sequence. For purine residues (adenine and guanine), *S*^2^ is determined for the C8–H8 dipolar interaction, while for pyrimidine residues (cytosine and uracil), values correspond to the C6–H6 dipolar interaction. All calculations are conducted only using the simulation data and MSM generated at 310K. Comprehensive details regarding the quantification of RDC values and *S*^2^ order parameter are provided in the text of Supplementary Information.

**Figure S21:**
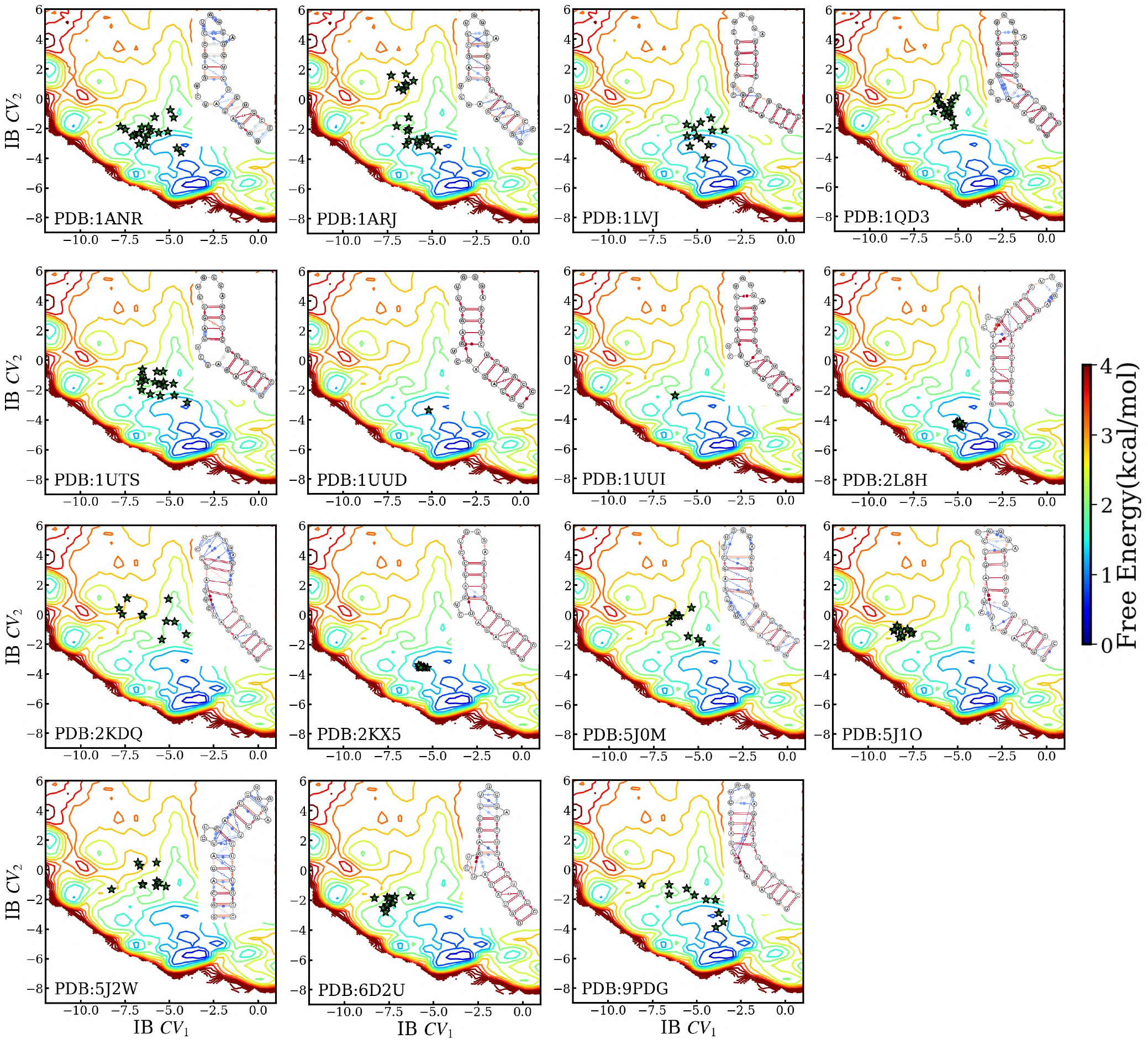
Projection of experimentally resolved HIV TAR stem–loop structures from the Protein Data Bank (PDB) onto the latent information bottleneck (IB) collective variable (CV) space using the trained latent thermodynamic flow (LaTF) model. Over the past decades, structures of *apo* HIV TAR RNA and its complexes with various binders, including small molecules and peptides, have been determined by solution nuclear magnetic resonance and deposited in the PDB. These structures are encoded using r-vectors and pairwise distances between 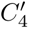 atoms, and subsequently projected through the trained encoder of the LaTF model. Each structure is represented as a star marker, labeled by its corresponding PDB ID. The background depicts the local free energy landscape derived from MSM-reweighted simulation data. Insets show the corresponding dynamic secondary structure ensembles constructed for each PDB entry.

**Figure S22:**
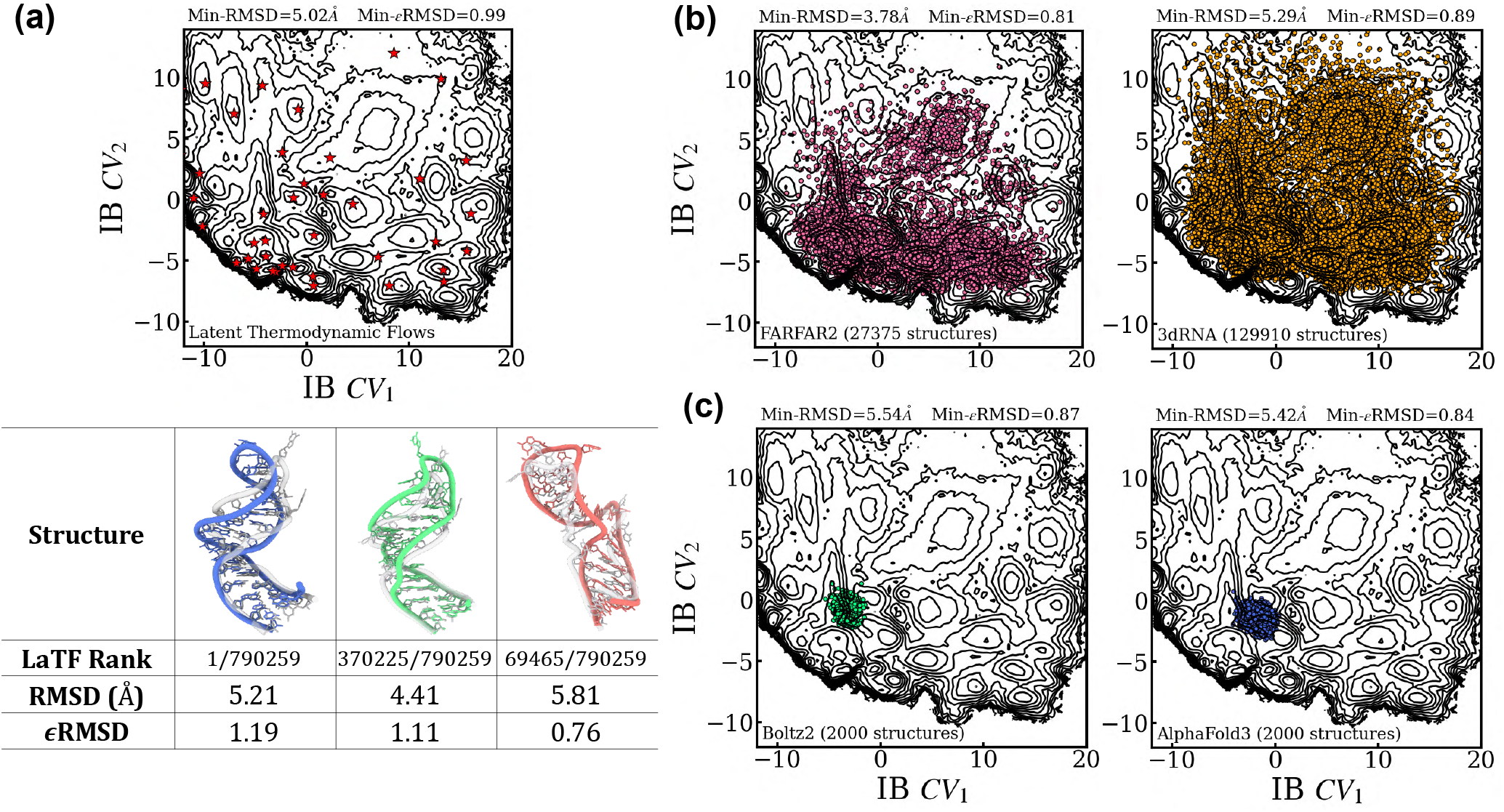
Performance assessment of tertiary structure prediction and modeling approaches for sampling the conformational heterogeneity of the HIV-TAR stemloop RNA. (a) The molecular dynamics conformations sampled at 310 K are projected onto the 2D latent information bottleneck (IB) space using the trained LaTF encoder. The corresponding free energy landscape is constructed from MSM-reweighted conformational ensemble. For each metastable state identified by the LaTF model, the most probable structure, as determined by the LaTF normalizing flow module, is indicated by a red star. Three representative structures from the dominant states (States 1–10) are selected for further analysis: (i) the top-ranked structure according to the LaTF likelihood, (ii) the structure with the smallest RMSD, and (iii) the structure with the smallest *ɛ*RMSD, both calculated with respect to the NMR structure (PDB ID: 1ANR). The corresponding structures, together with their ranks, RMSD values, and *ɛ*RMSD values, are summarized in the table. MD sampled conformations are shown in color, whereas the NMR structure is rendered transparently for comparison. (b) Assessment of thermodynamics-based fragment assembly methods for sampling the conformational landscape of the HIV-TAR stemloop RNA. FARFAR2 and 3dRNA are applied to generate tertiary structure ensembles from 365 candidate secondary structures. Conformational heterogeneity is sampled through Monte Carlo simulations across different temperatures guided by the knowledge-based scoring functions implemented in each method. The resulting structures are featurized using r-vector and pairwise distance descriptors and embedded into the latent IB space using the trained LaTF encoder. The minimum RMSD and *ɛ*RMSD values across all sampled structures, computed relative to the NMR structure, are reported. (c) Two deep learning–based structure prediction methods, Boltz2 and AlphaFold3, are employed to generate tertiary structures of HIV-TAR directly from its nucleotide sequence. Structural diversity is explored by varying the random seeds and diffusion sampling parameters of the generative models. For each method, 2,000 structures are generated and projected into the latent IB space using the trained LaTF encoder. The minimum RMSD and *ɛ*RMSD values across all generated 2,000 structures with reference of the NMR structure are also reported.

**Figure S23:**
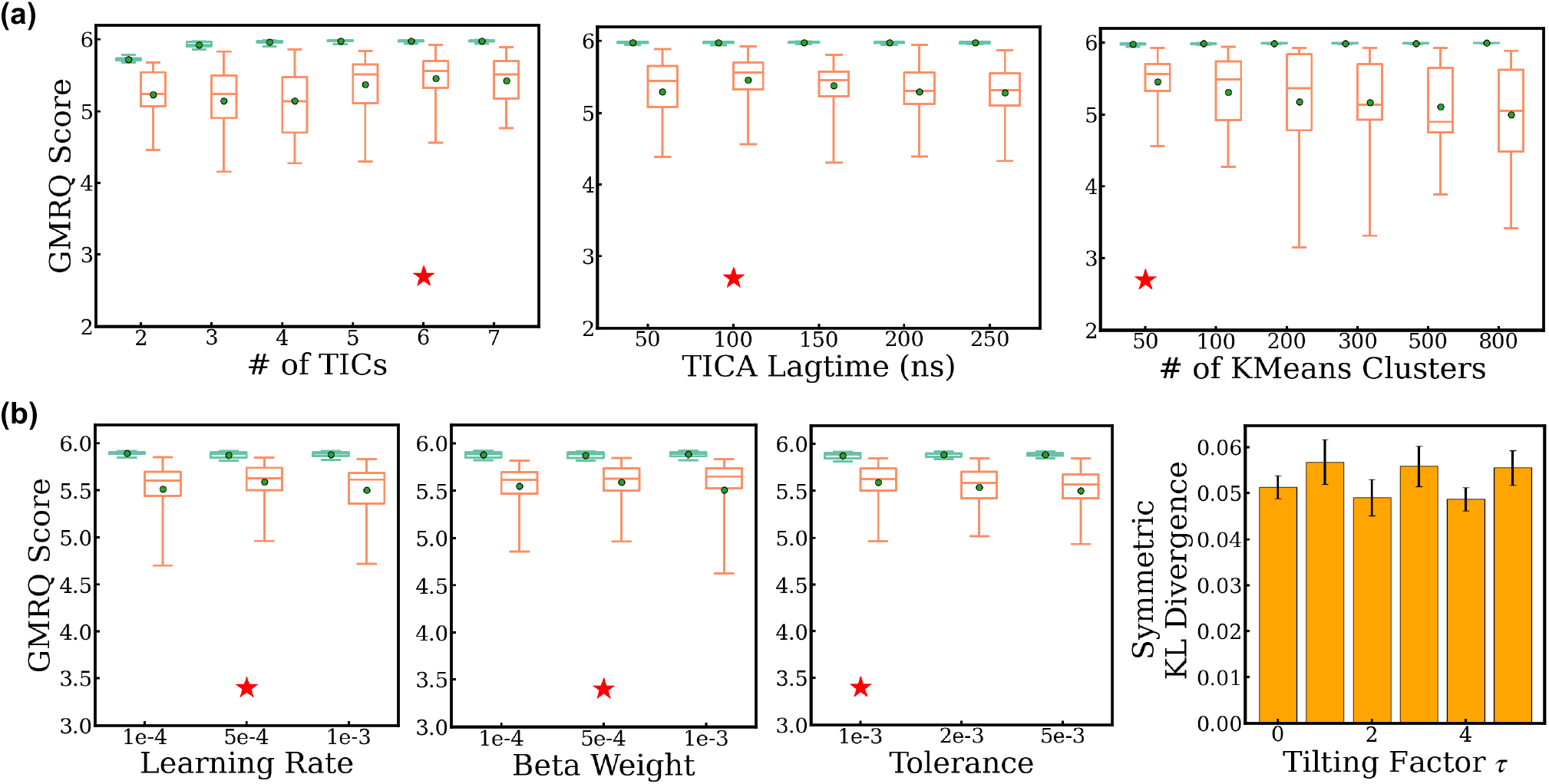
Selection of optimal hyperparameters for training the Latent Thermodynamic Flows model on the MicroROSE RNA Thermometer system at 310K. (a) Choice of hyperparameters for initial labeling with tICA and k-means clustering via cross-validation based on the GMRQ score. (b) Selection of the optimal set of LaTF training hyperparameters. All selection protocols follow the procedure described in Fig. S5.

**Figure S24:**
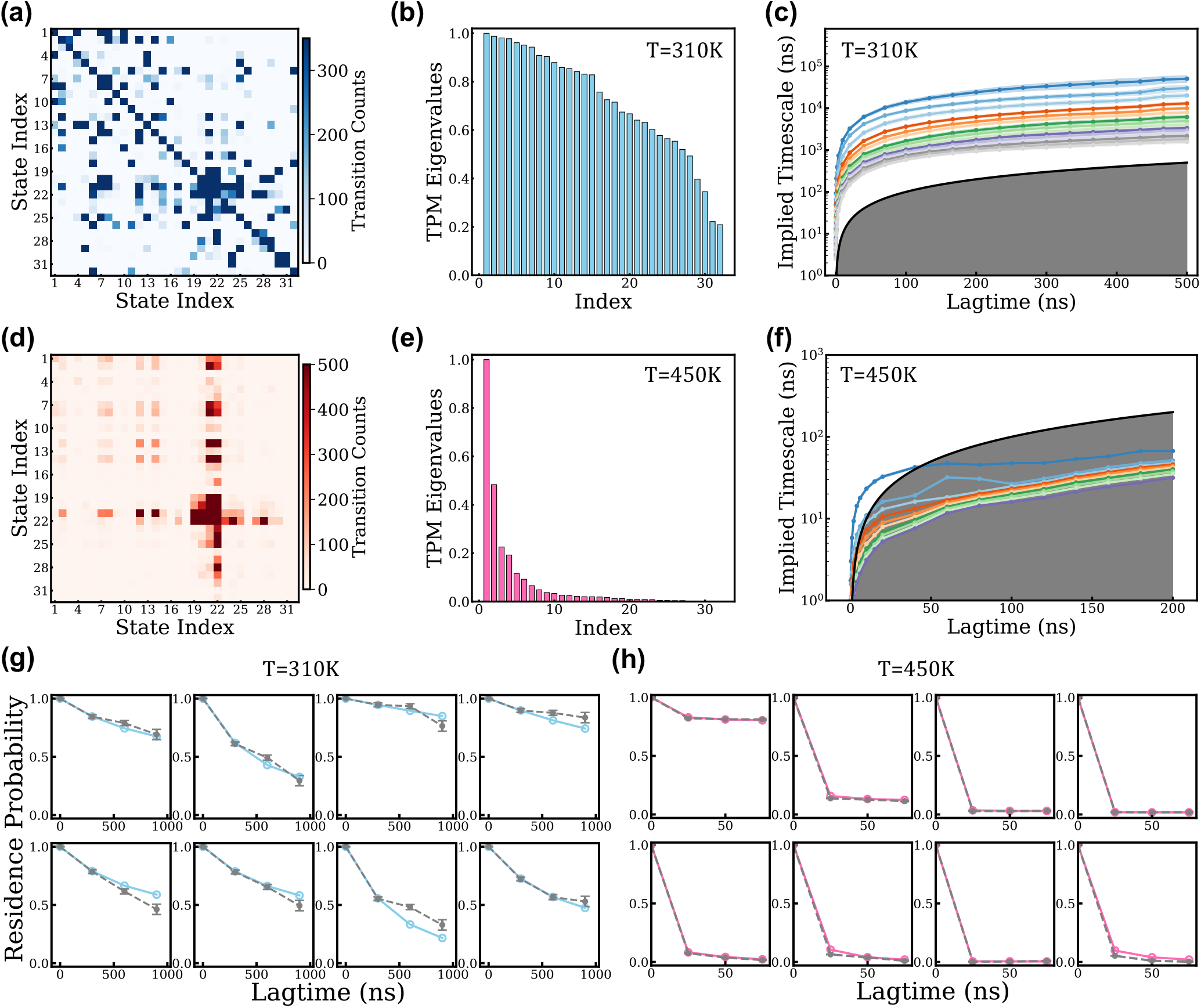
MSMs construction, characterization, and validation for the MicroROSE RNA Thermometer at 310 K and 450K. The LaTF model is trained on MD data collected at 310 K with a lag time of 50 ns. Metastable state definitions are constructed using the trained encoder and decoder, and MD data at 450 K is analyzed using the same state definitions. (a) Raw unsymmetrized TCM directly estimated from simulation data at 310 K with a lag time of 300 ns. (b) Ordered eigenvalues of the TPM at 310 K for a lag time of 300 ns. (c) The fifteen leading implied timescales as a function of transition lag time at 310 K. (d–f) Corresponding analyses for simulations performed at 450 K. The lag times for the TCM and TPM are set to 25 ns. (g) CK test of the MSM built with a lag time of 300 ns for the eight most populated metastable states at 310 K. (h) Corresponding CK test for the MSM constructed with a lag time of 25 ns for the eight most populated metastable states at 450 K. The gray error bars denote residence probabilities estimated directly from the raw MD simulation data, whereas the blue and red markers represent the corresponding probabilities obtained from MSM propagation. Uncertainties in both the implied timescales and CK tests indicate the standard derivations estimated via bootstrap resampling of the trajectory ensemble fifty times with replacement.

**Figure S25:**
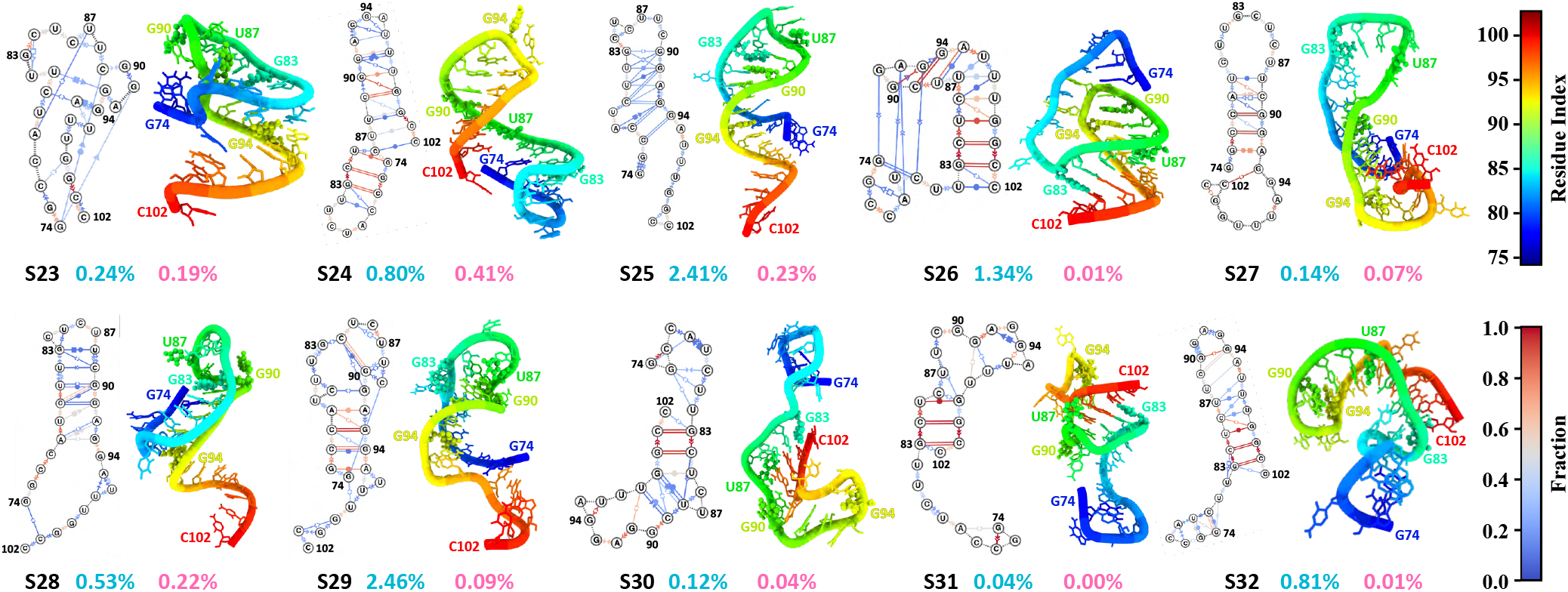
Representative dynamic secondary structures and tertiary structures of misfolded intermediates of the MicroROSE RNA Thermometer. The structural sub-ensembles within different metastable states are visualized as dynamic secondary structures with distinct interaction patterns colored according to their probabilities. The interaction patterns and probabilities are calculated over 700 randomly sampled structures from each state. Structural annotations follow the Leontis–Westhof classification[87]. The representative tertiary structure is chosen as the conformation with the top likelihood ranked by the LaTF normalizing flow model. G83 and G94 terminate the internal loop and are shown in sphere representation. Apical loop nucleotides U87 and U90 are also highlighted in sphere representation. Nucleotide numbering begins at 74 and extends to 102.

**Figure S26:**
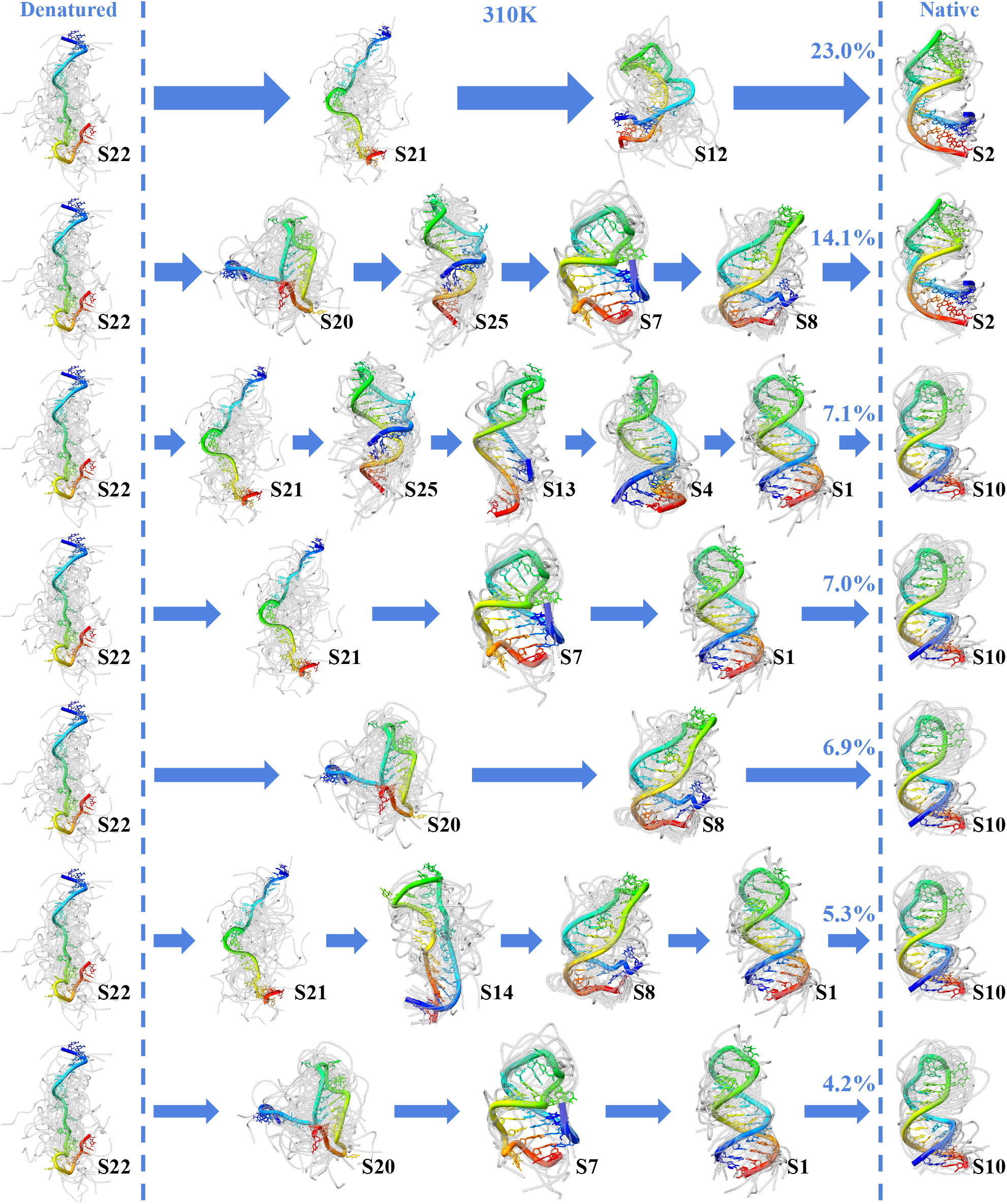
Representative kinetic folding pathways of the MicroROSE RNA Thermometer at 310 K that contribute to the dominant flux. The seven pathways with the highest reactive fluxes are shown. The denatured (unfolded) and native (folded) states are displayed in the leftmost and rightmost columns, respectively, separated by dashed lines. Each state is represented by its 30 highest-likelihood structures as scored by the LaTF normalizing flow model, with the top-ranked structure highlighted in color and the remaining structures shown in gray. The flux associated with each pathway is indicated, and arrow widths are scaled proportionally to the corresponding flux.

**Figure S27:**
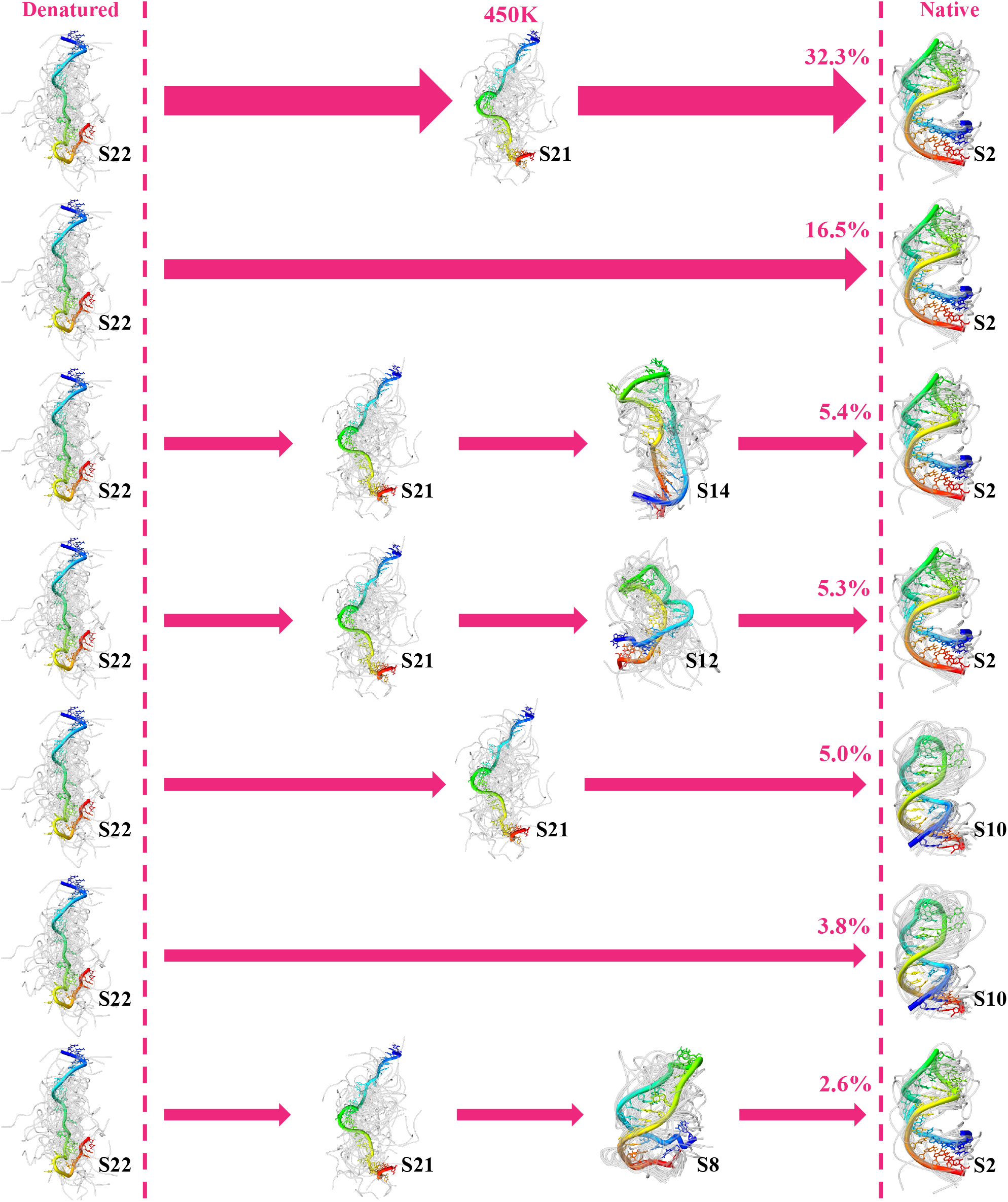
Representative kinetic folding pathways of the MicroROSE RNA Thermometer at 450 K that contribute to the dominant flux. The seven pathways with the highest reactive fluxes are presented in the same manner as Fig. S26.

**Figure S28:**
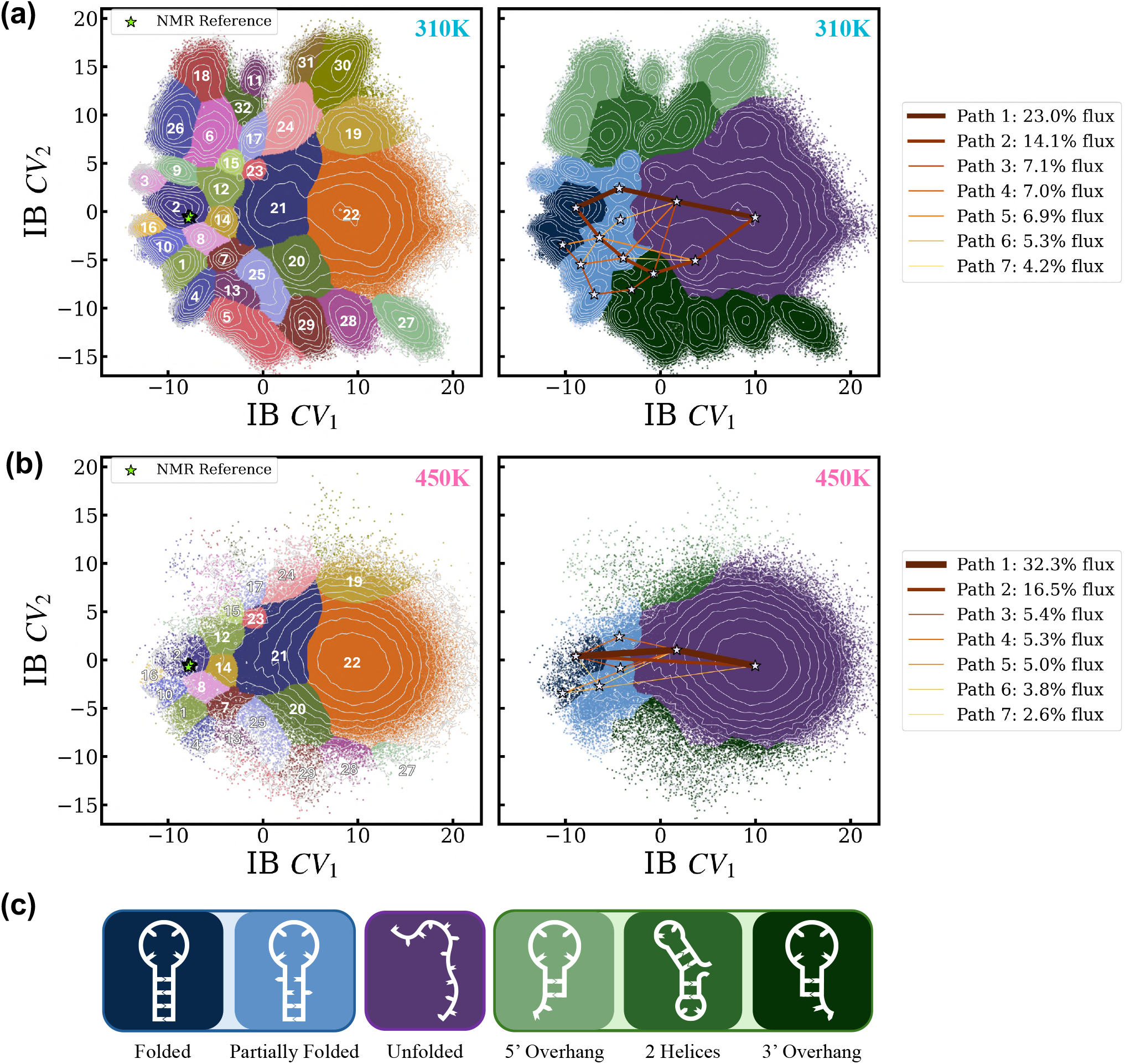
Coarse-graining of the microROSE conformational space into structurally distinct regions. (a) State labels, coarse-grained free-energy surface and top transition paths at 310K. (b) State labels, coarse-grained free-energy surface and top transition paths at 450K. (c) Legend for right panels: dark blue - folded, light blue - partially folded, purple - unfolded, light green - misfolded: 5’ overhang, medium green - misfolded: two stacked helices, dark green - misfolded: 3’ overhang. The left panels depict free-energy landscapes colored by predicted state labels obtained from the LaTF encoder-decoder. The reference PDB structures refined from NMR data are mapped into IB space by the LaTF encoder and shown as chartreuse stars. The right panels contain a coarse-grained description of the folding landscape, obtained by grouping structurally similar states into categories shown in (c). The transition paths shown in S26 and S27 are visualized over this coarse-grained coloring.

**Figure S29:**
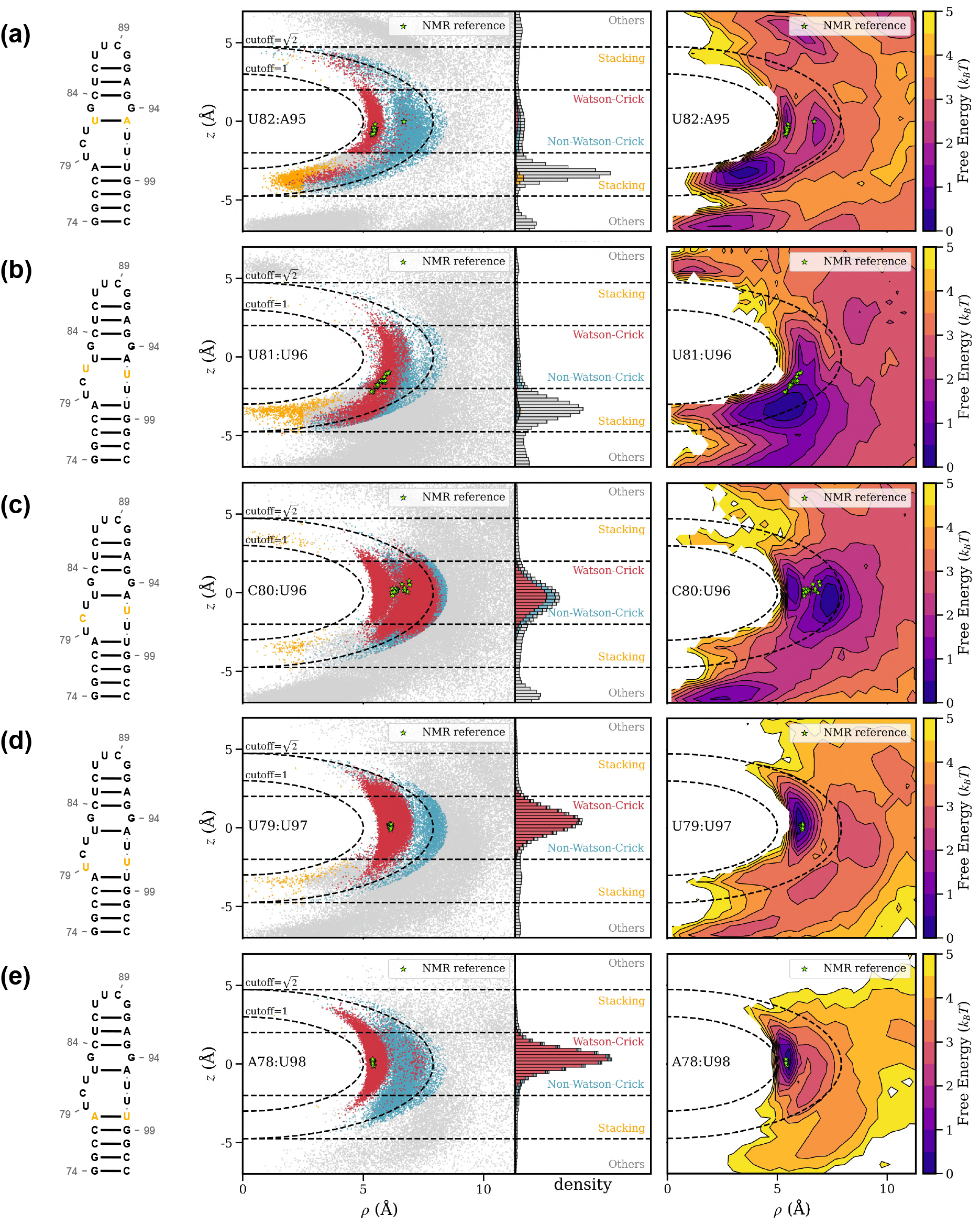
Illustration of the equilibrium interactions associated with RNAT internal loop nucleotide pairs at 310K. Five nucleotide interactions pairs are shown: (a) U82-A95, (b) U81-U96, (c) C80-U96, (d) U79-U97, (e) A78-U98. For each pair, the left column shows the consensus secondary structure taken from a *Barnaba* [63] annotation of PDB 2GIO with the specific nucleotides highlighted in orange. These visualizations were created using RNACanvas [88]. The center column shows a *ρ*-*z* plot of the interactions, colored by type, between each nucleotide pair sampled over the folded sub-ensemble (S2, S8, S10). The right column is a free-energy surface created from the (*ρ*-*z*) plot shown in the center column. All plots are re-weighted by the metastable state populations given by the Markov state model, as described in S8.

**Figure S30:**
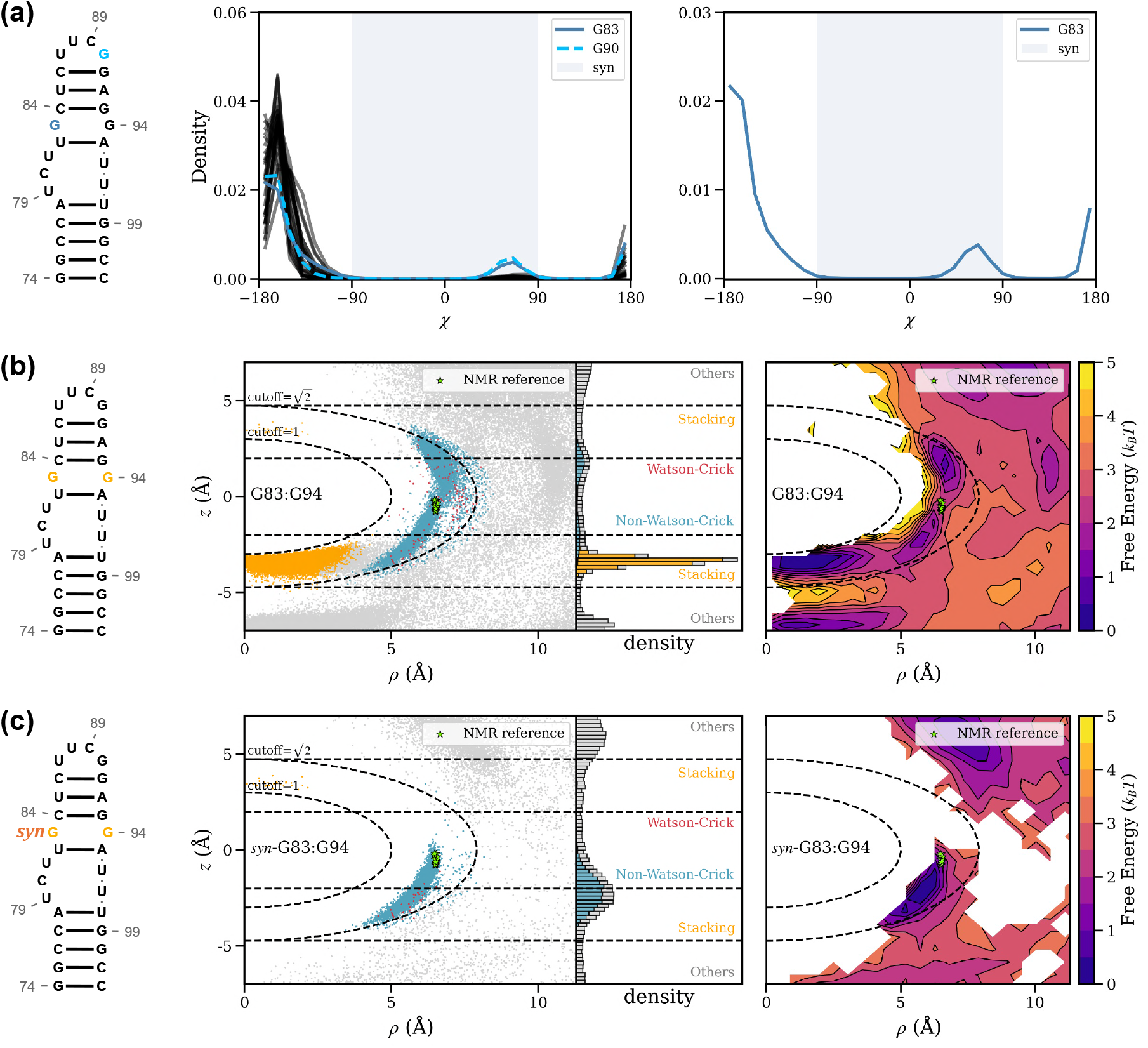
Illustration of the equilibrium interactions associated with the RNAT G83-G94 pair at 310K. (a) Left: PDB 2GIO consensus secondary structure with G83 and G90 highlighted. Center: *χ* torsion angle distribution for all nucleotides in the microROSE RNA Thermometer. G83 and G90 are highlighted as the only two nucleotides with a significant population in the *syn* conformation. Right: *χ* torsion angle distribution for G83. (b) Left: PDB 2GIO consensus secondary structure with G83 and G94 highlighted. Center & Right: G83-G94 interaction *ρ*-*z* scatterplot and free-energy surface for all structures in the folded sub-ensemble. (c) Left: PDB 2GIO consensus secondary structure with G83 and G94 highlighted. *syn* is added to note that this depicts the set of folded structures for which G83 is in the *syn* conformation. Center & Right: G83-G94 interaction *ρ*-*z* scatterplot and free-energy surface computed for only structures in the folded sub-ensemble where G83 is in a *syn* conformation. The *ρ*-*z* plots and consensus secondary structures were constructed as described in S29.

**Figure S31:**
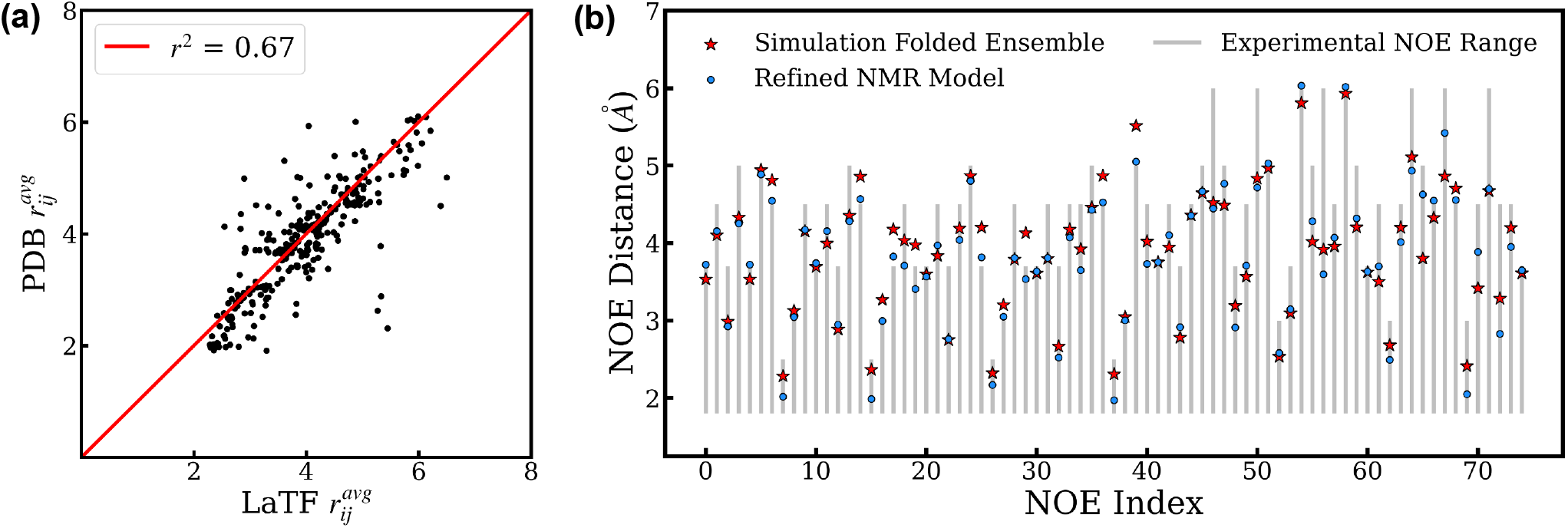
Comparison and validation of the structural heterogeneity of the MicroROSE RNAT Thermometer in the simulation-derived folded state against NMR measurements. (a) A Latent Thermodynamic Flows model trained on simulation data at 310 K identifies a dominant metastable state (state 2) that exhibits a significantly smaller structural deviation from the native NMR structure than all other states. Five hundred structures are randomly sampled from this state to represent the folded ensemble. The average inter-hydrogen distances quantified from these conformers are plotted against the corresponding distances calculated from the refined NMR structural models (PDB ID: 2GIO). n=338 as this calculation is performed for the 338 proton pairs specified by the reported NOE restraints. (b) The ranges of the first 75 inter-proton distances derived from nuclear Overhauser effect (NOE) measurements are shown as gray bars and used as reference values. Inter-proton distances averaged over the LaTF state 2 ensemble are shown as red stars. The corresponding inter-proton distances averaged over the NMR ensemble are shown as blue circles. We visualize only the first 75 distances for clarity but compute other statistics over the full set of 338 restraints.

**Figure S32:**
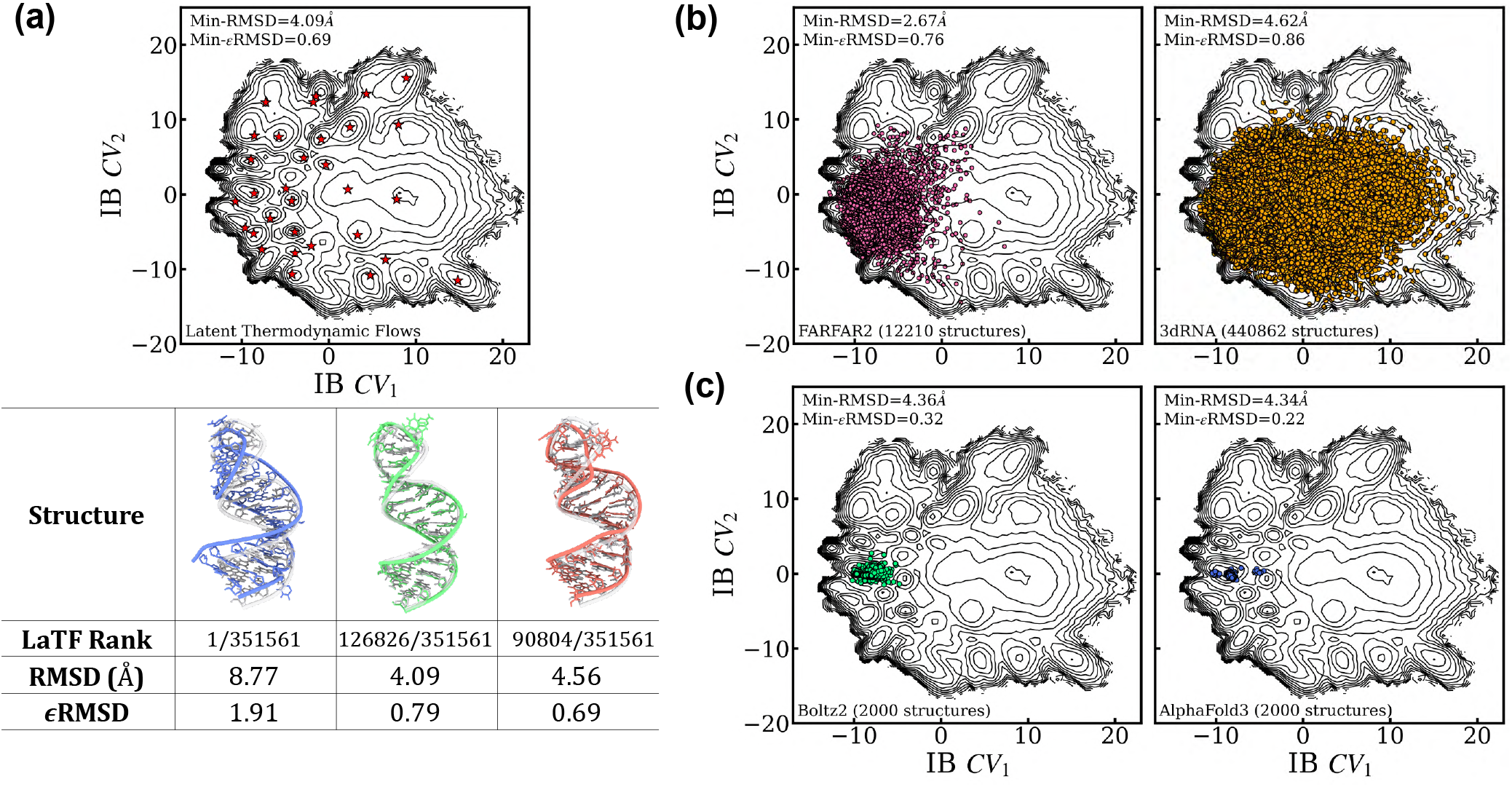
Performance assessment of tertiary structure prediction and modeling approaches for sampling the conformational heterogeneity of the MicroROSE RNA Thermometer (RNAT). (a) The molecular dynamics conformations sampled at 310 K are projected onto the 2D latent information bottleneck (IB) space using the trained LaTF encoder. The corresponding free energy landscape is constructed from MSM-reweighted conformational ensemble. For each metastable state identified by the LaTF model, the most probable structure, as determined by the LaTF normalizing flow module, is indicated by a red star. Three representative structures from the well-populated states (States 1–20) are selected for further analysis: (i) the top-ranked structure according to the LaTF likelihood, (ii) the structure with the smallest RMSD, and (iii) the structure with the smallest *ɛ*RMSD, both calculated with respect to the NMR structure (PDB ID: 2GIO). The corresponding structures, together with their ranks, RMSD values, and *ɛ*RMSD values, are summarized in the table. MD sampled conformations are shown in color, whereas the NMR structure is rendered transparently for comparison. (b) Assessment of thermodynamics-based fragment assembly methods for sampling the conformational landscape of the MicroROSE RNAT. FARFAR2 and 3dRNA are applied to generate tertiary structure ensembles from 1221 candidate secondary structures. Conformational heterogeneity is sampled through Monte Carlo simulations across different temperatures guided by the knowledge-based scoring functions implemented in each method. The resulting structures are featurized using *r* - vector and pairwise distance descriptors and embedded into the latent IB space using the trained LaTF encoder. The minimum RMSD and *ɛ*RMSD values across all sampled structures, computed relative to the NMR structure, are reported. (c) Two deep learning–based structure prediction methods, Boltz2 and AlphaFold3, are employed to generate tertiary structures of the MicroROSE RNAT directly from its nucleotide sequence. Structural diversity is explored by varying the random seeds and diffusion sampling parameters of the generative models. For each method, 2,000 structures are generated and projected into the latent IB space using the trained LaTF encoder. The minimum RMSD and *ɛ*RMSD values across all generated 2,000 structures with reference of the NMR structure are also reported.

